# Analysis of small RNA silencing in *Zymoseptoria tritici* – wheat interactions

**DOI:** 10.1101/501650

**Authors:** Graeme J. Kettles, Bernhard J. Hofinger, Pingsha Hu, Carlos Bayon, Jason J. Rudd, Dirk Balmer, Mikael Courbot, Kim E. Hammond-Kosack, Gabriel Scalliet, Kostya Kanyuka

## Abstract

Cross-kingdom small RNA (sRNA) silencing has recently emerged as a mechanism facilitating fungal colonization and disease development. Here we characterized RNAi pathways in *Zymoseptoria tritici*, a major fungal pathogen of wheat, and assessed their contribution to pathogenesis. Computational analysis of fungal sRNA and host mRNA sequencing datasets was used to define the global sRNA populations in *Z. tritici* and predict their mRNA targets in wheat. 389 *in planta*-induced sRNA loci were identified. sRNAs generated from some of these loci were predicted to target wheat mRNAs including those potentially involved in pathogen defense. However, molecular approaches failed to validate targeting of selected wheat mRNAs by fungal sRNAs. Mutant strains of *Z. tritici* carrying deletions of genes encoding key components of RNAi such as Dicer-like (DCL) and Argounate (AGO) proteins were generated, and virulence bioassays suggested that these are dispensable for full infection of wheat. Nonetheless, our results did suggest the existence of non-canonical DCL-independent pathway(s) for sRNA biogenesis in *Z. tritici*. dsRNA targeting essential fungal genes applied *in vitro* or generated from an RNA virus vector *in planta* in a procedure known as HIGS (Host-Induced Gene Silencing) was ineffective in preventing *Z. tritici* growth or disease. We also demonstrated that *Z. tritici* is incapable of dsRNA uptake. Collectively, our data suggest that RNAi approaches for gene function analyses in this fungal species and potentially also as a control measure may not be as effective as has been demonstrated for some other plant pathogenic fungi.

## 1 Introduction

In plants, RNA silencing also known as RNAi (RNA interference) describes a collection of related biochemical pathways where numerous small RNA (sRNA) species, typically 21-24 nucleotides (nt) in length e.g. miRNA (microRNA), siRNA (small interfering RNA), tasiRNA (trans-acting siRNA), natsiRNA (natural antisense siRNA) modify the expression of target mRNA molecules through classical Watson-Crick base-pairing (Ghildiyal & Zamore, 2009; Vickers *et al.*, 2015). Generation of sRNA is achieved through the action of a Dicer-like (DCL) RNA helicase with an RNase III activity acting on double-stranded RNA (dsRNA) molecules synthesized by an RNA-dependent RNA polymerase (RdRp) or single-stranded RNAs folded into a secondary hairpin-like structure (Xie *et al.*, 2004; Gasciolli *et al.*, 2005). Once produced, sRNAs exert their activity on targets bearing sufficiently complementarity through the activity of an RNA-induced Silencing Complex (RISC) containing an Argonaute (AGO) class endonuclease, which binds sRNAs, in its core (Vazquez *et al.*, 2010). Target mRNAs are typically negatively-regulated either through cleavage or translational inhibition (Vazquez *et al.*, 2010). RNAi pathways are deeply integrated in plant immune processes and are involved in defense responses against viral, bacterial and fungal pathogens, and invertebrate pests (Hamilton & Baulcombe, 1999; Navarro *et al.*, 2006; Pandey & Baldwin, 2007; Ellendorff *et* a*l.*, 2009; Kettles *et al.*, 2013). Additionally, they orchestrate numerous developmental processes, responses to changes in the abiotic environment, and DNA methylation and heterochromatin formation (Fujii *et al.*, 2005; Sunkar *et al.*, 2007; Matzke *et al.*, 2007). In filamentous fungi, a diverse range of equivalent RNAi pathways have broadly similar requirement for DCL, AGO and RdRp as in plants. These mechanisms are involved in protection against invasive nucleic acids e.g. those generated from retrotransposons (Nolan *et al.*, 2005; Segers *et al.*, 2007), control of heterochromatin formation (Volpe *et al.*, 2002), and in the meiotic silencing by unpaired DNA pathway (Chang *et al.*, 2012). Intriguingly, RNAi pathways have also been significantly modified or lost in some fungal lineages thus suggesting that in some circumstances loss of RNAi may confer significant evolutionary advantage (Nicolás *et al.*, 2013; Billmyre *et al.*, 2013).

Recent observations, particularly using the *Arabidopsis thaliana* - *Botrytis cinerea* pathosystem, have suggested that natural cross-kingdom gene silencing can occur in some plant-pathogen interactions (Weiberg *et al.*, 2013; Wang *et al.*, 2016). It has been demonstrated that *B. cinerea* expresses numerous sRNAs during infection of Arabidopsis, and that some of these sRNAs can inhibit accumulation of certain plant defense-related transcripts apparently facilitating fungal colonization and disease development (Weiberg *et al.*, 2013; Cai *et al.*, 2018). Therefore, it appears that some fungal sRNAs can function in a manner analogous to the pathogen effector proteins. Furthermore, the discovery that sRNAs can transit between plant and fungal cells to modify gene expression in the recipient cell, opens the possibility for novel crop protection strategies based on RNAi. One such strategy, known as host-induced gene silencing (HIGS), typically involves generation of transgenic plants expressing long dsRNAs or hairpin RNAs with homology to the essential pathogen mRNAs. Uptake of sRNAs generated *in planta* from these dsRNA species by the pathogen induces silencing of the target genes and ultimately suppression of the disease. HIGS has been demonstrated in several fungal and oomycete pathosystems (Nowara *et al.*, 2010; Koch *et al.*, 2013; Ghag *et al.*, 2014; Cheng *et al.*, 2015; Chen *et al.*, 2016; Song & Thomma, 2018; Qi *et al.*, 2018). To overcome potential difficulties with generating transgenic plant material and the associated GMO safety aspects, a new spray-induced gene silencing (SIGS) strategy involving exogenous application of synthetic dsRNA or siRNA molecules (‘RNA fungicides’) to the plants for the control of fungal pathogens has recently been described (Wang *et al.*, 2016; Koch *et al.*, 2016; Machado *et al.*, 2018; McLoughlin *et al.*, 2018).

The ascomycete fungus *Zymoseptoria tritici* is the causative agent of Septoria tritici blotch (STB) disease and is the major threat to bread and pasta wheat (*Triticum aestivum* and *Triticum durum*) production globally (Dean *et al.*, 2012). *Z. tritici* is a hemibiotrophic foliar pathogen, which invades leaf tissue through natural openings such as stomata. *Z. tritici* remains exclusively apoplastic through its infection cycle, which is characterized by an extended symptomless infection phase (10-14 days) followed by the rapid transition to necrotrophy (Kema *et al.*, 1996; Keon *et al.*, 2007). Considerable progress has been made in understanding the infection biology of *Z. tritici*, including the role of fungal secreted proteins in facilitating or hindering plant colonization (Marshall *et al.*, 2011; Kettles & Kanyuka, 2016; Zhong *et al.*, 2017; Kettles *et al.*, 2017; Kema *et al.*, 2018; Meile *et al.*, 2018). Whilst the existence of numerous *Z. tritici* resistance genes (*Stb* genes) in wheat germplasm has been known genetically for some time, cloning of the first of these genes (*Stb6*) was only described in 2018 (Saintenac *et al.*, 2018).

Recent developments in the fledgling field of cross-kingdom RNAi prompted us to ask whether this phenomenon was involved in the colonization of wheat by *Z. tritici*, an exclusively extracellular (apoplastically) dwelling pathogen. Here we report the identification and characterization of the sRNA populations generated by this fungal species during wheat leaf infection, predict putative wheat transcripts that may be subject to cross-kingdom RNAi and carry out validation of a subset of such interactions experimentally. The role of RNAi pathways in the infection biology of *Z. tritici* was assessed through the generation of targeted single gene deletion mutants. We also assessed whether *Z. tritici* has a capacity to uptake exogenously applied long dsRNA and sRNA and explored HIGS and *in vitro* RNAi as alternative approaches for characterizing fungal gene function and potentially also for control of this economically important fungal pathogen.

## 2 Materials and Methods

### 2.1 Plant and fungal material for small RNA sequencing

The *Zymoseptoria tritici* isolate IPO323 and wheat (*T. aestivum*) cv. Bobwhite, which is fully susceptible to this and many other *Z. tritici* isolates, were used in all experiments. Fungal Czapek-Dox Broth (CDB) cultures were propagated in shake flasks at 220 rpm and 15°C for 4 d and then harvested via filtration. Plant inoculation experiments were done as described previously (Rudd *et al.*, 2015) using a suspension of 1 x 10^7^ spores·mL^-1^ in water supplemented with 0.1% (v/v) Silwet L-77. Mock inoculations of plants were made using a 0.1% (v/v) Silwet L-77 water solution. Each biological replicate plant sample for RNA isolation was made up of five 6-cm long leaf segments each collected from a separate individual mock- or *Z. tritici-*inoculated plant randomly distributed in a single temperature-controlled glasshouse. Leaves and fungal *in vitro* samples were immediately frozen in liquid nitrogen and stored at −80°C before used for RNA purification.

### 2.2 RNA sequencing and bioinformatics analysis

Wheat cv. Bobwhite leaf tissue samples mock-inoculated and those inoculated with *Z. tritici* isolate IPO323 were collected at 4 dpi (asymptomatic stage), 9 dpi (first signs of host cell death), 13 dpi (extensive host cell death and a few fungal pycnidia initials visible), and 21 dpi (end of infection with numerous pycnidia visible), respectively. Healthy, untreated 17 days old wheat cv. Bobwhite leaf tissue were also collected to serve as an additional control. Samples were used for RNA extraction with the ZR RNA MiniPrep kit (Zymo Research). Concentration, quality and integrity of each RNA preparation was verified using NanoDrop Spectrophotometer (Thermo Fisher Scientific), Qubit 2.0 Fluorometer (Thermo Fisher Scientific), and 2100 Bioanalyzer (Agilent). sRNA libraries were constructed using the TruSeq small-RNA Library Preparation kit (Illumina) and sequenced as 50 cycle single end reads on a HiSeq 2000 (Illumina) by GENEWIZ. The sequencing data was deposited to the European Nucleotide Archive under the study PRJEB28454. The raw reads were trimmed using Trim Galore! (https://github.com/FelixKrueger/TrimGalore) to remove the adapter sequences with a minimum length cut-off 17 nt and trimmed fastq sequences analyzed using FastQC (https://www.bioinformatics.babraham.ac.uk/projects/fastqc/).

The trimmed reads were aligned to the genomes of *Z. tritici* IPO323 (https://genome.jgi.doe.gov/Mycgr3/Mycgr3.home.html) and wheat cv. Chinese Spring (ftp://ftp.ensemblgenomes.org/pub/plants/release-26/fasta/triticum_aestivum/dna/) using Bowtie 2 (Langmead & Salzberg, 2012). End-to-end model was used in Bowtie 2 and the score-min was set as L, 0, −0.1. All other parameters were default. Mapped reads from all the samples were merged together for sRNA loci discovery using SiLoCo (UEA sRNA toolkit; http://srna-workbench.cmp.uea.ac.uk/) with the following key parameters: maximum gap 50 nt, maximum sRNA size 30 nt, minimum sRNA size 19 nt, and minimum hit 100 reads. sRNA reads from each sample were counted exactly as they were based on the defined sRNA locus space from SiLoCo results and no apportioning method was applied to the read counts. Namely, if a read S was aligned to N places, read S were counted for N times. The typical sRNA loci, by contrast to the null loci, could be characterized by the significantly higher abundance of reads and the dominant read length distribution (rather than random read length distribution). Based on both read coverage depth and read length distribution, a Bayesian model is applied to estimate posterior probability for a given locus and used this as an indicator of likelihood for a locus to be a *bona fide* sRNA locus (Method S1). Briefly, sRNA loci were selected for further investigation based on the following four criteria: (1) the difference of the maximum posterior probability between the fungus containing samples (infected wheat and *in vitro* culture) and the fungus free samples (mock-inoculated healthy wheat) is > 0.8; (2) posterior probability for at least one of the infected wheat samples is > 0.8; (3) differential expression FDR is < 0.05 in at least one comparison between the infected wheat samples and the corresponding mock samples; and (4) dominant length of sRNAseq reads that map to the locus must be < 25 nt.

For the sRNA differential expression analysis, comparisons between *Z. tritici-*inoculated and the corresponding mock-inoculated samples were performed for each sRNA locus. The quantitative measurement of sRNA expression was based on reads counts. The raw data counts on sRNA locus space were normalized by the trimmed mean of M-values (TMM) method (Robinson & Oshlack, 2010). Differential expression analysis was performed using the edgeR package (Robinson *et al.*, 2010) and a generalized linear model using negative binomial distribution was applied.

For sRNA target prediction, the input sRNA sequences were 21-24 nt reads from each sRNA locus, and the target DNA regions were the wheat cv. Chinese Spring chromosome-specific cDNA sequence database produced by the International Wheat Genome Sequencing Consortium (IWGSC) in late 2015, at the time when this bioinformatics part of the study was done (ftp://ftp.ensemblgenomes.org/pub/plants/release-22/fasta/triticum_aestivum/cdna/). An initial sRNA target prediction was done using the microRNA target prediction tool (UEA sRNA toolkit). To meet more stringent target criteria, a Python script was written to filter out the initial results. Briefly, sRNA target prediction criteria followed the (Weiberg *et al.*, 2013) study: (1) no gap or bulge is allowed for sRNA/target duplex; (2) no mismatches in positions 10-11; (3) no adjacent mismatches in positions 2-12; (4) cutoff score ≤ 4.5, where the mismatch and G-U match penalty is 2 and 1, respectively in position 2-12 and mismatch penalty 1 and G-U match penalty 0.5 in other positions; and (5) minimum free energy (MFE) of sRNA-target duplex > 0.74.

The RNA Sequencing (RNAseq) data from the *Z. tritici* IPO323 infection time-course on a fully susceptible wheat cv. Riband and the corresponding mock-inoculated wheat leaf samples produced in our previous study ((Rudd *et al.*, 2015); the National Centre for Biotechnology Information Sequence Read Archive (SRA), project PRJEB8798) was re-analyzed by mapping the clean sequencing reads with Bowtie 2 (Langmead & Salzberg, 2012) to the chromosome-specific protein coding sequences (CDS) database of wheat cv. Chinese Spring (ftp://ftp.ensemblgenomes.org/pub/plants/release-26/fasta/triticum_aestivum/cds/). Transcript abundances were quantified from the SAMtools (Li *et al.*, 2009) sorted bam files using eXpress (Roberts & Pachter, 2013) and only the sequences with > 1 raw CPM in at least one time-treatment were kept for downstream analyses. Differential expression analysis was performed using the edgeR package (Robinson *et al.*, 2010) and a generalized linear model using negative binomial distribution was applied. The Benjamin-Hochberg false discovery rate (FDR) correction was used to adjust *p* values based on the exact Fisher test (Robinson *et al.*, 2010). The genes were considered differentially expressed if they had a log fold change ≥ 1 or ≤ −1 and FDR ≤ 0.05. Wheat transcript sequences were annotated using the Mercator pipeline (Lohse *et al.*, 2014).

### 2.3 RNA extractions for RT-PCR analyses

All sample material was snap frozen in liquid nitrogen and homogenized using a mortar and pestle. RNA was recovered using Trizol reagent (Invitrogen) following the manufacturer’s instructions, with a 3 h ethanol precipitation at −20°C to ensure maximal recovery of sRNA. Total RNA was treated with RQ1 DNase (Promega) following the manufacturer’s instructions. DNA-free total RNA was recovered by ethanol precipitation for 3 h at −20°C followed by quantification using a NanoDrop spectrophotometer.

### 2.4 qRT-PCR

1-2 µg DNA-free total RNA was used as template for cDNA synthesis using Superscript III (Invitrogen). cDNA was diluted 1:10 with distilled H2O prior to assembling reactions. SYBR Green JumpStart Taq ReadyMix (Sigma) reactions were assembled following the manufacturer’s instructions. Reactions were run on a CFX384 real-time PCR system (Biorad) using the following thermocycle: 95°C for 3 min, then 40 cycles of 95°C (30 s), 60°C (30 s), 72°C (30 s). Followed by melt curve analysis of products: 95°C (10 s), then 65 - 95°C at 0.5°C increments with 5 s at each.

### 2.5 Stem-loop qRT-PCR

Reactions were assembled following the stem-loop pulsed reverse transcription protocol and the miRNA SYBR Green 1 assay protocol, both previously described (Varkonyi-Gasic *et al.*, 2007) with the exception that SYBR Green JumpStart Taq ReadyMix (Sigma) was used. Reactions were run on a CFX384 real-time PCR system (Biorad) using the following thermocycle: 95°C for 2 min, then 40 cycles of 95°C (15 s), 60°C (15 s), 72°C (15 s). Followed by melt curve analysis of products: 95°C (10 s), then 65 - 95°C at 0.5°C increments with 5 s at each.

### 2.6 Fungal mutagenesis

Generation of gene deletion strains using *Agrobacterium tumefaciens* mediated transformation of *Z. tritici* IPO323 *Δku70* strain (Bowler *et al.*, 2010) has been described previously (Motteram *et al.*, 2009). Several independent transformants with each gene deleted were validated by PCR on genomic DNA. Primer sequences used for the development of single gene deletion constructs and validation of transformants are shown in Table S1.

### 2.7 Fungal bioassays

Inoculation of transgenic *Z. tritici* strains to wheat seedlings and assessment of disease was done following a previously described procedure (Motteram *et al.*, 2009; Rudd *et al.*, 2015).

### 2.8 5’-RACE

Assays were performed using components of the GeneRacer kit (Invitrogen) using the method of Llave *et al.* (2002). DNA-free RNA from both mock (negative control) and fungus-infected plant tissue were used as templates. The interaction between Ta-miR156 and the *TaSPL3* transcript was used as a positive control (Liu *et al.*, 2017).

### 2.9 *Barley stripe mosaic virus* (BSMV)-mediated HIGS

Approximately ∼300-bp fragments of coding sequences (CDS) of *Z. tritici* genes *ZtTUBa* (*Mycgr3G76039*), *ZtTUBb* (*Mycgr3G102950*), *ZtCYP51* (*Mycgr3G110231*), and *ZtALG2* (*Mycgr3G75289*) or the Green Fluorescent Protein (GFP)-encoding gene were selected for development of BSMV-HIGS constructs following interrogation of *Z. tritici* IPO323 and wheat transcript databases using si-Fi v. 3.1.0 (siRNA Finder; http://labtools.ipk-gatersleben.de/) software. These fragments were then amplified by PCR from total cDNA from an *in vitro* cultured *Z. tritici* IPO323, the full-length cDNA clone WT-CYP51 (kindly provided by Hans Cools, Rothamsted Research) or the BSMVγ::GFP plasmid (Haupt *et al.*, 2001) using Invitrogen Taq DNA Polymerase (Thermo Fisher Scientific) and primers carrying 5’ ligation independent cloning (LIC) adaptors specified in Table S1. Each PCR product was cloned into the binary BSMV vector pCa-γbLIC using LIC (Yuan *et al.*, 2011) in antisense orientation. Sequence verified HIGS constructs pCa-γbLIC:asZtTUBa, pCa-γbLIC:asZtTUBb, pCa-γbLIC:asZtCYP51, pCa-γbLIC:asZtALG2, and pCa-γbLIC:asGFP were then transformed into the *A. tumefaciens* strain GV3101 by electroporation and used in conjunction with *A. tumefaciens* strains transformed with pCaBS-α (BSMV RNAα) and pCaBS-β (BSMV RNAβ) for Agrobacterium mediated inoculation of *Nicotiana benthamiana* plants following a procedure described in Lee *et al.* (2015b). Sap from the virus infected *N. benthamiana* plants was used for rub-inoculation of wheat cv. Riband plants. Wheat inoculation with BSMV:asTaChlH served as a positive control for RNA silencing (Lee *et al.*, 2015b). At 14 days post virus inoculation, the BSMV-HIGS-treated plants were challenge inoculated with *Z. tritici* IPO323 or the transgenic, GFP-expressing strain B3 (Rohel *et al.*, 2001) and progression of fungal infection was visually monitored during the next 22-25 days, following the final visual assessment of the disease as well as by quantifying asexual fungal sporulation following the procedures previously described (Lee *et al.*, 2015b).

### 2.10 *In vitro* RNAi

Pairs of PCR primers, each carrying the same extension sequence (5’-tcctaatacgactcactatagggag-3’) at their 5’-end, that corresponds to a binding site for T7 RNA Polymerase, were used for amplification of ∼300-bp fragments of CDS of *Z. tritici* genes *ZtTUBa*, *ZtTUBb*, and *ZtCYP51* or the negative control GFP-encoding gene (Table S1). The pCa-γbLIC derivatives described above were used as templates for PCRs that were performed using Invitrogen Taq DNA Polymerase (Thermo Fisher Scientific). The resulting PCR products served as templates for the synthesis of double-stranded RNA (dsRNA) using the Ambion® MEGAscript® RNAi kit (Thermo Fisher Scientific) following the manufacturer’s instructions. Three different amounts of dsRNA (12.5 ng, 125 ng and 1250 ng) in 2.5 µl elution buffer (Ambion® MEGAscript® RNAi kit) were mixed with 100 µl water suspensions of *Z. tritici* IPO323 conidiospores at four different concentrations (5 x 10^6^, 5 x 10^5^, 5 x 10^4^, and 5 x 10^3^ mL^-1^) in wells of 96-well culture plates. The plates were incubated at room temperature for 1 h, 4 h and 20.5 h following which the two 5 µl aliquots of germinating spores were taken from each well and spotted as replicas onto Petri dishes containing Yeast Peptone Dextrose Agar (YPDA). The YPDA plates were incubated at 17°C in the dark and the appearance and growth of fungal colonies was monitored macroscopically and by viewing under a binocular microscope over the period of 7 days.

Capacity to uptake exogenously applied control long dsRNA and sRNA by *Z. tritici* and *B. cinerea* was assessed as follows. Fungal material was harvested by scratching from culture plates and adjusting in liquid minimal synthetic nutrient medium (SNA) to approximately 10^5^ spores·mL^-1^. Fungal spore preparations were co-incubated with 100 nM BLOCK-iT Alexa Fluor Red siRNA (ThermoFisher catalog No. 14750100) or with 1 µg Cy3-labelled *in vitro*-transcribed 250-bp long dsRNA against GFP in microtiter plates at 21°C, shielded from light. Uptake of fluorescently labelled RNA molecules by germinated spores extending hyphae was monitored after 12 h and 48 h of incubation using a fluorescence microscope, and images were captured with bright field and fluorescent (absorption 555 nm, emission 565 nm) settings.

## 3 Results

### 3.1 Identification and validation of fungal sRNA transcriptionally-induced *in planta*

The availability of a fully sequenced genome for *Z. tritici* isolate IPO323 allowed identification of the key components of RNA silencing machinery in this fungal species. Only one gene, *Mycgr3G47983*, is annotated as the “DsRNA-specific nuclease” in the *Z. tritici* IPO323 genome (Ensembl Fungi, release 41). BLAST analyses using this gene (named *ZtDCL*), the corresponding protein sequence, or the well-characterized *DCL-1* (*NCU08270*) and *DCL-2* (*NCU06766*) genes of the model fungal species *Neurospora crassa* against the *Z. tritici* IPO323 genome revealed no additional sequences potentially encoding DCLs. Similar analyses revealed the presence of four candidate AGO protein encoding genes: *Mycgr3G38035, Mycgr3G10621, Mycgr3G90232, Mycgr3G25632*. However, only the first two (named *ZtAGO1* and *ZtAGO2*, respectively) appeared to be functional with the other two coding for truncated proteins missing one or more essential N-terminal domains and showing very low level of expression (Figure S1; Table S2). Two candidate RdRp encoding genes, *Mycgr3G51407* and *Mycgr3G49833*, were also identified. This provided indirect evidence that a system for sRNA biogenesis may exist in *Z. tritici*.

We therefore carried our deep sequencing of sRNA preparations from *in vitro* cultured fungus, as well as from the infected fully susceptible wheat cv. Bobwhite plants. Over 9.5 million reads ranging in size from 16 to 52 nt that mapped to the fungal genome were obtained from the *in vitro* samples. Moreover, over 0.74, 1.28, 2.65 and 3.45 million reads that mapped to the *Z. tritici* IPO323 genome were obtained from the infected wheat leaf tissues sampled at the four different time points corresponding to the four critical stages of disease development, namely 4 dpi (“biotrophic”/ asymptomatic stage), 9 dpi (transition from “biotrophic” to “necrotrophic” stage), 13 dpi (“necrotrophic” stage), and 21 dpi (profuse asexual sporulation) (Rudd *et al.*, 2015). This provided direct evidence that *Z. tritici* is capable of generating sRNAs during both *in vitro* and *in planta* growth. The fungal sRNA length distribution was much broader than that typical of vascular plants. Also, in addition to the typical ∼ 20-24 nt sRNA peaks, two extra peaks centered at ∼ 30 nt and ∼ 50 nt were observed in the samples (Figure S2). Further analysis indicated that the majority of these longer sequences originate either from tRNA or rRNA loci, or from wheat chloroplast sequences (data not shown).

All adapter trimmed sRNA reads that mapped to the *Z. tritici* IPO323 genome without mismatches and had no perfect matches in the wheat genome were used for sRNA loci prediction using SiLoCo. This resulted in the identification of 1619 candidate sRNA loci. sRNA reads were counted based on the defined sRNA loci for each sample. Based on the read count abundance and read length distribution, posterior probability per sRNA locus per sample was estimated. sRNA loci differential expression between infected and mock samples were also estimated. sRNA loci were selected for further investigation following a pipeline described in the Materials and Methods, and Method S1. All sRNA loci annotated as rRNAs and tRNAs were excluded from further analysis apart form a small number of those that mapped to the non-coding strand of these genes. This analysis funneled down the candidate sRNA loci list from 1619 to 389 potentially genuine sRNA loci that differed in length and total expression level and originated predominantly from intergenic regions (76%) (Figure S3; Table S3). These loci were active at one or sometimes at several often adjacent timepoints during wheat infection (Figure 1). Notably, all chromosomes of *Z. tritici* including the 13 core and 8 shorter accessory (dispensable) chromosomes (Goodwin *et al.*, 2011) were predicted to host sRNA loci, and there was a trend for longer chromosomes hosting higher number of sRNA loci (Figure S4; Table S3).

**Figure 1.**
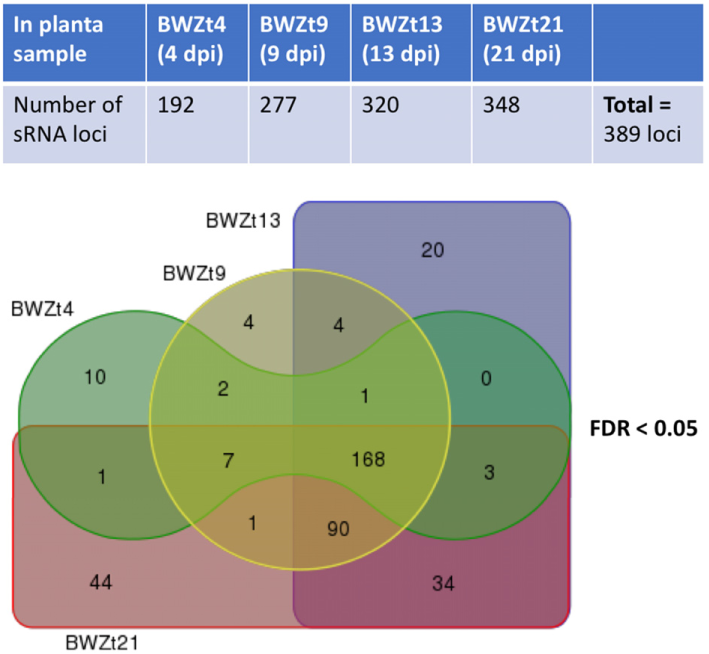
Predicted *Zymoseptoria tritici* sRNA loci active during different timepoints of wheat infection. Numbers of sRNA loci upregulated at 4, 9, 13, and 21 days post inoculation (dpi) of a susceptible wheat cv. Bobwhite with *Z. tritici* isolate IPO323 are shown in the upper panel. A Venn diagram showing numbers of infection stage-specific sRNA loci as well as those shared between the consecutive infection timepoints is presented in the lower panel. False discovery rate (FDR) rate for differential sRNA loci expression in at least one comparison between the infected wheat samples and the corresponding mock samples was set to < 0.05.

To verify the sRNAseq data, we performed a new *Z. tritici* IPO323 infection time course on the susceptible wheat cv. Bobwhite, sampling the leaf tissue at 4 dpi, 9 dpi, and 13 dpi. RNA from this material was used to assess an *in planta* expression of four fungal sRNAs, selected from the sRNAseq dataset, using a sequence guided stem-loop qRT-PCR assay (Figure 2). The same technique was also used to evaluate expression of these sRNAs in liquid (CDB) fungal culture (Figure 2). Expression of these sRNAs in the infected plant material was found to be considerably higher than in the fungus cultured *in vitro* (Figure 2). Expression was highest at the 4 dpi and 9 dpi timepoints, which correspond to the symptomless and transition phases of colonization by this fungus. By 13 dpi, when significant regions of necrotic tissue were present in leaves infected under glasshouse conditions, the expression of all four fungal sRNAs had returned to levels similar to those identified *in vitro* (Figure 2). These experiments demonstrated that all fungal sRNAs investigated were significantly induced during wheat infection, thereby validating the data generated by sRNAseq.

**Figure 2.**
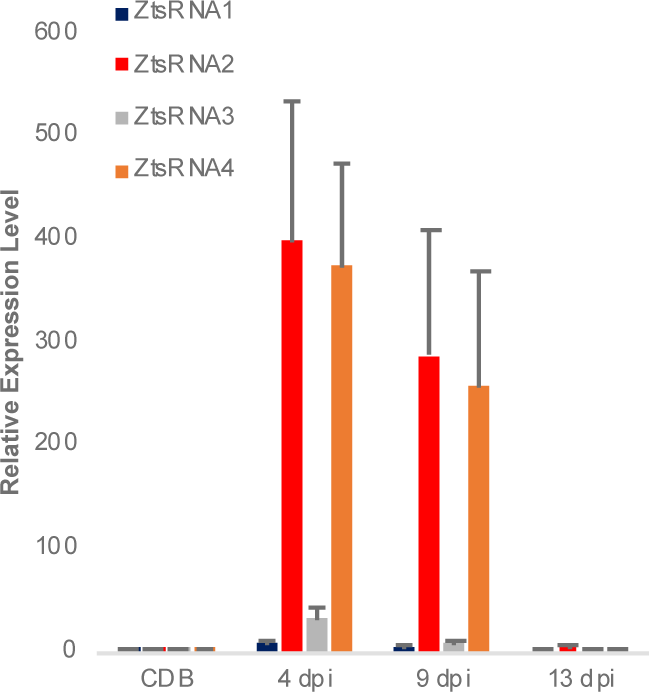
Expression profiling of *Zymoseptoria tritici* mature sRNAs by stem-loop qRT-PCR. Relative expression of four *Z. tritici* sRNAs (ZtsRNA1 - ZtsRNA4) from the fungus grown *in vitro* (CDB) or *in planta* (leaf tissue at 4, 9, and 13 days post inoculation, dpi). Bars indicate SE.

### 3.2 sRNA target prediction: identification and validation of candidate wheat transcripts repressed during fungal infection

For a given sRNA locus, 21-24 nt sequences were retrieved from each infected sample alignment bam file for target prediction on transcripts of the wheat cv. Chinese Spring (the only wheat genotype for which the draft genome sequence was available at the time of the study). Here we used sRNA target prediction criteria similar to those utilized in the previously published study (Weiberg *et al.*, 2013). This identified a total of 262 sRNAs, with 140 of which having unique sequences and 122 others falling into 43 sequence related groups (Table S4), that were computationally predicted to target 737 wheat transcripts (Table S5). Majority of these sRNAs were 21 nt long and displayed a bias towards uridine in their 5’ termini (Figure S5), features characteristic of functional sRNAs capable of directing cleavage or translational silencing of the corresponding target mRNAs.

It was expected that the wheat transcripts successfully targeted by fungal sRNAs would display significant downregulation during either specific or all stages of the *Z. tritici* – wheat infection. To predict wheat genes whose expression was significantly reduced during fungal infection we re-analyzed our previously published RNAseq data set from the *Z. tritici* IPO323 infection time course on the fully susceptible wheat cv. Riband and the corresponding mock-inoculated plants (Rudd *et al.*, 2015) by re-mapping the sequence reads to the wheat cv. Chinese Spring chromosome-specific CDS ‘Triticum_aestivum.IWGSC2.26.cds.all.fa’ followed by differential gene expression analysis. This data helped us to prioritize 10 wheat mRNAs, predicted to be targeted by the four selected *Z. tritici* sRNAs for cleavage, for further investigation based on their expression profile and potential role in pathogen defense (Table 1; Table S5). Most of these wheat transcripts exhibited reduced expression in the infected wheat cv. Riband tissue compared to the corresponding mock-inoculated healthy control plants in at least one of the three assessed timepoints, with the expression levels often being lower at 9 dpi and 14 dpi (transition and necrotrophic phases) than at 4 dpi (symptomless phase) (Figure 3). However, there were also some instances where expression of predicted target mRNAs was higher in the infected tissue in comparison to the mock-treated controls, specifically for Traes_2AS_EC975D5AB and Traes_5BS_409B24307 at 4 dpi and for Traes_2DS_9B86CE58D at 14 dpi (Figure 3). A qRT-PCR was then used to analyze expression of these predicted target mRNAs during *Z. tritici* IPO323 infection of wheat cv. Bobwhite (Figure 3). All target wheat mRNAs again showed downregulation in the infected leaf tissue compared to the mock-treated controls in at least one infection timepoint. Therefore, the qRT-PCR data for wheat cv. Bobwhite agreed well with the expression data obtained using RNAseq for wheat cv. Riband, indicating similar regulation of expression of these mRNAs in the two different fully susceptible wheat cultivars. Expression profiles of the two mRNAs, Traes_2BS_E5732CD2C and Traes_4BS_5E12F0B27, showed some differences between the two cultivars (Figure 3). However, alternatively, this may be due to qRT-PCR assays being more effective than RNAseq in distinguishing between highly similar homoeologous transcripts.

**Figure 3.**
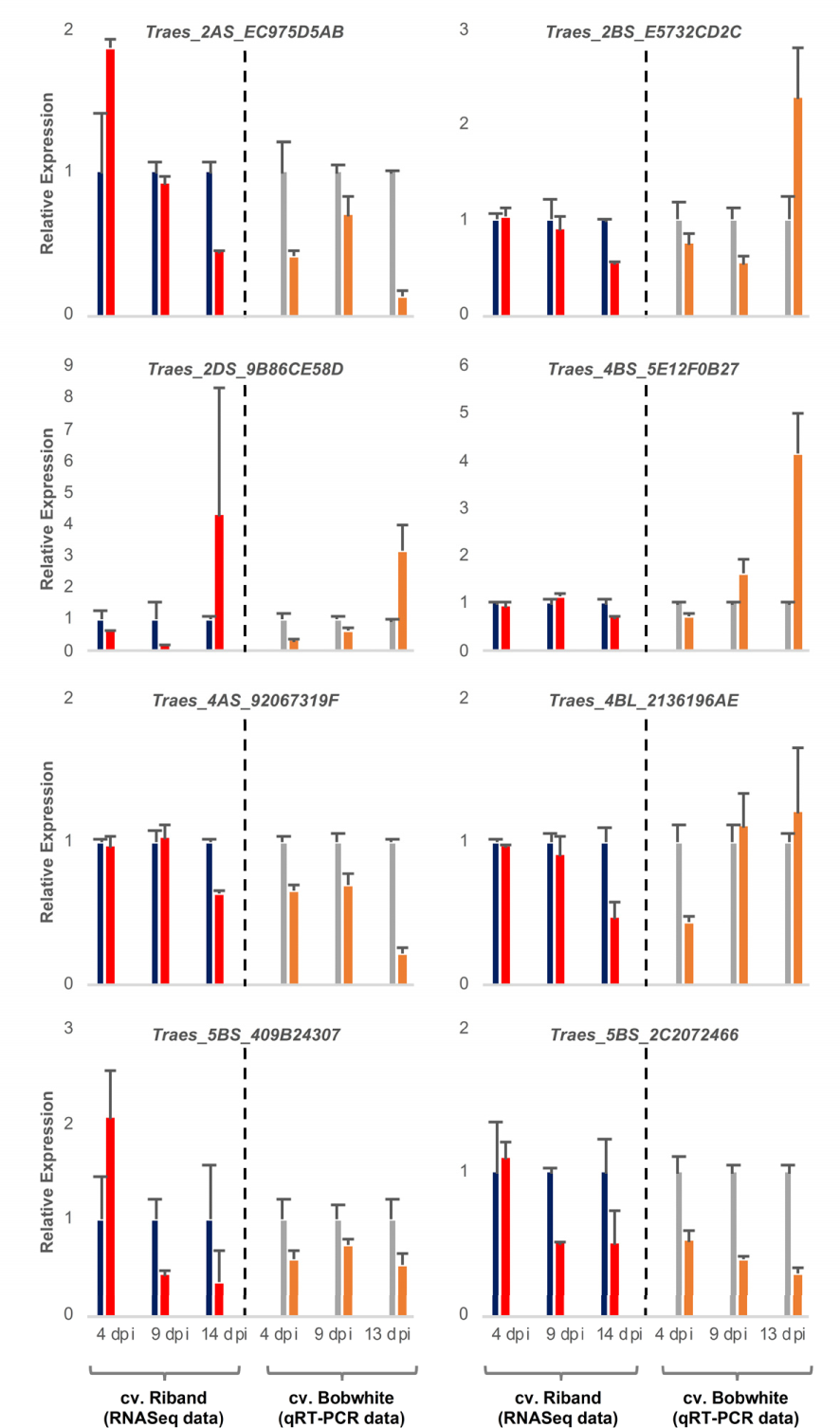
Expression profiling of eight candidate wheat target mRNAs during *Zymoseptoria tritici* IPO323 infection of wheat cv. Bobwhite leaf tissue. Left panels show RNAseq data (re-analysis of Rudd *et al.* 2015) for candidate genes converted to relative expression for presentation for mock (blue) and *Z. tritici* IPO323 (red) treatments. Right panels show qRT-PCR analysis of duplicate sample set, mock (grey) and *Z. tritici* IPO323 (orange). Bars indicate SE.

**Table 1.**
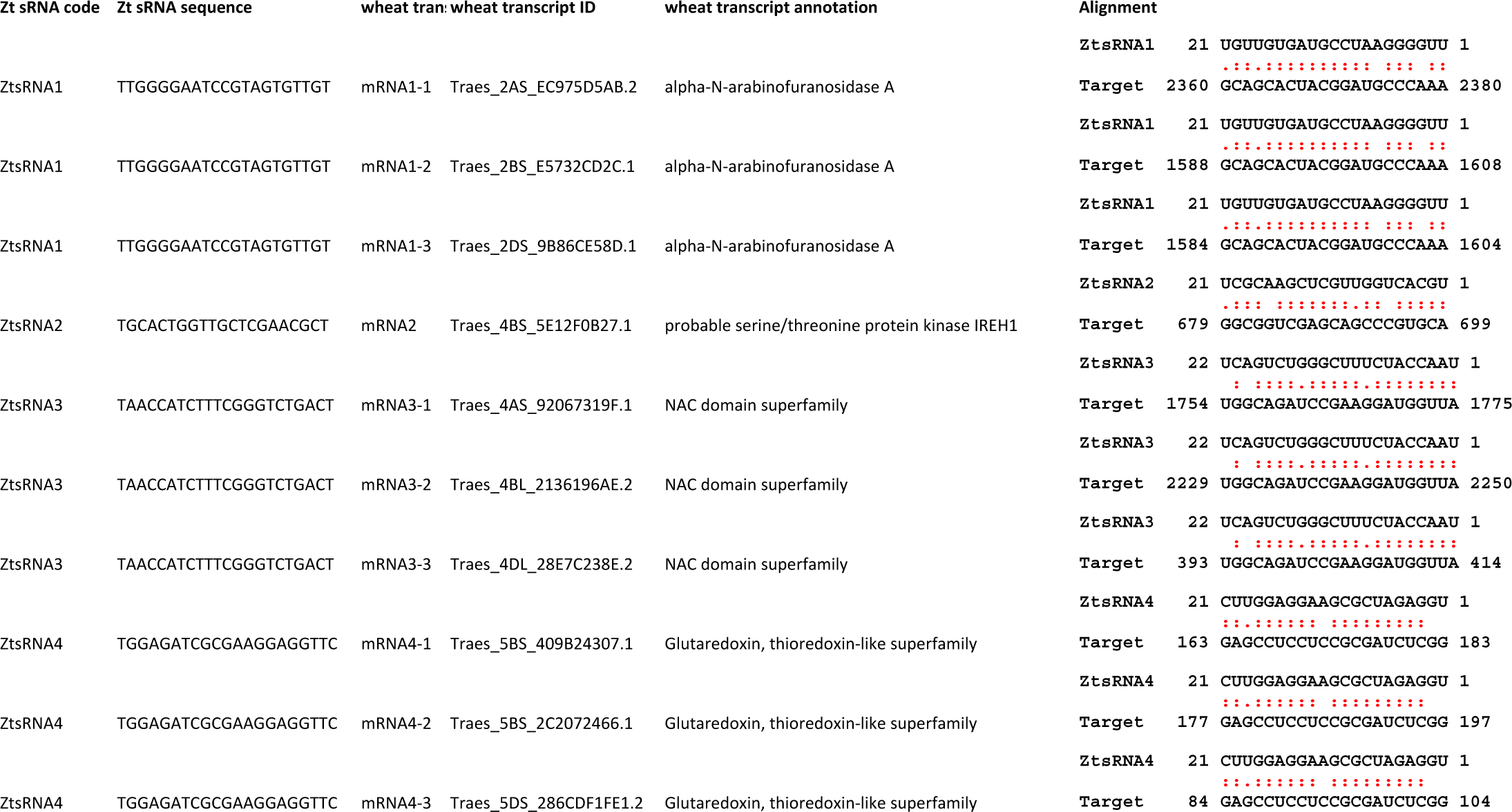
*Zymoseptoria tritici* IPO323 mature sRNAs selected for detailed study and their respective wheat mRNA targets..

### 3.3 RNAi-deficient fungal mutant strains are fully pathogenic on wheat

To further examine the potential role of the fungal RNAi pathway in a compatible interaction between wheat and *Z. tritici* we produced several *Z. tritici* mutants deficient in the key RNAi components. Gene deletion strains for the single predicted *ZtDCL* gene and both of the genes, *ZtAGO1* and *ZtAGO2*, predicted to encode Argonaute proteins were generated and validated by PCR. All mutant strains were produced in the *Z. tritici* IPO323 *Δku70* background (Bowler *et al.*, 2010) which maximizes homologous recombination. Three independent mutants for each target gene were selected for further study. In replicated infection bioassays using the susceptible wheat cv. Bobwhite, all generated mutant strains, *Δdcl*, *Δago1*, and *Δago2*, displayed virulence phenotypes indistinguishable to the parental IPO323 *Δku70* strain (Figure 4). The rate of disease lesion development on the infected leaves was similar to the IPO323 *Δku70* control in all experiments, with inoculation zones becoming fully necrotic by 16 dpi (Figure 4). These experiments therefore demonstrated that *ZtDCL*, *ZtAGO1*, and *ZtAGO2* genes are dispensable for virulence.

**Figure 4.**
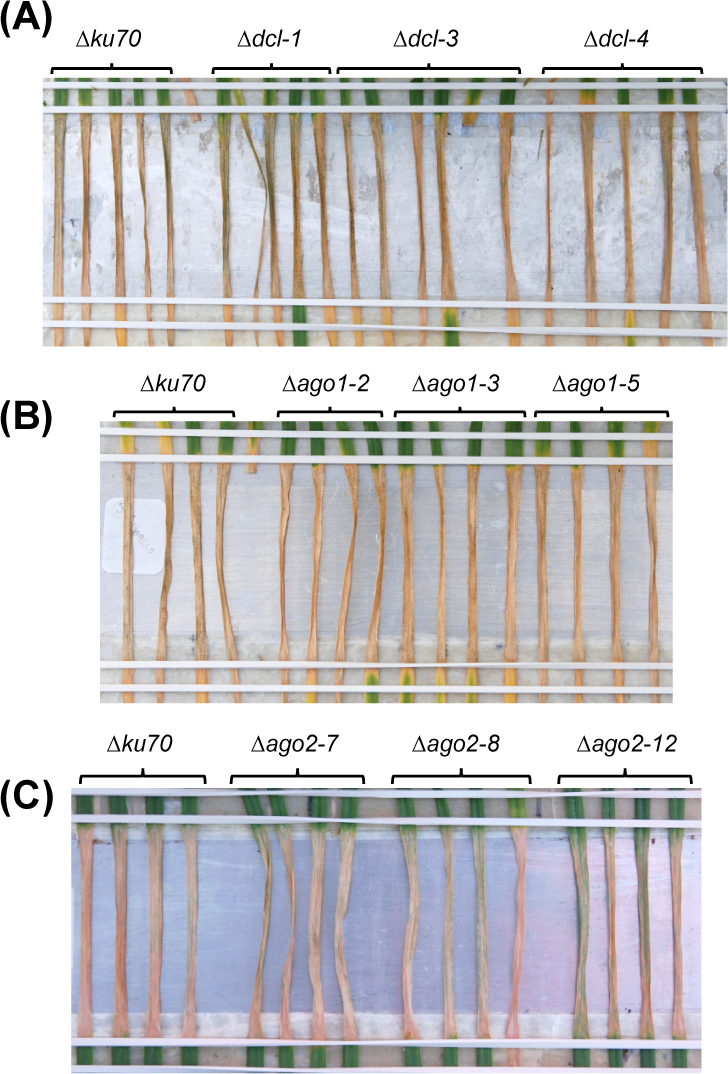
*Zymoseptoria tritici* infection bioassays. *Z. tritici* IPO323 strain *Δku70* and its derivatives deficient in RNA silencing pathway (*Δdcl*, *Δago1* or *Δago2*) were inoculated onto the wheat cv. Bobwhite. Fungal inoculations were done using suspension of conidiospores at 1 x 10^6^ mL^-1^ and the inoculated leaves were photographed at 16 days post inoculation.

### 3.4 Evidence for DCL-independent biogenesis of sRNA in *Z. tritici*

As described above, the *Z. tritici* mutant *Δdcl* displayed no reduction in virulence in the wheat infection bioassay. To further investigate the role of fungal sRNAs as potential virulence factors during infection by this foliar pathogen, we carried out new wheat infection time courses with both *Z. tritici* IPO323 *Δku70* (control) and the *Δdcl* mutant followed by stem-loop qRT-PCR analysis of fungal sRNA expression (Figure 5). Unexpectedly, expression of three out of four fungal sRNAs investigated, namely ZtsRNA1, ZtsRNA3 and ZtsRNA4, was maintained in the *Δdcl* mutant at levels comparable to those in IPO323 *Δku70* control (Figure 5). Moreover, as with the control strain, sRNA expression of these 3 sRNAs in the *Δdcl* mutant was considerably higher when the fungus was recovered from the infected wheat leaf tissue in comparison to the fungus cultured *in vitro* (Figure 5). Highest sRNA expression levels were detected at either 4 dpi (ZtsRNA1, ZtsRNA3) or 9 dpi (ZtsRNA4). In contrast, expression of ZtsRNA2 was abolished in the *Δdcl* mutant (Figure 5). No expression of this sRNA was detected in the fungus grown either *in vitro* or *in planta*. This data suggested that *ZtDCL* is required for the biogenesis of ZtsRNA2, whereas the other three sRNAs may be generated independently of *ZtDCL*.

**Figure 5.**
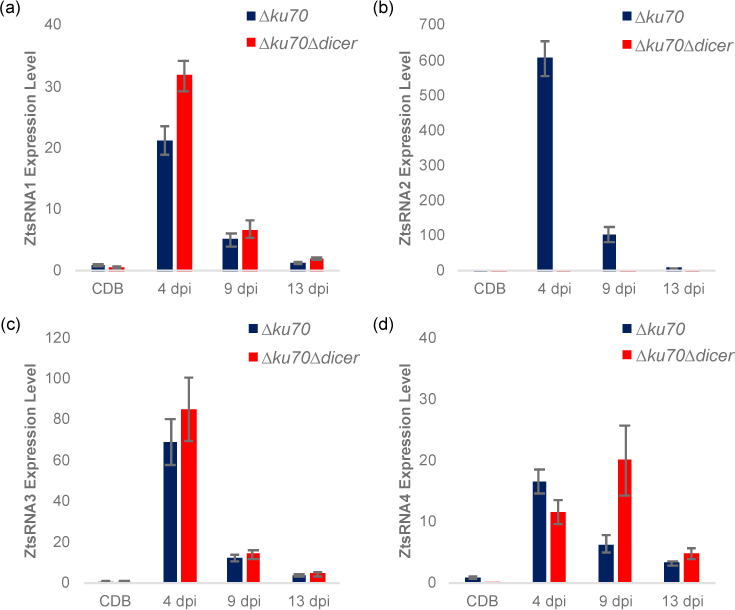
Expression profiling of fungal sRNAs in an RNAi-deficient *Zymoseptoria tritici* mutant *Δdicer* (IPO323 *Δku70 Δdcl*). Stem-loop qRT-PCR analysis of four ZtsRNAs from *Z. tritici* IPO323 *Δku70* (blue) and *Z. tritici* IPO323 *Δku70 Δdcl* (red). Fungal material from an *in vitro* culture (Czapek Dox Broth, CDB) and *in planta* (leaf tissue at 4, 9 and 13 days post inoculation, dpi) was assessed. Bars represent the mean and SE. The *Z. tritici* IPO323 *Δku70* (CDB) samples were set to 1 in all panels.

To further examine the role of ZtsRNA2 in fungal virulence, a qRT-PCR analysis was performed on the same sample set to assess expression levels of the predicted wheat target mRNA Traes_4BS_5E12F0B27. Surprisingly, expression of this mRNA was similar in the wheat leaf tissue infected with either *Z. tritici* IPO323 *Δku70* or *Δdcl* at all three timepoints (Figure S6). Given the absence of ZtsRNA2 in the *Z. tritici Δdcl* mutant, this indicated that Traes_4BS_5E12F0B27 is likely not a genuine target of this fungal sRNA. To verify this, and to assess the interactions between the three other *Z. tritici* sRNAs and their putative targets, 5’-RACE assays were performed to detect mRNA cleavage fragments resulting from targeting. These assays were unable to detect any fragments associated with the predicted sRNA-guided target cleavage (Figure S7). Therefore, it appears that none of the wheat mRNAs investigated represent genuine targets of the four fungal sRNAs, ZtsRNA1 - ZtsRNA4.

### 3.5 Transient host-induced gene silencing (HIGS) is ineffective in protecting wheat plants against Septoria tritici blotch disease

The data presented above suggested that although *Z. tritici* produces numerous sRNAs both *in vitro* and during infection of wheat, at least those that are *ZtDCL-*dependent do not seem to contribute to fungal virulence. Next, we asked whether the plant-derived sRNAs could translocate and induce RNAi in *Z. tritici* during infection. To address this, we used a plant RNA virus-based vector as a potent RNA silencing inducer in a procedure known as transient HIGS (Lee *et al.*, 2012). In this procedure, a recombinant *Barley-stripe mosaic virus* (BSMV) carrying a fragment of target fungal pathogenicity or essential for life gene is inoculated onto wheat leaves, which triggers a massive production of sRNAs from the replicating virus genome (i.e. long double-stranded RNA, dsRNA) by the plant’s own RNAi machinery (Lee *et al.*, 2012). Uptake of these long dsRNAs or fungal-gene specific sRNAs by *Z. tritici* during attempted infection of BSMV-HIGS-treated plants would be expected to result in substantially reduced disease levels.

Four *Z. tritici* genes were selected for silencing in this experiment: *ZtCYP51* (*Mycgr3G110231*; encodes cytochrome P450 lanosterol C-14a-demethylase), *ZtTUBa* (*Mycgr3G76039*; encodes α-tubulin), *ZtTUBb* (*Mycgr3G102950*; encodes β-tubulin), and *ZtALG2* (*Mycgr3G75289*; encodes α-1,2-mannosyl-transferase). The first three of these genes are known fungicide targets (Siegel, 1981; Hollomon *et al.*, 1998), whereas *ZtALG2* is essential for hyphal growth and pathogenicity (Motteram *et al.*, 2011). The similarly sized (∼ 300-bp) fragments of protein coding sequences were PCR amplified and cloned individually into the BSMV vector pCa-γbLIC in anti-sense orientation. As predicted *in silico* using si-Fi v. 3.1.0 software, all selected target CDS fragments were likely to generate comparable numbers of silencing-efficient small interfering RNAs (siRNAs). Moreover, the selected CDS fragments were not likely to generate siRNAs capable of inducing silencing of off-target genes in *Z. tritici* (except for *ZtTUBa* construct; see Table S6) or in its host, wheat. BSMV vectors carrying ∼ 300-bp fragments of a non-fungal and a non-plant origin gene, such as the jellyfish *Green Fluorescent Protein* (*GFP*) gene, and the wheat *Magnesium Chelatase subunit H* (*TaMgChlH*) gene involved in chlorophyll biosynthesis were used as the negative and positive controls for gene silencing, respectively (Lee *et al.*, 2015b). As expected, leaves of plants pre-treated with BSMV:asTaMgChlH appeared orangey-yellow rather than green (Figure S8), reporting successful silencing of *TaMgChlH* that resulted in chlorophyll deficiency (Lee *et al.*, 2015b). The rate of disease lesion development on wheat leaves infected with the two different *Z. tritici* strains, IPO323 and B3, was similar on all experimental plants i.e. those untreated with BSMV and also those pre-treated with the various BSMV-HIGS constructs (apart from BSMV:asTaMgChlH) or a negative control BSMV:asGFP. Visually similar levels of STB disease were observed by 22-25 days post fungal inoculation, when *Z. tritici* had completed its life cycle (data not shown). The disease was further quantified by counting *Z. tritici* pycnidiospores washed from the diseased leaves to verify the visual assessment. Indeed, very similar counts of pycnidiospores were obtained in washes from the virus-free wheat plants and from those that were pre-treated with BSMV:asGFP or the various BSMV-HIGS constructs (for the exception of BSMV:asTaMgChlH) prior to fungal inoculation (Figure 6). The number of fungal spores washed from each leaf was somewhat higher in the experiments involving *Z. tritici* IPO323. This may be either due to the possible differences in virulence between the different fungal strains, or because in the experiments involving *Z. tritici* IPO323 the leaves were sampled 3 days later post fungal inoculation and incubated under high relative humidity for 42 h rather than 24 h (before counting pycnidiospores) compared to the experiments involving *Z. tritici* B3 (Figure 6). By contrast and in agreement with our previous study (Lee *et al.*, 2015a) the ability of *Z. tritici* to complete its asexual infection cycle was severely impaired in *TaMgChlH*-silenced wheat plants as only a very small number of pycnidiospores was recovered in washes from these plants (Figure 6). This suggested, albeit indirectly, that the plant sRNA generating silencing pathways were successfully activated in these BSMV-mediated gene silencing experiments. However, these experiments failed to generate any evidence suggesting that BSMV-HIGS could work successfully to protect wheat plants against STB disease.

**Figure 6.**
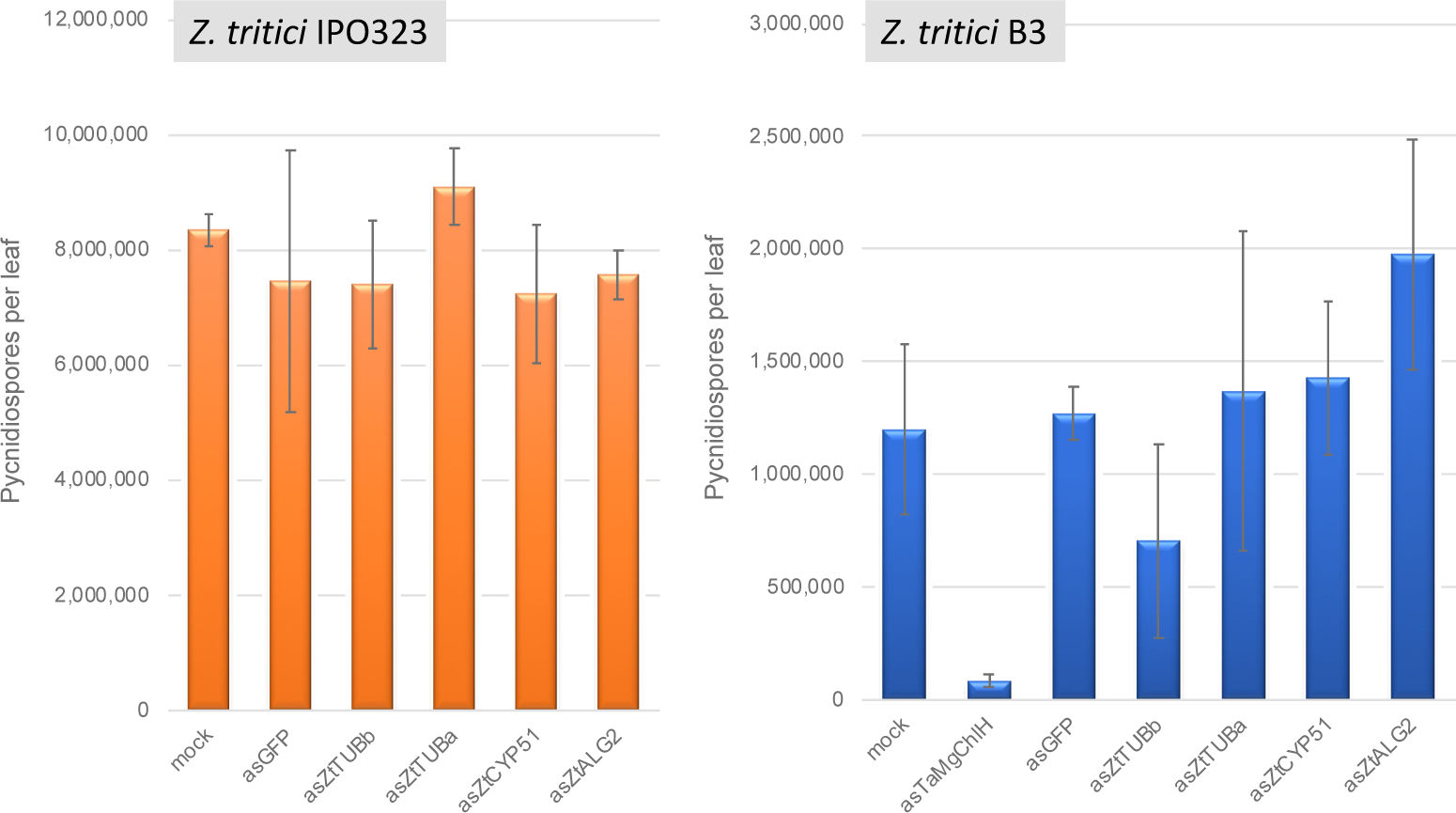
Assessment of *Zymoseptoria tritici* strains IPO323 and B3 pycnidiospore production in wheat cv. Riband plants pre-treated with various BSMV-HIGS and BSMV-VIGS constructs.

### 3.6 *Z. tritici* is unable to uptake externally applied dsRNAs *in vitro*

In parallel to the transient HIGS experiments described above, we also investigated whether external application of dsRNAs targeting essential for life fungal genes could impair or restrict *Z. tritici* growth *in vitro*. We, therefore, treated preparations of germinating *Z. tritici* conidiospores with *in vitro* synthesized ∼ 300 nt long dsRNAs targeting *ZtCYP51*, *ZtTUBa* and *ZtTUBb* genes. All these genes as mentioned above are known targets for fungicides, and knock-out mutations in these genes are thought to be lethal as none have been reported to date. GFP-specific dsRNA of a similar size was used as a negative control. The same gene regions of these genes, as in the BSMV-HIGS experiments described above, were selected to serve as templates for production of dsRNAs. Four different concentrations of *Z. tritici* conidiospore preparations (i.e. 10x serial dilutions ranging from 5 x 10^6^ mL^-1^ to 5 x 10^3^ mL^-1^) were incubated in sterile deionized water, which stimulates spores germination and hyphal extension (King *et al.*, 2017), with the three increasing amounts of dsRNAs (12.5 ng, 125 ng and 1250 ng per 100 µL volume) for 1 h, 4 h and overnight. After co-incubations with dsRNA, aliquots of fungal suspensions were spotted onto YPD agar plates and incubated at 17°C in the dark for 5-7 days or until fungal colonies could be visualized. No differences were observed in the appearance or growth rates between the colonies originating from untreated *Z. tritici* samples versus those treated with the three different amounts of dsRNA for any of the dsRNA treatments (Figure 7, and data not shown). This suggested that *Z. tritici*, in contrast to other fungal species such as *B. cinerea*, *Verticillium dahliae*, *Sclerotinia sclerotiorum*, *Fusarium graminearum* and *Fusarium asiaticum* (Wang *et al.*, 2016; Koch *et al.*, 2016; McLoughlin *et al.*, 2018), either unable to uptake external long dsRNA or that dsRNAs are ineffective in triggering gene silencing in this fungal species.

**Figure 7.**
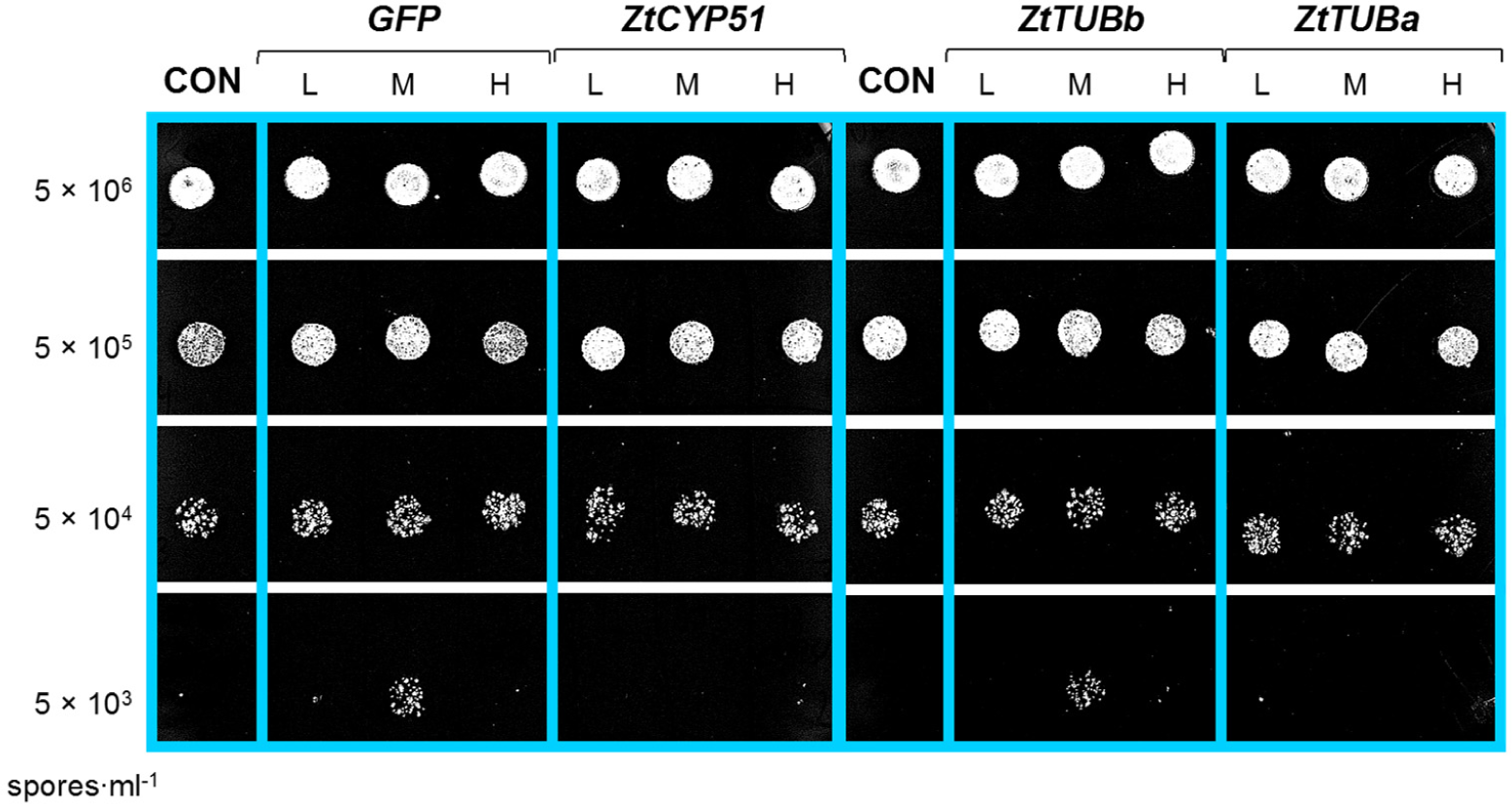
*In vitro* silencing experiment using *Zymoseptoria tritici*-specific dsRNAs generated in vitro. One hundred microliters of *Z. tritici* IPO323 conidiospore suspensions at four different concentrations (indicated on the left) were treated overnight with 12.5 ng (L), 125 ng (M) or 1250 ng (H) of dsRNA specific for different *Z. tritici* mRNAs before plating out onto YPD agar plates and growing for 4 days at 17°C in the dark. CON, untreated conidiospores of *Z. tritici*.

To directly assess the uptake of dsRNA by *Z. tritici*, we co-incubated germinating conidiospores in liquid SNA medium with either Cy3-labelled 250 nt long dsRNA derived from the *GFP* gene or Alexa Fluor® 555-labeled siRNA in the dark. *B. cinerea* was used as a positive control in these experiments as this fungal species is known to be engaging in bidirectional cross-kingdom RNAi and readily take up external dsRNAs (Wang *et al.*, 2016). Germinated spores were monitored using fluorescent microscopy and, as expected, fluorescent signals were detected in the *B. cinerea* from 12 h of co-incubation with labelled dsRNA species onwards (Figure 8). However, no fluorescent signals were detected in germinated *Z. tritici* conidiospores, which clearly extended hyphal strands, even by 48 h post co-incubation with either long or short labelled dsRNA species (Figure 8). Thus, it appears that *Z. tricici* lacks the capacity to uptake exogenous dsRNAs.

**Figure 8.**
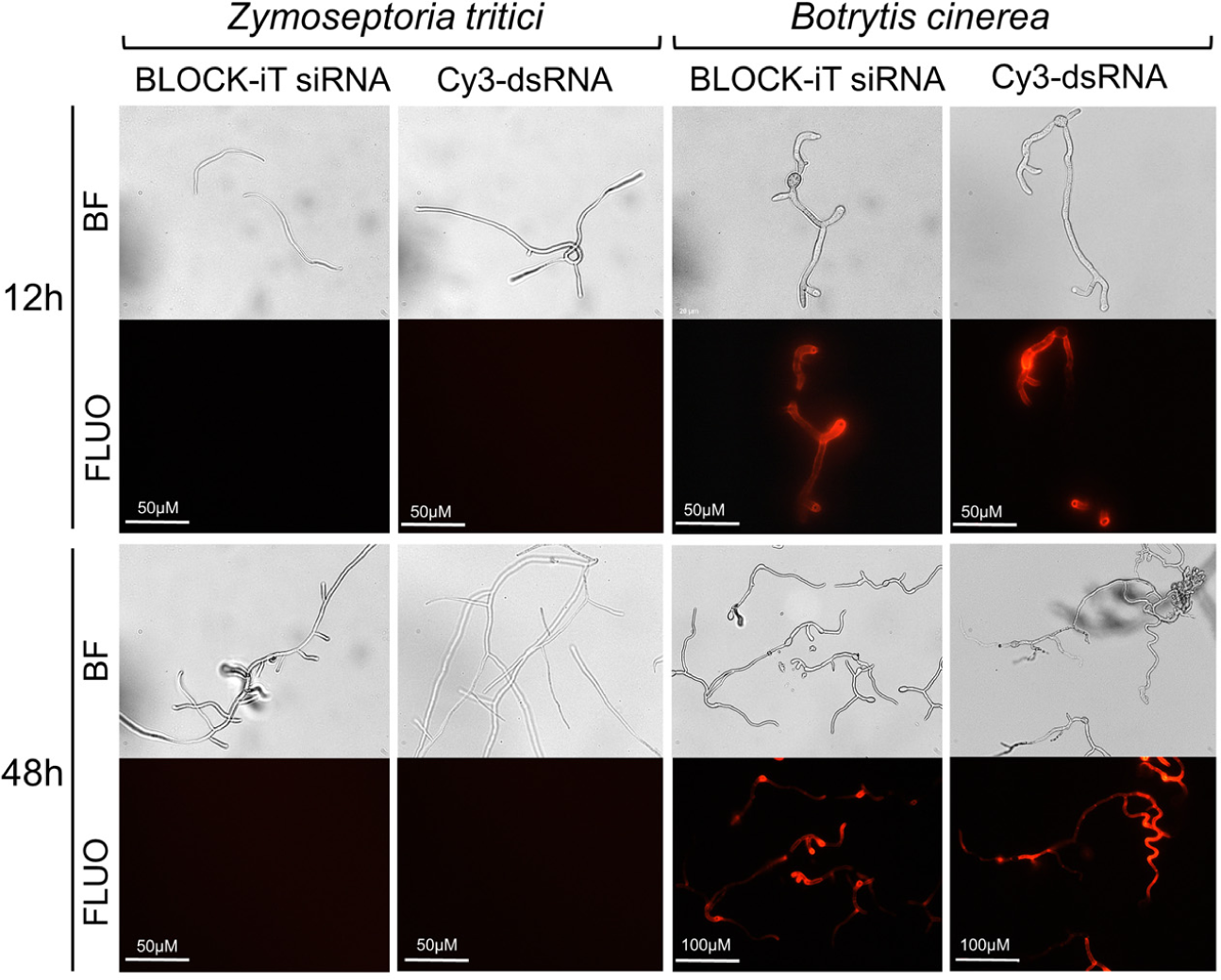
Assessment of dsRNA molecules uptake in *Zymoseptoria tritici* and *Botrytis cinerea*. Uptake of fluorescently labelled short dsRNA (Alexa Fluor Red BLOCK-iT siRNA) or 250-long long Cy3-labelled dsRNA against GFP was monitored at 12 h and 48 h of co-incubation using microscopy. Images are depicted as captured with bright field (BF) and fluorescent (FLUO) absorption 555 nm, emission 565 nm) settings.

## 4 Discussions

This study represents the most extensive attempt to date to characterize the sRNA transcriptome of the foliar wheat pathogen *Z. tritici* IPO323, with 389 sRNA loci predicted to be active at one or more timepoints during infection of a susceptible wheat cv. Bobwhite (Figure 1; Table S3). Furthermore, here we examined the possible role of cross-kingdom sRNA transfer and gene silencing in the interaction between this fungus and its natural host (wheat) and assessed the potential of RNAi as a possible future strategy for control of Septoria tritici blotch disease in wheat.

The contribution of the fungal RNAi pathway to virulence was assessed through production of targeted deletions of the *ZtDCL*, *ZtAGO1* and *ZtAGO2* genes. Fungal mutants for each of these three genes exhibited wild-type levels of virulence on wheat suggesting that these important components of the canonical RNAi pathway in *Z. tritici* are dispensable for successful wheat infection. These mutants also displayed no abnormal yeast-like growth morphology when the fungus was cultured on the rich growth media such as YPD (data not shown). Together, this implied that the *Z. tritici* RNAi pathway is superfluous or plays only a minimal role during either yeast-like or filamentous growth of this species. However, subsequent experiments revealed that expression of some fungal sRNAs is retained in the *Z. tritici Δdcl* mutant, at the level similar to that in the wild-type fungus (Figure 5). Given that this was observed for three of four examined sRNAs of interest (Figure 5), this data provided preliminary evidence that a proportion of the total sRNA pool in *Z. tritici* may be generated in a *ZtDCL*-independent manner. It would be interesting to investigate this further by determining a complete repertoire of sRNAs produced independently of the canonical RNAi pathway for example through sRNA sequencing from *in vitro* and *in planta* samples infected with the *Z. tritici Δdcl* mutant. It would also be of interest to further investigate the components and mechanisms of a *ZtDCL*-independent sRNA biogenesis in *Z. tritici* and compare to those that have previously been reported for other fungal species (Chang *et al.*, 2012). Simultaneous inactivation of both, this as yet uncharacterized pathway and a canonical Dicer-dependent sRNA biogenesis pathway, via deletion of key genes involved in both pathways in the same strain will be necessary to definitively conclude the involvement (or lack of) of sRNAs in successful infection by this fungus.

Two hundred and thirty-four *Z. tritici* sRNA loci were predicted to generate sRNAs putatively targeting 737 wheat transcripts with annotations for some suggestive of their potential role in plant immunity (Table S5), implying the existence of cross-kingdom RNAi in this pathosystem. It might be expected that the abundance of candidate wheat mRNA targets is reduced during *Z. tritici* infection due to the fungal sRNA-mediated cleavage. However, less than one third of the predicted targets showed downregulation of more than 2-fold even at one time point during *Z. tritici* IPO323 infection of a susceptible wheat cv. Riband, and downregulation of only ∼ 70 of these transcripts was found to be statistically significant. Moreover, we were unable to detect the predicted ZtsRNA1 - ZtsRNA4 catalyzed cleavage fragments in any of their 10 predicted wheat mRNA targets using 5’-RACE (Figure S7). It is possible that this could be because the predicted wheat mRNA – *Z. tritici* sRNA interactions result in translational repression rather than transcript cleavage. It may also be possible that the analyzed wheat mRNA targets identified computationally are either not genuine targets, or not sufficiently well-targeted for the effects to be detectable using the methods described here.

The rational for this investigation was based on recent findings describing bidirectional cross-kingdom RNAi for the necrotrophic fungus *B. cinerea* (Wang *et al.*, 2016). The data presented here suggest that for *Z. tritici* fungal RNAi pathways do not play a significant role in the successful colonization of wheat plants. This agrees well with the data from a complementary study by Ma *et al.* (2018) in which no strong evidence for silencing of wheat genes in the susceptible cv. Drifter by sRNAs of the *Z. tritici* isolate 3D7 was found. It is likely that evasion or manipulation of host immunity by this fungal species is more reliant on previously described mechanism such as the deployment of proteinaceous effectors and /or the production of specific metabolites (Deller *et al.*, 2011; Kettles & Kanyuka, 2016). Whilst we conclude that fungal RNAi plays a minimal role during infection, the genome of *Z. tritici* does encode key components of the canonical RNAi pathway such as DCL, AGOs and RdRps. Therefore, it seemed possible that the plant-derived or exogenously applied long dsRNAs or sRNAs if taken up by *Z. tritici* could induce RNAi in this fungus. However, there were no reductions in the amount of *Z. tritici* infection in the BSMV-HIGS experiments targeting fungal genes essential for life or pathogenicity towards wheat (Figure 6) and also no observable effects on growth of *Z. tritici* when germinating fungal conidiospores were co-incubated with the fungal-derived dsRNA targeting essential genes such as *ZtCYP51*, *ZtTUBa* and *ZtTUBb* (Figure 7). These data could be explained by the observations that *Z. tritici* is incapable or very inefficient in taking up exogenous dsRNAs (Figure 8). We therefore conclude that the prospects for topical chemical intervention via next-generation RNA fungicides or through the generation of the stable transgenic HIGS lines in wheat targeting *Z. tritici* genes that are essential for life appear doubtful. We also hypothesize that the RNAi machinery in *Z. tritici* may either be completely redundant or play a role perhaps during growth under certain stressful conditions e.g. fungicide treatment or only during specific developmental stages. This wouldn’t be unprecedented as some fungal species, including *Ustilago maydis,* which is a serious pathogen of maize, have lost RNAi (Billmyre *et al.*, 2013). Furthermore, knockout of either DCL, AGO, or RdRp encoding genes in another fungal pathogen of wheat, *F. graminearum*, did not compromise pathogenicity towards wheat (Kim *et al.*, 2015; and own unpublished data) and it has been recently demonstrated that the RNAi machinery in *F. graminearum* is mainly involved in the regulation of sexual development (Kim *et al.*, 2015; Zeng *et al.*, 2018). It would be interesting to investigate in future studies whether the same may be true for *Z. tritici*. Also, it seems possible that the RNAi machinery in this fungal species may play a role in defense against mycoviruses. A mycovirus in several *Z. tritici* isolates has been previously reported and shown to be highly expressed under various stress conditions (Kema *et al.*, 2008).

## 5 Author Contributions

The KK, KHK, GS, and MC conceived the project. KK generated material for sRNAseq and re-analyzed the previously published RNAseq data set. sRNAseq analysis and prediction of wheat mRNA targets was done by PH. GK and CB performed experimental validation of fungal sRNA and wheat mRNA expression and developed and characterized *Z. tritici* gene deletion mutants. DB and GS assessed capacity of *Z. tritici* to uptake dsRNAs. JR and DB advised on various aspects of *Z. tritici* biology and RNAi, respectively. BH performed BSMV-HIGS and the *in vitro* RNAi experimentation. GK, BH, KHK and KK analyzed the data. GK and KK wrote the manuscript.

## 7 Acknowledgments

The authors would like to thank Rothamsted Horticultural Services for excellent plant care, and Stephanie Widdison (Syngenta) for constructive comments upon the text, tables and figures.

## 8 Data Availability Statement

The sRNA sequencing data was deposited to the European Nucleotide Archive https://www.ebi.ac.uk/ena under the study PRJEB28454.

## 11 Supporting Information

**Table S1.** Primers used in this study.

**Table S2.** Expression levels of the key predicted RNAi machinery components in *Zymoseptoria tritici* isolate IPO323.

**Table S3.** Computationally predicted *Zymoseptoria tritici* IPO323 sRNA loci.

**Table S4.** *Zymoseptoria tritici* IPO323 sRNA loci and mature sRNAs predicted to target wheat transcripts.

**Table S5.** Wheat transcripts predicted to be targeted by *Zymoseptoria tritici* IPO323 mature sRNAs.

**Table S6.** *Zymoseptoria tritici* (Zt) genes selected for gene silencing using BSMV-HIGS.

**Methods S1.** Small RNA loci likelihood estimation.

**Figure S1.** Domain architecture of candidate AGO proteins in *Zymoseptoria tritici* isolate IPO323.

**Figure S2.** Examples of the distribution of sRNAseq reads mapped to the *Zymoseptoria tritici* IPO323 genome from *in vitro* culture and infected wheat cv. Bobwhite samples at 13 days post inoculation (dpi).

**Figure S3.** Origin, length and expression levels of 389 sRNA loci identified to be active in *Zymoseptoria tritici* isolate IPO323 during wheat infection.

**Figure S4.** Numbers of sRNA loci residing on individual chromosomes of *Zymoseptoria tritici* isolate IPO323. Core and accessory chromosomes are shown in blue and orange, respectively.

**Figure S5.** Size distribution and percentage of nucleotides in the 5’ position of 262 mature *Zymoseptoria tritici* sRNAs predicted to target wheat transcripts.

**Figure S6.** Expression of wheat mRNA Traes_4BS_5E12F0B27 in leaf tissue at 4, 9, and 13 days post inoculation (dpi) with RNAi-competent and RNAi-deficient *Zymoseptoria tritici* strains. Bars indicate SE.

**Figure S7.** 5’-RACE on the selected wheat targets. Assays were performed using both mock-infected (-) and *Zymoseptoria tritici* IPO323-infected (+) wheat cDNA as templates. Reaction products from lanes indicated in red were sequenced.

**Figure S8.** Leaves of wheat cv. Riband plants at 14 days post pre-treatment with the BSMV:asTaMgChlH construct showing RNAi-induced chlorophyll deficiency (orangey-yellow coloration).

## References

Billmyre RB, Calo S, Feretzaki M, Wang X, Heitman J. 2013. RNAi function, diversity, and loss in the fungal kingdom. Chromosome Research 21: 561–572.

Bowler J, Scott E, Tailor R, Scalliet G, Ray J, Csukai M. 2010. New capabilities for *Mycosphaerella graminicola* research. Molecular Plant Pathology 11: 691–704.

Cai Q, Qiao L, Wang M, He B, Lin F-M, Palmquist J, Huang S-D, Jin H. 2018. Plants send small RNAs in extracellular vesicles to fungal pathogen to silence virulence genes. Science 360: 1126–1129.

Chang S-S, Zhang Z, Liu Y. 2012. RNA interference pathways in fungi: mechanisms and functions. Annual Review of Microbiology 66: 305–323.

Chen W, Kastner C, Nowara D, Oliveira-Garcia E, Rutten T, Zhao Y, Deising HB, Kumlehn J, Schweizer P. 2016. Host-induced silencing of *Fusarium culmorum* genes protects wheat from infection. Journal of Experimental Botany 67: 4979–4991.

Cheng W, Song X-S, Li H-P, Cao L-H, Sun K, Qiu X-L, Xu Y-B, Yang P, Huang T, Zhang J-B, et al. 2015. Host-induced gene silencing of an essential chitin synthase gene confers durable resistance to Fusarium head blight and seedling blight in wheat. Plant Biotechnology Journal 13: 1335–1345.

Dean R, Van Kan Jall, Pretorius ZA, Hammond-Kosack KE, Di Pietro A, Spanu PD, Rudd JJ, Dickman M, Kahmann R, Ellis J, et al. 2012. The Top 10 fungal pathogens in molecular plant pathology. Molecular Plant Pathology 13: 414–30.

Deller S, Hammond-Kosack KE, Rudd JJ. 2011. The complex interactions between host immunity and non-biotrophic fungal pathogens of wheat leaves. Journal of Plant Physiology 168: 63–71.

Ellendorff U, Fradin EF, de Jonge R, Thomma Bphj. 2009. RNA silencing is required for Arabidopsis defence against Verticillium wilt disease. Journal of Experimental Botany 60: 591–602.

Fujii H, Chiou T-J, Lin S-I, Aung K, Zhu J-K. 2005. A miRNA involved in phosphate-starvation response in Arabidopsis. Current Biology 15: 2038–2043.

Gasciolli V, Mallory AC, Bartel DP, Vaucheret H. 2005. Partially redundant functions of Arabidopsis DICER-like enzymes and a role for DCL4 in producing trans-acting siRNAs. Current Biology 15: 1494–1500.

Ghag SB, Shekhawat UKS, Ganapathi TR. 2014. Host-induced post-transcriptional hairpin RNA-mediated gene silencing of vital fungal genes confers efficient resistance against Fusarium wilt in banana. Plant Biotechnology Journal 12: 541–53.

Ghildiyal M, Zamore PD. 2009. Small silencing RNAs: an expanding universe. Nature Reviews. Genetics 10: 94–108.

Goodwin SB, M’barek S Ben, Dhillon B, Wittenberg AHJ, Crane CF, Hane JK, Foster AJ, Van der Lee TAJ, Grimwood J, Aerts A, et al. 2011. Finished genome of the fungal wheat pathogen *Mycosphaerella graminicola* reveals dispensome structure, chromosome plasticity, and stealth pathogenesis. PLoS Genetics 7: e1002070. doi: 10.1371/journal.pgen.1002070

Hamilton AJ, Baulcombe DC. 1999. A species of small antisense RNA in posttranscriptional gene silencing in plants. Science 286: 950–952.

Haupt S, Duncan GH, Holzberg S, Oparka KJ. 2001. Evidence for symplastic phloem unloading in sink leaves of barley. Plant Physiology 125: 209–218.

Hollomon DW, Butters JA, Barker H, Hall L. 1998. Fungal beta-tubulin, expressed as a fusion protein, binds benzimidazole and phenylcarbamate fungicides. Antimicrobial Agents and Chemotherapy 42: 2171–2173.

Kema GHJ, van der Lee TAJ, Mendes O, Verstappen ECP, Lankhorst RK, Sandbrink H, van der Burgt A, Zwiers LH, Csukai M, Waalwijk C. 2008. Large-scale gene discovery in the septoria tritici blotch fungus *Mycosphaerella graminicola* with a focus on in planta expression. Molecular Plant-Microbe Interactions 21: 1249–1260.

Kema GHJ, Mirzadi Gohari A, Aouini L, Gibriel HAY, Ware SB, van den Bosch F, Manning-Smith R, Alonso-Chavez V, Helps J, Ben M’Barek S, et al. 2018. Stress and sexual reproduction affect the dynamics of the wheat pathogen effector AvrStb6 and strobilurin resistance. Nature Genetics 50: 375–380.

Kema GHJ, Yu D, Rijkenberg FHJ, Shaw MW, Baayen RP. 1996. Histology of the pathogenesis of *Mycosphaerella graminicola* in wheat. Phytopathology 86: 777–786.

Keon J, Antoniw J, Carzaniga R, Deller S, Ward JL, Baker JM, Beale MH, Hammond-Kosack K, Rudd JJ. 2007. Transcriptional adaptation of *Mycosphaerella graminicola* to programmed cell death (PCD) of its susceptible wheat host. Molecular Plant-Microbe Interactions 20: 178–193.

Kettles GJ, Bayon C, Canning G, Rudd JJ, Kanyuka K. 2017. Apoplastic recognition of multiple candidate effectors from the wheat pathogen *Zymoseptoria tritici* in the nonhost plant *Nicotiana benthamiana*. New Phytologist 213: 338–350.

Kettles GJ, Drurey C, Schoonbeek H, Maule AJ, Hogenhout SA. 2013. Resistance of *Arabidopsis thaliana* to the green peach aphid, *Myzus persicae*, involves camalexin and is regulated by microRNAs. New Phytologist 198: 1178–1190.

Kettles GJ, Kanyuka K. 2016. Dissecting the molecular interactions between wheat and the fungal pathogen *Zymoseptoria tritici*. Frontiers in Plant Science 7: 508. doi: 10.3389/fpls.2016.00508

Kim H-K, Jo S-M, Kim G-Y, Kim D-W, Kim Y-K, Yun S-H. 2015. A large-scale functional analysis of putative target genes of mating-type loci provides insight into the regulation of sexual development of the cereal pathogen *Fusarium graminearum*. PLoS Genetics 11: e1005486. doi: 10.1371/journal.pgen.1005486

King R, Urban M, Lauder RP, Hawkins N, Evans M, Plummer A, Halsey K, Lovegrove A, Hammond-Kosack K, Rudd JJ. 2017. A conserved fungal glycosyltransferase facilitates pathogenesis of plants by enabling hyphal growth on solid surfaces. PLoS Pathogens 13: e1006672. doi: 10.1371/journal.ppat.1006672

Koch A, Biedenkopf D, Furch A, Weber L, Rossbach O, Abdellatef E, Linicus L, Johannsmeier J, Jelonek L, Goesmann A, et al. 2016. An RNAi-based control of *Fusarium graminearum* infections through spraying of long dsRNAs involves a plant passage and is controlled by the fungal silencing machinery. PLoS pathogens 12: e1005901. doi: 10.1371/journal.ppat.1005901

Koch A, Kumar N, Weber L, Keller H, Imani J, Kogel K-H. 2013. Host-induced gene silencing of cytochrome P450 lanosterol C14α-demethylase-encoding genes confers strong resistance to Fusarium species. Proceedings of the National Academy of Sciences of the United States of America 110: 19324–19329.

Langmead B, Salzberg SL. 2012. Fast gapped-read alignment with Bowtie 2. Nature Methods 9: 357–359.

Lee W-S, Devonshire BJ, Hammond-Kosack KE, Rudd JJ, Kanyuka K. 2015a. Deregulation of plant cell death through disruption of chloroplast functionality affects asexual sporulation of *Zymoseptoria tritici* on wheat. Molecular Plant-Microbe Interactions 28: 590–604.

Lee W-S, Hammond-Kosack KE, Kanyuka K. 2012. Barley stripe mosaic virus-mediated tools for investigating gene function in cereal plants and their pathogens: Virus-induced gene silencing, Host-mediated gene silencing, and Virus-mediated overexpression of heterologous protein. Plant Physiology 160: 582–590.

Lee W-S, Rudd JJ, Kanyuka K. 2015b. Virus induced gene silencing (VIGS) for functional analysis of wheat genes involved in *Zymoseptoria tritici* susceptibility and resistance. Fungal Genetics and Biology 79: 84–88.

Li H, Handsaker B, Wysoker A, Fennell T, Ruan J, Homer N, Marth G, Abecasis G, Durbin R. 2009. The Sequence Alignment/Map format and SAMtools. Bioinformatics 25: 2078–2079.

Liu J, Cheng X, Liu P, Sun J. 2017. miR156-targeted SBP-box transcription factors interact with DWARF53 to regulate *TEOSINTE BRANCHED*1 and *BARREN STALK1* expression in bread wheat. Plant Physiology 174: 1931–1948.

Llave C, Xie Z, Kasschau KD, Carrington JC. 2002. Cleavage of Scarecrow-like mRNA targets directed by a class of Arabidopsis miRNA. Science 297: 2053–2056.

Lohse M, Nagel A, Herter T, May P, Schroda M, Zrenner R, Tohge T, Fernie AR, Stitt M, Usadel B. 2014. Mercator: a fast and simple web server for genome scale functional annotation of plant sequence data. Plant, Cell & Environment 37: 1250–1258.

Ma X, Bologna N, Palma-Guerrero J. 2018. Small RNA bidirectional crosstalk during the interaction between wheat and *Zymoseptoria tritici*. *BioRxiv*, submitted.

Machado AK, Brown NA, Urban M, Kanyuka K, Hammond-Kosack KE. 2018. RNAi as an emerging approach to control Fusarium head blight disease and mycotoxin contamination in cereals. Pest Management Science 74: 790–799.

Marshall R, Kombrink A, Motteram J, Loza-Reyes E, Lucas J, Hammond-Kosack KE, Thomma BPHJ, Rudd JJ. 2011. Analysis of two in planta expressed LysM effector homologs from the fungus *Mycosphaerella graminicola* reveals novel functional properties and varying contributions to virulence on wheat. Plant Physiology 156: 756–769.

Matzke M, Kanno T, Huettel B, Daxinger L, Matzke AJM. 2007. Targets of RNA-directed DNA methylation. Current Opinion in Plant Biology 10: 512–519.

McLoughlin AG, Wytinck N, Walker PL, Girard IJ, Rashid KY, de Kievit T, Fernando WGD, Whyard S, Belmonte MF. 2018. Identification and application of exogenous dsRNA confers plant protection against *Sclerotinia sclerotiorum* and *Botrytis cinerea*. Scientific Reports 8: 7320. doi: 10.1038/s41598-018-25434-4

Meile L, Croll D, Brunner PC, Plissonneau C, Hartmann FE, McDonald BA, Sánchez-Vallet A. 2018. A fungal avirulence factor encoded in a highly plastic genomic region triggers partial resistance to septoria tritici blotch. New Phytologist 219: 1048–1061.

Motteram J, Küfner I, Deller S, Brunner F, Hammond-Kosack KE, Nürnberger T, Rudd JJ. 2009. Molecular characterization and functional analysis of MgNLP, the sole NPP1 domain–containing protein, from the fungal wheat leaf pathogen *Mycosphaerella graminicola*. Molecular Plant-Microbe Interactions 22: 790–799.

Motteram J, Lovegrove A, Pirie E, Marsh J, Devonshire J, van de Meene A, Hammond-Kosack K, Rudd JJ. 2011. Aberrant protein N-glycosylation impacts upon infection-related growth transitions of the haploid plant-pathogenic fungus *Mycosphaerella graminicola*. Molecular Microbiology 81: 415–433.

Navarro L, Dunoyer P, Jay F, Arnold B, Dharmasiri N, Estelle M, Voinnet O, Jones JDG. 2006. A plant miRNA contributes to antibacterial resistance by repressing auxin signaling. Science 312: 436–439.

Nicolás FE, Torres-Martínez S, Ruiz-Vázquez RM. 2013. Loss and retention of RNA interference in fungi and parasites. PLoS Pathogens 9: e1003089. doi: 10.1371/journal.ppat.1003089

Nolan T, Braccini L, Azzalin G, De Toni A, Macino G, Cogoni C. 2005. The post-transcriptional gene silencing machinery functions independently of DNA methylation to repress a LINE1-like retrotransposon in *Neurospora crassa*. Nucleic Acids Research 33: 1564–1573.

Nowara D, Gay A, Lacomme C, Shaw J, Ridout C, Douchkov D, Hensel G, Kumlehn J, Schweizer P. 2010. HIGS: host-induced gene silencing in the obligate biotrophic fungal pathogen Blumeria graminis. Plant Cell 22: 3130–3141.

Pandey SP, Baldwin IT. 2007. RNA-directed RNA polymerase 1 (RdR1) mediates the resistance of *Nicotiana attenuata* to herbivore attack in nature. Plant Journal 50: 40–53.

Qi T, Zhu X, Tan C, Liu P, Guo J, Kang Z, Guo J. 2018. Host-induced gene silencing of an important pathogenicity factor *PsCPK1* in *Puccinia striiformis* f. sp. tritici enhances resistance of wheat to stripe rust. Plant Biotechnology Journal 16: 797–807.

Roberts A, Pachter L. 2013. Streaming fragment assignment for real-time analysis of sequencing experiments. Nature Methods 10: 71–73.

Robinson MD, McCarthy DJ, Smyth GK. 2010. edgeR: a Bioconductor package for differential expression analysis of digital gene expression data. Bioinformatics 26: 139–140.

Robinson MD, Oshlack A. 2010. A scaling normalization method for differential expression analysis of RNA-seq data. Genome Biology 11: R25. doi: 10.1186/gb-2010-11-3-r25

Rohel EA, Payne AC, Fraaije BA, Hollomon DW. 2001. Exploring infection of wheat and carbohydrate metabolism in *Mycosphaerella graminicola* transformants with differentially regulated green fluorescent protein expression. Molecular Plant-Microbe Interactions 14: 156–163.

Rudd JJ, Kanyuka K, Hassani-Pak K, Derbyshire M, Andongabo A, Devonshire J, Lysenko A, Saqi M, Desai NM, Powers SJ, et al. 2015. Transcriptome and metabolite profiling of the infection cycle of *Zymoseptoria tritici* on wheat reveals a biphasic interaction with plant immunity involving differential pathogen chromosomal contributions and a variation on the hemibiotrophic lifestyle definition. Plant Physiology 167: 1158–1185.

Saintenac C, Lee W-S, Cambon F, Rudd JJ, King RC, Marande W, Powers SJ, Bergès H, Phillips AL, Uauy C, et al. 2018. Wheat receptor-kinase-like protein Stb6 controls gene-for-gene resistance to fungal pathogen *Zymoseptoria tritici*. Nature Genetics 50: 368–374.

Segers GC, Zhang X, Deng F, Sun Q, Nuss DL. 2007. Evidence that RNA silencing functions as an antiviral defense mechanism in fungi. Proceedings of the National Academy of Sciences of the United States of America 104: 12902–12906.

Siegel MR. 1981. Sterol-inhibiting fungicides: effects on sterol biosynthesis and sites of action. Plant Disease 65: 986–989.

Song Y, Thomma BPHJ. 2018. Host-induced gene silencing compromises Verticillium wilt in tomato and Arabidopsis. Molecular Plant Pathology 19: 77–89.

Sunkar R, Chinnusamy V, Zhu J, Zhu J-K. 2007. Small RNAs as big players in plant abiotic stress responses and nutrient deprivation. Trends in Plant Science 12: 301–309.

Varkonyi-Gasic E, Wu R, Wood M, Walton EF, Hellens RP. 2007. Protocol: a highly sensitive RT-PCR method for detection and quantification of microRNAs. Plant Methods 3: 12. doi: 10.1186/1746-4811-3-12

Vazquez F, Legrand S, Windels D. 2010. The biosynthetic pathways and biological scopes of plant small RNAs. Trends in Plant Science 15: 337–45.

Vickers KC, Roteta LA, Hucheson-Dilks H, Han L, Guo Y. 2015. Mining diverse small RNA species in the deep transcriptome. Trends in Biochemical Sciences 40: 4–7.

Volpe TA, Kidner C, Hall IM, Teng G, Grewal SIS, Martienssen RA. 2002. Regulation of heterochromatic silencing and histone H3 lysine-9 methylation by RNAi. Science 297: 1833–1837.

Wang M, Weiberg A, Lin F-M, Thomma BPHJ, Huang H-D, Jin H. 2016. Bidirectional cross-kingdom RNAi and fungal uptake of external RNAs confer plant protection. Nature Plants 2: 16151. doi: 10.1038/nplants.2016.151

Weiberg A, Wang M, Lin F-M, Zhao H, Zhang Z, Kaloshian I, Huang H-D, Jin H. 2013. Fungal small RNAs suppress plant immunity by hijacking host RNA interference pathways. Science 342: 118–123.

Xie Z, Johansen LK, Gustafson AM, Kasschau KD, Lellis AD, Zilberman D, Jacobsen SE, Carrington JC. 2004. Genetic and functional diversification of small RNA pathways in plants. PLoS Biology 2: E104. doi: 10.1371/journal.pbio.0020104

Yuan C, Li C, Yan L, Jackson AO, Liu Z, Han C, Yu J, Li D. 2011. A high throughput *Barley stripe mosaic virus* vector for virus induced gene silencing in monocots and dicots. PloS One 6: e26468. doi: 10.1371/journal.pone.0026468

Zeng W, Wang J, Wang Y, Lin J, Fu Y, Xie J, Jiang D, Chen T, Liu H, Cheng J. 2018. Dicer-like proteins regulate sexual development via the biogenesis of perithecium-specific microRNAs in a plant pathogenic fungus *Fusarium graminearum*. Frontiers in Microbiology 9: 818. doi: 10.3389/fmicb.2018.00818

Zhong Z, Marcel TC, Hartmann FE, Ma X, Plissonneau C, Zala M, Ducasse A, Confais J, Compain J, Lapalu N, et al. 2017. A small secreted protein in *Zymoseptoria tritici* is responsible for avirulence on wheat cultivars carrying the *Stb6* resistance gene. New Phytologist 214: 619–631.

