## Supplemental Material for "Analysis of small RNA silencing in *Zymoseptoria tritici* – wheat interactions"

**Figure S1.** Domain architecture of candidate AGO proteins in *Zymoseptoria tritici* isolate IPO323.

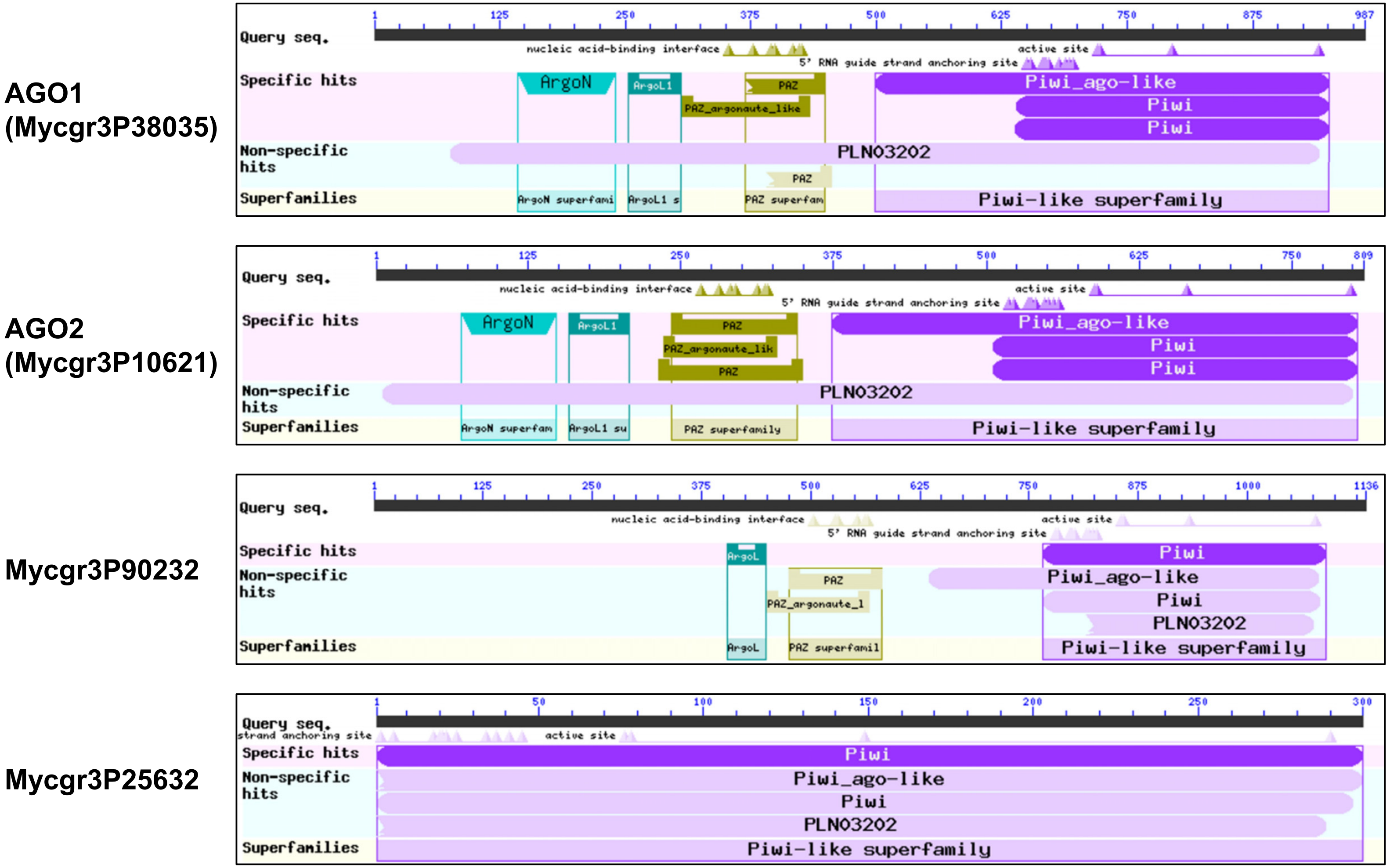

**Figure S2.** Examples of the distribution of sRNAseq reads mapped to the *Zymoseptoria tritici* IPO323 genome from *in vitro* culture and infected wheat cv. Bobwhite samples at 13 days post inoculation (dpi).

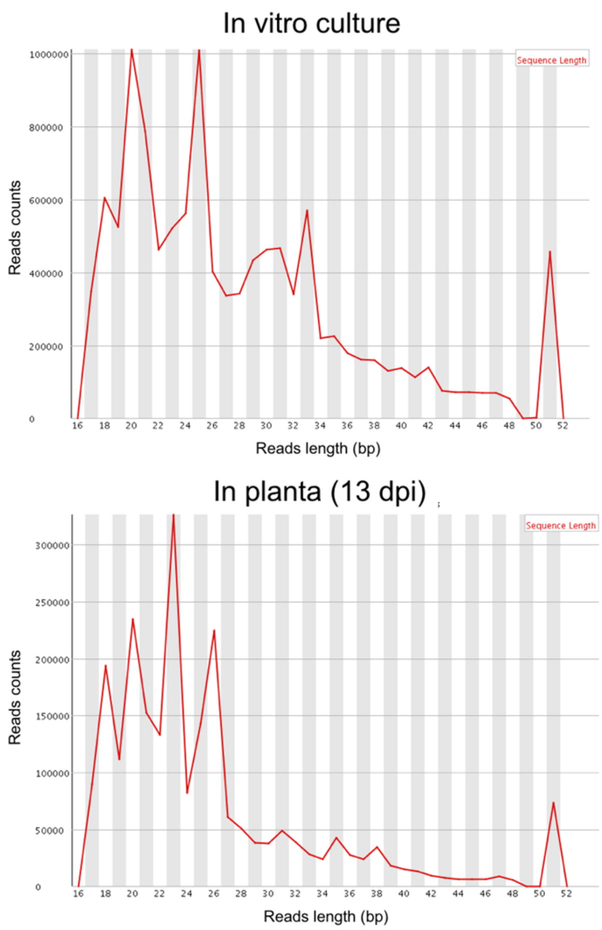

**Figure S3.** Origin, length and expression levels of 389 sRNA loci identified to be active in *Zymoseptoria tritici* isolate IPO323 during wheat infection.

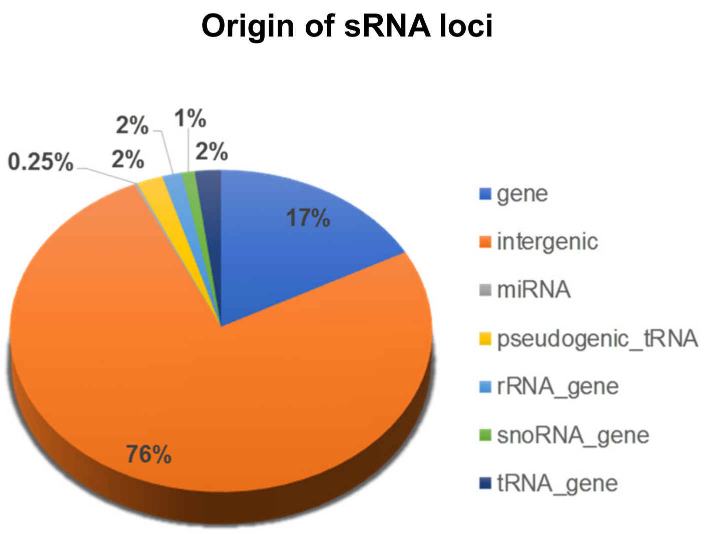


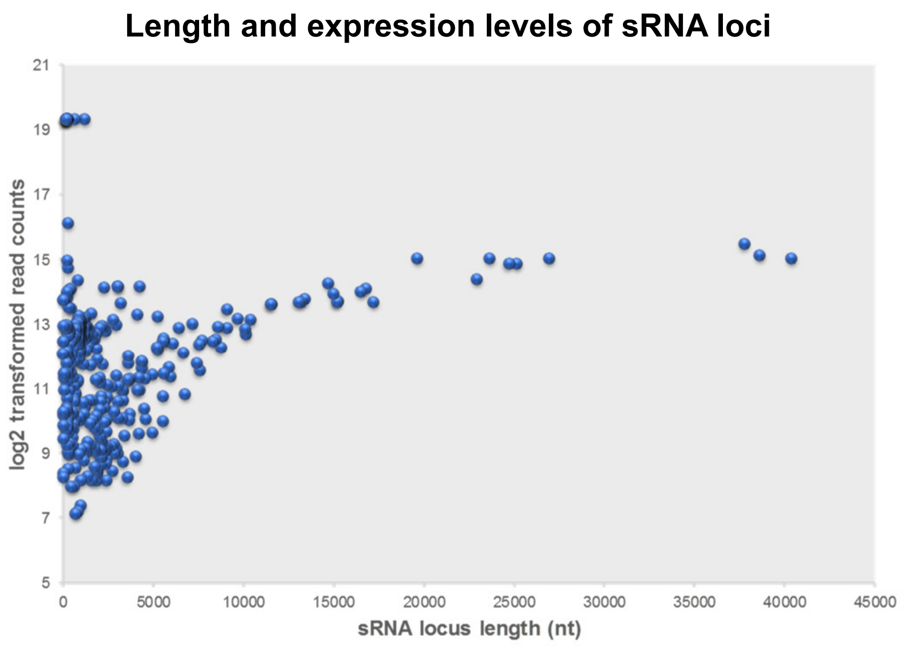

**Figure S4.** Numbers of sRNA loci residing on individual chromosomes of *Zymoseptoria tritici* isolate IPO323. Core and accessory chromosomes are shown in blue and orange, respectively.

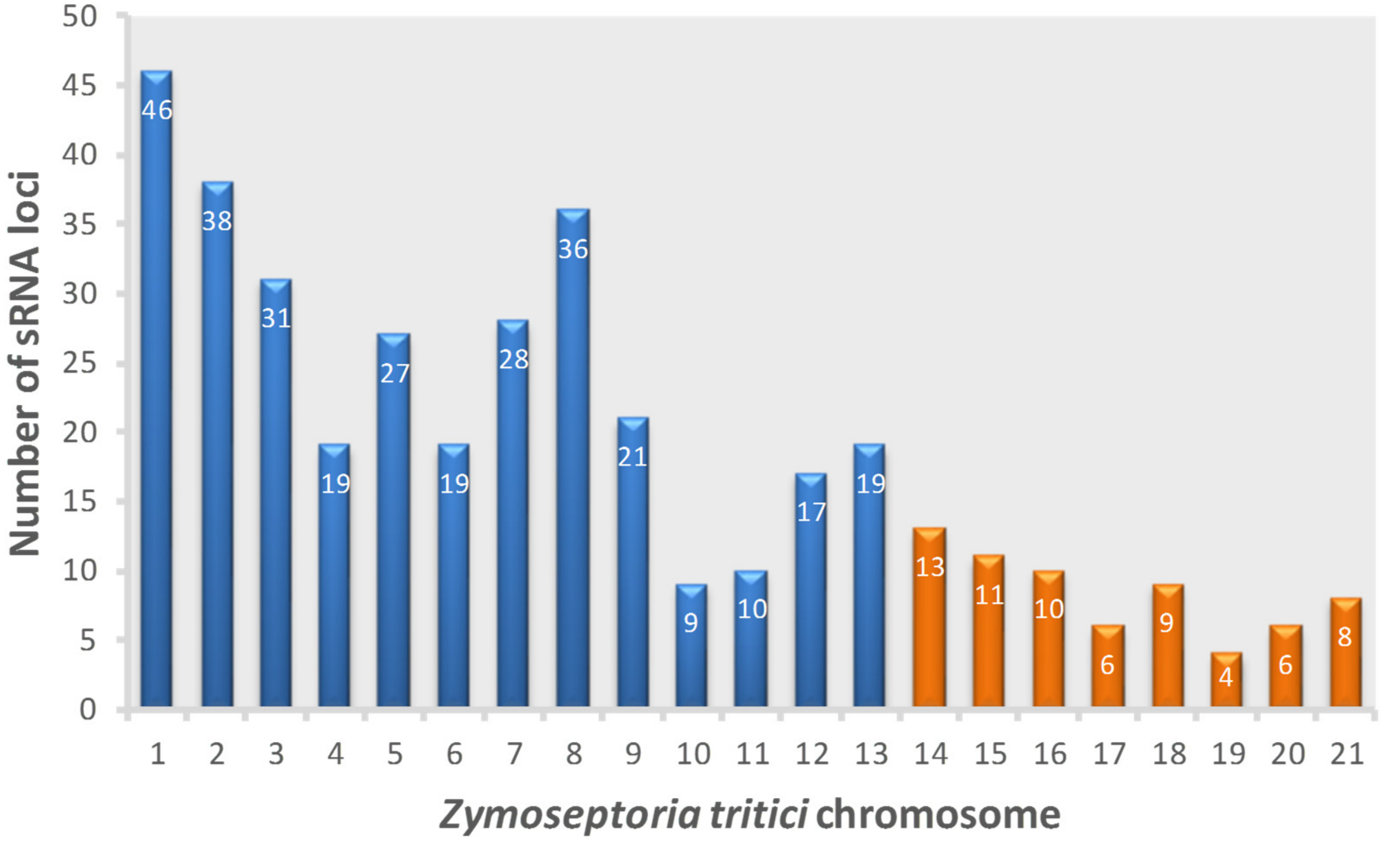

**Figure S5.** Size distribution and percentage of nucleotides in the 5' position of 262 mature *Zymoseptoria tritici* sRNAs predicted to target wheat transcripts.

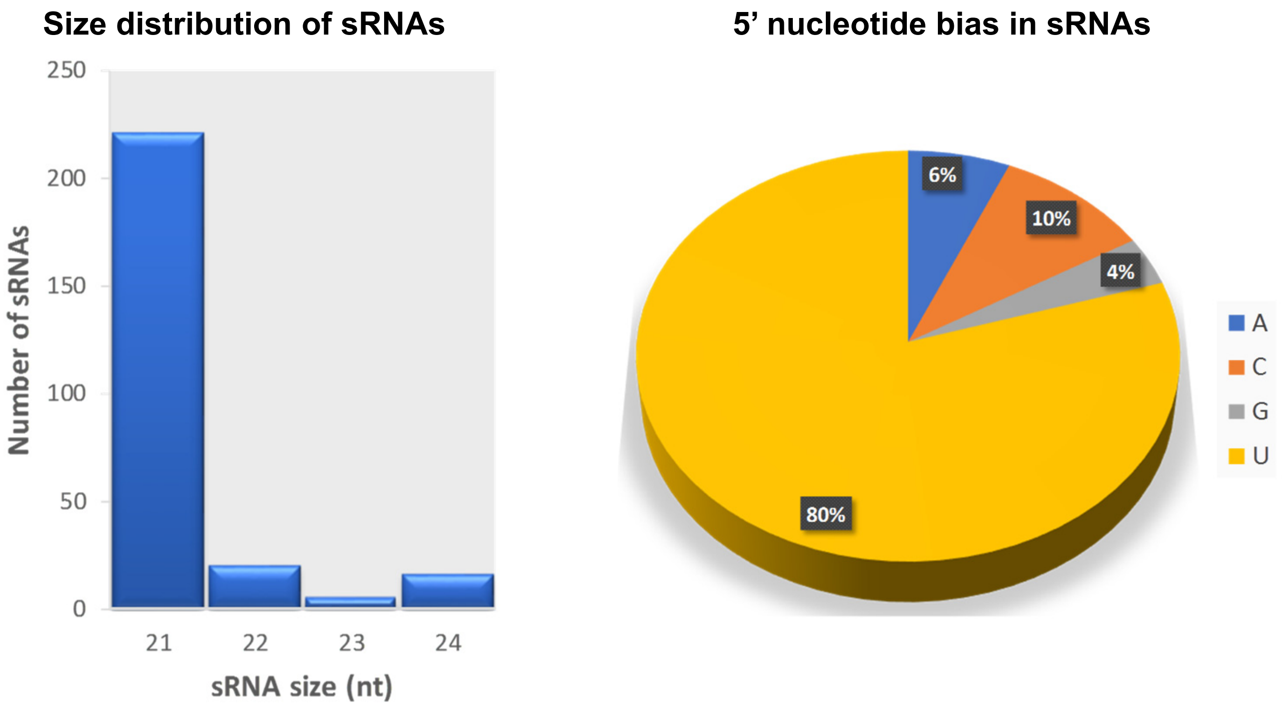

**Figure S6.** Expression of wheat mRNA Traes_4BS_5E12F0B27 in leaf tissue at 4, 9, and 13 days post inoculation (dpi) with RNAi-competent and RNAi-deficient *Zymoseptoria tritici* strains. Bars indicate SE.

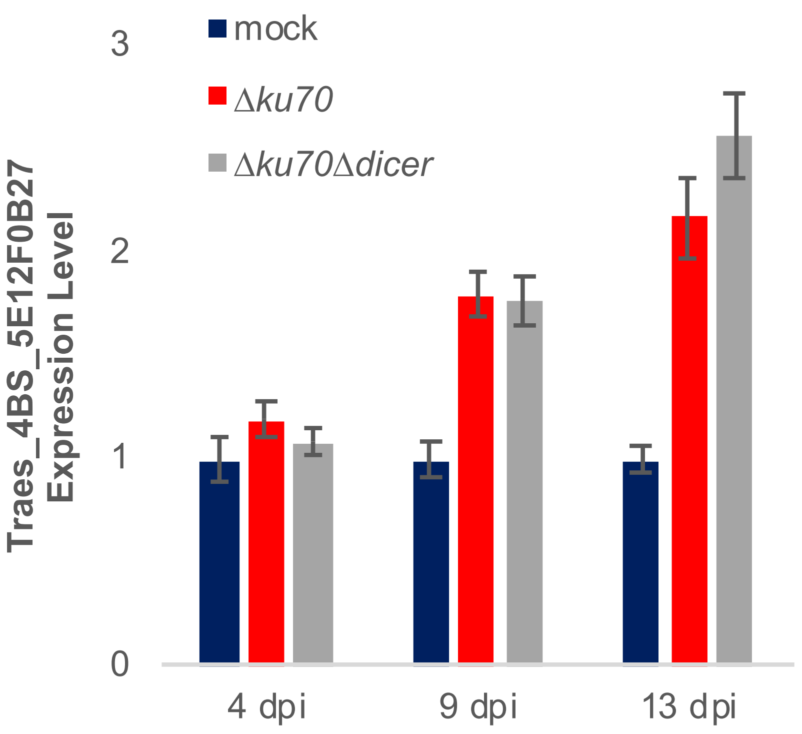

**Figure S7.** 5'-RACE on the selected wheat targets. Assays were performed using both mock-infected (-) and *Zymoseptoria tritici* IPO323-infected (+) wheat cDNA as templates. Reaction products from lanes indicated in red were sequenced.

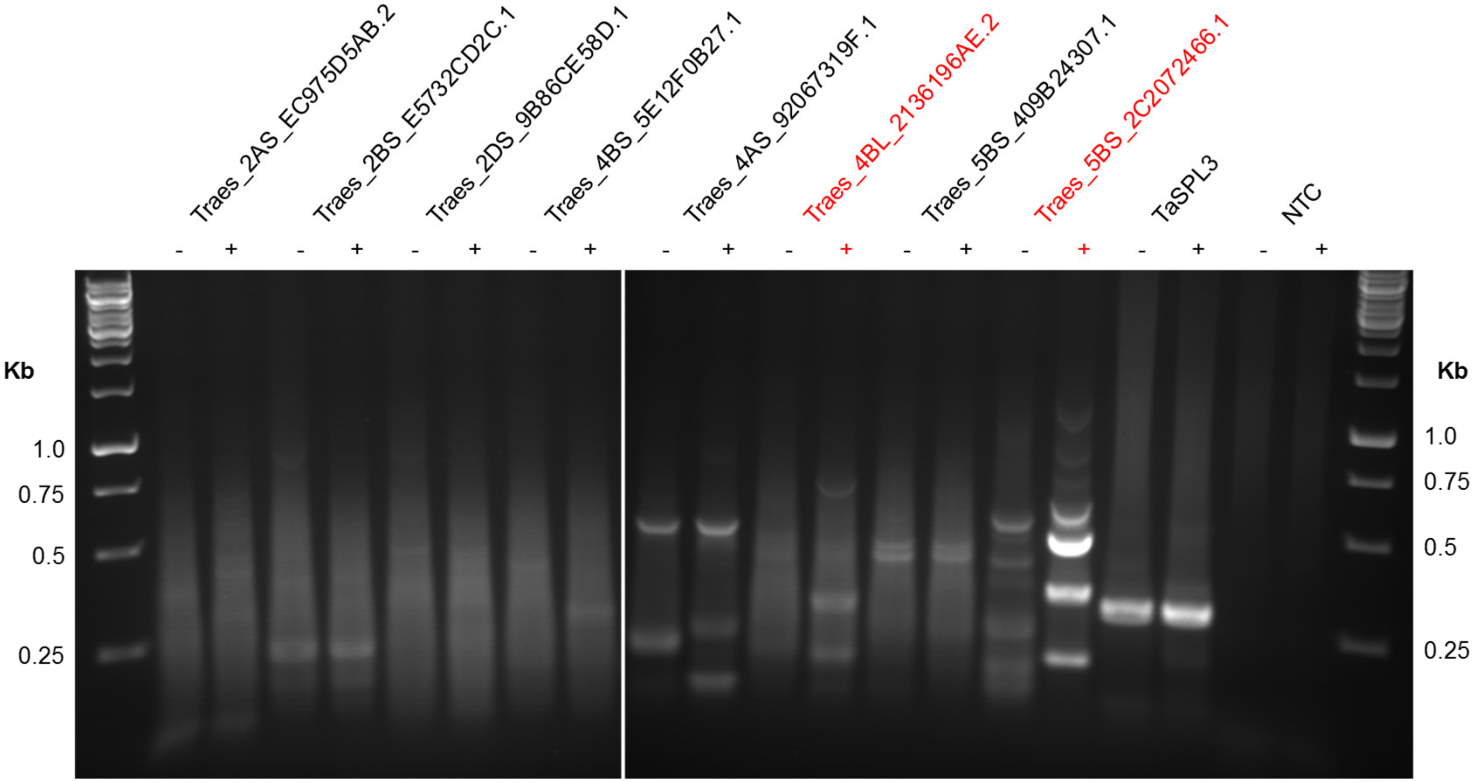

**Figure S8.** Leaves of wheat cv. Riband plants at 14 days post pre-treatment with the BSMV:asTaMgChlH construct showing RNAi-induced chlorophyll deficiency (orangey-yellow coloration).

Leaves from individual plants pre-treated with BSMV:asTaMgChlH

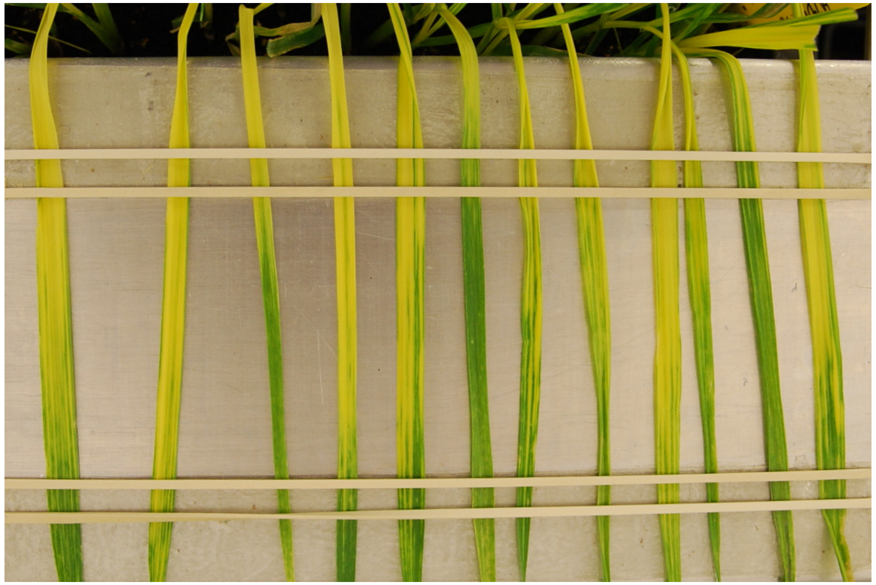

14 days post
virus inoculation

**Methods S1:** Small RNA locus likelihood estimation.

Small RNA (sRNA) locus likelihood estimation is to provide a posterior probability value indicating the probability of a given locus to be a true small RNA locus. The posterior probability is based on the sRNA sequencing reads coverage depth and read length distribution information for a given locus. A true sRNA locus is assumed to have higher coverage than a null locus does, which may have some short RNA sequences aligned that originate for example from degradation of mRNA. In addition, sRNA locus usually has a dominant length reads, e.g. 21 nt or 24 nt reads are highly dominated in a sRNA locus compared to other lengths reads. On the other hand, a locus with random, degraded mRNA products usually displays a more uniform distribution of all types short length reads. The following method to estimate sRNA locus probability is based on the above two assumptions.

**Bayesian model for sRNA locus prediction**

Let *I_l~B(1, p), l=1,…L*, be the Bernoulli random variable indicating whether a locus *l* is a sRNA locus, *Prob(I_l)=p*, where:

$$I_l = \begin{cases} 0 & if\ I_l = 0 \\ 1 & if\ I_l = 1 \end{cases} \qquad (1)$$

Then the posterior probability for a sRNA locus is:

$$\Pr(I_l = 1 \mid data) = \frac{p \Pr(data \mid I_l = 1)}{p \Pr(data \mid I_l = 1) + (1-p)\Pr(data \mid I_l = 0)} \qquad (2)$$

The *Prob(data|I_l=1)* is based on both read coverage depth and read length distribution, therefore for locus *l*:

$$\Pr(data \mid I_l = 1) = \Pr(readcount_l \mid I_l = 1)\Pr(length - distribution_l \mid I_l = 1) \qquad (3)$$

And *Prob(data|I_l=0)* for null locus *l:*

$$\Pr(data \mid I_l = 0) = \Pr(readcount_l \mid I_l = 0)\Pr(length - distribution_l \mid I_l = 0) \qquad (4)$$

**Negative binomial model for read count data**

Negative binomial distribution is applied to model read count data. $u_s$ and $\varphi_s$ denotes the mean count and dispersion for sRNA loci, respectively; $u_n$ and $\varphi_n$ denotes the mean count and dispersion for null loci, respectively. Therefore, for specific locus *l*, we have:

$$y_l \mid (u_s, \varphi_s, I_l = 1) \sim NB(u_s, \varphi_s)$$

$$y_l \mid (u_n, \varphi_n, I_l = 0) \sim NB(u_n, \varphi_n) \qquad (5)$$

where $y_j$ is a read count for a given locus *j*.

Finally, based on the negative binomial probability mass function, we have:

$$\Pr(readcount_l \mid I_l = 0) = P(Y = y_j \mid \mu_n, \varphi_n) = \frac{\Gamma(y + \varphi_n^{-1})}{\Gamma(\varphi_n^{-1})\Gamma(y+1)}\left(\frac{1}{1+\mu_n\varphi_n}\right)^{\varphi^{-1}}\left(\frac{\mu_n}{\varphi_n^{-1}+\mu_n}\right)^{y}$$

**Table S1.** Primers used in this study.

| Primer Name | Sequence | Purpose |
|---|---|---|
| TamRNA1-1 F | GCGGCTACAAGTTGATGATGAC | qRT-PCR (wheat target mRNA Traes_2AS_EC975D5AB) |
| TamRNA1-1 R | TCTCCGGACGACCACAATATTG | qRT-PCR (wheat target mRNA Traes_2AS_EC975D5AB) |
| TamRNA1-2 Fnew | TTCTGTGTGCCTGGATTGTATGTC | qRT-PCR (wheat target mRNA Traes_2BS_E5732CD2C) |
| TamRNA1-2 Rnew | GAACTTGTATGATGTGTTGAAAGTGGCG | qRT-PCR (wheat target mRNA Traes_2BS_E5732CD2C) |
| TamRNA1-3 Fnew | TGGAGCCCAGCAAGCACTCGG | qRT-PCR (wheat target mRNA Traes_2DS_9B86CE58D) |
| TamRNA1-3 Rnew | GCTTGTAATGCAGGGCACAACG | qRT-PCR (wheat target mRNA Traes_2DS_9B86CE58D) |
| TamRNA 2Fnew | CCGTAACATTCACTGGGTATAAATC | qRT-PCR (wheat target mRNA Traes_4BS_5E12F0B27) |
| TamRNA 2 Rnew | AAATATACTCTAGATGCACCAGGC | qRT-PCR (wheat target mRNA Traes_4BS_5E12F0B27) |
| TamRNA3-1 F | CCTTCCTCTCGGTGGGTGTTG | qRT-PCR (wheat target mRNA Traes_4AS_92067319F) |
| TamRNA 3-1 R | CCCATCTTTGCAGCGCCTGCT | qRT-PCR (wheat target mRNA Traes_4AS_92067319F) |
| TamRNA 3-2 Fnew | TGAACAGAACTGTATCTGCTGTAAG | qRT-PCR (wheat target mRNA Traes_4BL_2136196AE) |
| TamRNA 3-2 R | ATGCGGCGGCGGGTCATGG | qRT-PCR (wheat target mRNA Traes_4BL_2136196AE) |
| TamRNA4-1 Fnew | TCAGTCGCGTACCTTCTTGTC | qRT-PCR (wheat target mRNA Traes_5BS_409B24307) |
| TamRNA4-1 Rnew | CGAAGTGTTCTGACAAAGCCG | qRT-PCR (wheat target mRNA Traes_5BS_409B24307) |
| TamRNA4-2 F | CATTTAGGCGCCAGAGAATAATGC | qRT-PCR (wheat target mRNA Traes_5BS_2C2072466) |
| TamRNA4-2R | GACAGGAACAGGTACTCCCATG | qRT-PCR (wheat target mRNA Traes_5BS_2C2072466) |
| TaCDC48 newF | GTCCTCCTGGCTGTGGTAAAAC | wheat reference gene for plant tissue qPCR |
| TaCDC48 newR | AGCAGCTCAGGTCCCTTGATAC | wheat reference gene for plant tissue qPCR |
| ZtB-tubulin F | CCTACTTCGTCGAGTGGATT | *Z. tritici* reference gene for fungal tissue stem-loop qPCR |
| ZtB-tubulin R | TACTCGGACACAAGATCGTT | *Z. tritici* reference gene for fungal tissue stem-loop qPCR |
| universal R | GTGCAGGGTCCGAGGT | stem-loop qRT-PCR (universal reverse primer for all sRNA targets) |
| Zt-sRNA1 F | GCGATCTTGGGGAATCCGTAG | stem-loop qRT-PCR |
| Zt-sRNA1 SLRT | GTCGTATCCAGTGCAGGGTCCGAGGTATTCGCACTGGATACGACACAACA | stem-loop cDNA synthesis |
| Zt-sRNA2 F | TGGACTGCACTGGTTGCTCG | stem-loop qRT-PCR |
| Zt-sRNA2 SLRT | GTCGTATCCAGTGCAGGGTCCGAGGTATTCGCACTGGATACGACAGCGTT | stem-loop cDNA synthesis |
| Zt-sRNA3 F | GGCAGTAACCATCTTTCGGGT | stem-loop qRT-PCR |
| Zt-sRNA3 SLRT | GTCGTATCCAGTGCAGGGTCCGAGGTATTCGCACTGGATACGACAGTCAG | stem-loop cDNA synthesis |
| Zt-sRNA4 F | GTGAACTGGAGATCGCGAAGG | stem-loop qRT-PCR |
| Zt-sRNA4 SLRT | GTCGTATCCAGTGCAGGGTCCGAGGTATTCGCACTGGATACGACGAACCT | stem-loop cDNA synthesis |
| ZtDCL-1 | GGGCCCGGCGCGCCGAATTCGAGCTGACAACTATCTGATCTATTGCG | Gibson cloning of the KO construct into vector pCHYG |
| ZtDCL-2 | TCAGTTATCGAATATTGTGTTTCGACGACATC | Gibson cloning of the KO construct into vector pCHYG |
| ZtDCL-3 | GTCGAAACACAATATTCGATAACTGATATTGAAGG | Gibson cloning of the KO construct into vector pCHYG |
| ZtDCL-4 | CATCCAACCACCCCTCGAGGTCGACGGTATC | Gibson cloning of the KO construct into vector pCHYG |
| ZtDCL-5 | CGTCGACCTCGAGGGGTGGTTGGATGGTGATTG | Gibson cloning of the KO construct into vector pCHYG |
| ZtDCL-6 | CACGTGGTGGTGGTGGTGGTGGCTAGCGTTAACACGGCCATTCCGTTCACGA | Gibson cloning of the KO construct into vector pCHYG |
| ZtAGO2-1 | GGGCCCGGCGCGCCGAATTCGAGCTGCCTTGGACTCGGACCGG | Gibson cloning of the KO construct into vector pCHYG |
| ZtAGO2-2 | TCAGTTATCGAATGACAGTGCGGTTGGTGGG | Gibson cloning of the KO construct into vector pCHYG |
| ZtAGO2-3 | CAACCGCACTGTCATTCGATAACTGATATTGAAGG | Gibson cloning of the KO construct into vector pCHYG |
| ZtAGO2-4 | GCTGAAGTGACTTCTCGAGGTCGACGGTATC | Gibson cloning of the KO construct into vector pCHYG |
| ZtAGO2-5 | CGTCGACCTCGAGAAGTCACTTCAGCTTCTTTTC | Gibson cloning of the KO construct into vector pCHYG |
| ZtAGO2-6 | CACGTGGTGGTGGTGGTGGTGGCTAGCGTTAACAGGAGAGAGGGATGATACC | Gibson cloning of the KO construct into vector pCHYG |
| ZtAGO1-1 | GGGCCCGGCGCGCCGAATTCGAGCTCTCGAGCCTCCACCACTAC | Gibson cloning of the KO construct into vector pCHYG |
| ZtAGO1-2 | TCAGTTATCGAATGAAGTTGAGGTTGGGTAACG | Gibson cloning of the KO construct into vector pCHYG |
| ZtAGO1-3 | CAACCTCAACTTCATTCGATAACTGATATTGAAGG | Gibson cloning of the KO construct into vector pCHYG |
| ZtAGO1-4 | TCTTCGAGCTCTCCTCGAGGTCGACGGTATC | Gibson cloning of the KO construct into vector pCHYG |
| ZtAGO1-5 | CGTCGACCTCGAGGAGAGCTCGAAGAGAAATC | Gibson cloning of the KO construct into vector pCHYG |
| ZtAGO1-6 | CACGTGGTGGTGGTGGTGGTGGCTAGCGTTAACACTTACCTGTACGCCTATTCAG | Gibson cloning of the KO construct into vector pCHYG |
| ztTUBalpha_F2 | aaggaagtttaaCACCTACCGCAGCCTTTTCCAC | BSMV-HIGS |
| ztTUBalpha_R2 | aaccaccaccaccgtGTGCTGGGTACACGCAGAACTCA | BSMV-HIGS |
| ztTUBbeta_F1 | aaggaagtttaaCACCCAGCAAATCTTCGACCCTA | BSMV-HIGS |
| ztTUBbeta_R1 | aaccaccaccaccgtCTGAACATGGCGGAGAACTGGTC | BSMV-HIGS |
| ztCYP51_F1 | aaggaagtttaaCCGTTTTGGACTCTCGTCTCTT | BSMV-HIGS |
| ztCYP51_R1 | aaccaccaccaccgtCAGTCCTACGTGACCTTGATCG | BSMV-HIGS |
| ztALG2_F1 | aaggaagtttaaCCTCGTGTACTTGCTGTTCACC | BSMV-HIGS |
| ztALG2_R1 | aaccaccaccaccgtCGATTCGCGATCCTCTGTACTA | BSMV-HIGS |
| ztTUBalpha_T7P_F2 | tcctaatacgactcactatagggagCACCTACCGCAGCCTTTTCCAC | in vitro RNAi |
| ztTUBalpha_T7P_R2 | tcctaatacgactcactatagggagGTGCTGGGTACACGCAGAACTCA | in vitro RNAi |
| ztTUBbeta_T7P_F1 | tcctaatacgactcactatagggagCACCCAGCAAATCTTCGACCCTA | in vitro RNAi |
| ztTUBbeta_T7P_R1 | tcctaatacgactcactatagggagCTGAACATGGCGGAGAACTGGTC | in vitro RNAi |
| ztCYP51_T7P_F1 | tcctaatacgactcactatagggagCCGTTTTGGACTCTCGTCTCTT | in vitro RNAi |
| ztCYP51_T7P_R1 | tcctaatacgactcactatagggagCAGTCCTACGTGACCTTGATCG | in vitro RNAi |
| GFP_T7P_F1 | tcctaatacgactcactatagggagGCACAAATTTTCTGTCAGTGGA | in vitro RNAi |
| GFP_T7P_R1 | tcctaatacgactcactatagggagGTCCGAGAATGTTTCCATCTTC | in vitro RNAi |

**Table S2.** Expression levels of the key predicted RNAi machinery components in *Zymoseptoria tritici* isolate IPO323.

| | | | Mean FPKM values† | | | | | | |
|---|---|---|---|---|---|---|---|---|---|
| **Gene** | **Ensembl Fungi gene code** | **Locus** | **CDB‡** | **PDB** | **1 dpi** | **4 dpi** | **9 dpi** | **14 dpi** | **21 dpi** |
| *ZtDCL1* | Mycgr3G47983 | 9:1660746-1665410 | 1.3 | 1.7 | 1.0 | 0.7 | 1.1 | 1.3 | 2.2 |
| *ZtAGO1* | Mycgr3G38035 | 3:2630539-2633932 | 8.9 | 3.0 | 14.1 | 5.4 | 18.8 | 11.7 | 18.1 |
| *ZtAGO2* | Mycgr3G10621 | 11:940972-943399 | 4.0 | 2.7 | 4.8 | 5.1 | 3.7 | 2.6 | 3.4 |
| *pseudoAGO3* | Mycgr3G90232 | 1:5667618-5671029 | 0.1 | 0.2 | 0.3 | 0.8 | 0.4 | 0.1 | 0.2 |
| *pseudoAGO4* | Mycgr3G25632 | 1:4995448-4996504 | 1.3 | 1.3 | 7.2 | 0.0 | 2.0 | 2.0 | 1.0 |
| *ZtRDRP1* | Mycgr3G51407 | 13:605705-608630 | 1.0 | 0.9 | 1.5 | 1.6 | 1.3 | 0.7 | 0.4 |
| *ZtRDRP2* | Mycgr3G49833 | 11:202398-206395 | 0.6 | 0.5 | 0.2 | 0.6 | 1.4 | 1.2 | 1.4 |

† Expression levels of the candidate genes was inferred based on the previously published RNAseq data (Rudd *et al.,* 2015) and is shown as FPKM (fragments per kilobase of exon model per million reads mapped) values

‡ *Zymoseptoria tritici* isolate IPO323 was grown in vitro in low nutrients CDB (Czapek Dox Broth) or high nutrients PDB (Potato Dextrose Broth) media. The same fungal isolate was also inoculated onto the leaves of susceptible wheat cv. Riband and gene expression assessed at 1, 4, 9, 14, and 21 dpi (days post inoculation).

**Table S3.** Computationally predicted *Zymoseptoria tritici* IPO323 sRNA loci.

| sRNA locus ID | length (nt) | read counts from all samples | origin | sRNA loci active at 4, 9, 13, or 21 dpi during wheat infection | | | | posterior probability difference† | Min FDR differential expression‡ |
|---|---|---|---|---|---|---|---|---|---|
| 1.1088031.1088594 | 564 | 938 | intergenic | BWZt4 | BWZt9 | | BWZt21 | 1.00 | 0.00 |
| 1.132780.135800 | 3021 | 498 | gene | | | BWZt13 | BWZt21 | 1.00 | 0.01 |
| 1.1432722.1433785 | 1064 | 6332 | gene | BWZt4 | BWZt9 | BWZt13 | BWZt21 | 1.00 | 0.00 |
| 1.168753.169514 | 762 | 7814 | intergenic | BWZt4 | BWZt9 | BWZt13 | BWZt21 | 1.00 | 0.00 |
| 1.2.24760 | 24759 | 29600 | intergenic | BWZt4 | BWZt9 | BWZt13 | BWZt21 | 0.99 | 0.00 |
| 1.2088218.2090311 | 2094 | 525 | gene | | | | BWZt21 | 1.00 | 0.02 |
| 1.24832.33907 | 9076 | 7406 | intergenic | BWZt4 | BWZt9 | BWZt13 | BWZt21 | 1.00 | 0.00 |
| 1.2630102.2631752 | 1651 | 1029 | gene | | | | BWZt21 | 1.00 | 0.00 |
| 1.3024339.3025761 | 1423 | 1546 | gene | | | BWZt13 | BWZt21 | 1.00 | 0.02 |
| 1.3107284.3109571 | 2288 | 17700 | gene | BWZt4 | BWZt9 | BWZt13 | BWZt21 | 1.00 | 0.00 |
| 1.3175256.3175844 | 589 | 2550 | gene | BWZt4 | BWZt9 | BWZt13 | BWZt21 | 1.00 | 0.00 |
| 1.3404853.3405015 | 163 | 648000 | intergenic | BWZt4 | BWZt9 | BWZt13 | BWZt21 | 1.00 | 0.00 |
| 1.3788931.3792998 | 4068 | 470 | gene | | | | BWZt21 | 1.00 | 0.02 |
| 1.3817478.3818480 | 1003 | 282 | gene | | | | BWZt21 | 1.00 | 0.02 |
| 1.3864242.3867547 | 3306 | 423 | gene | | | | BWZt21 | 1.00 | 0.01 |
| 1.4059657.4059846 | 190 | 840 | intergenic | | | BWZt13 | BWZt21 | 1.00 | 0.01 |
| 1.4098242.4102440 | 4199 | 762 | gene | | | BWZt13 | BWZt21 | 1.00 | 0.00 |
| 1.4411833.4412084 | 252 | 31300 | pseudogenic_tRNA | BWZt4 | BWZt9 | BWZt13 | BWZt21 | 1.00 | 0.00 |
| 1.4420671.4420786 | 116 | 2793 | gene | | | BWZt13 | | 1.00 | 0.00 |
| 1.4461121.4461429 | 309 | 2637 | tRNA_gene | BWZt4 | BWZt9 | BWZt13 | BWZt21 | 1.00 | 0.00 |
| 1.4500965.4502701 | 1737 | 872 | gene | | | BWZt13 | | 1.00 | 0.02 |
| 1.4519579.4520495 | 917 | 6544 | intergenic | BWZt4 | BWZt9 | BWZt13 | BWZt21 | 1.00 | 0.00 |
| 1.4532087.4532257 | 171 | 2776 | rRNA_gene | BWZt4 | BWZt9 | BWZt13 | BWZt21 | 1.00 | 0.00 |
| 1.4859784.4862055 | 2272 | 1326 | gene | | | | BWZt21 | 1.00 | 0.01 |
| 1.5027663.5030714 | 3052 | 1058 | gene | | | BWZt13 | BWZt21 | 1.00 | 0.00 |
| 1.5208279.5209283 | 1005 | 6982 | intergenic | BWZt4 | BWZt9 | BWZt13 | BWZt21 | 1.00 | 0.00 |
| 1.52723.55652 | 2930 | 2665 | intergenic | | BWZt9 | BWZt13 | BWZt21 | 1.00 | 0.00 |
| 1.5370559.5371316 | 758 | 1768 | gene | | | | BWZt21 | 1.00 | 0.01 |
| 1.55724.59340 | 3617 | 3434 | intergenic | | BWZt9 | BWZt13 | BWZt21 | 1.00 | 0.00 |
| 1.5574650.5575522 | 873 | 9652 | intergenic | | | BWZt13 | BWZt21 | 1.00 | 0.00 |
| 1.5935918.5953168 | 17251 | 12900 | gene | | | BWZt13 | | 1.00 | 0.00 |
| 1.59399.69532 | 10134 | 7238 | intergenic | | BWZt9 | BWZt13 | BWZt21 | 1.00 | 0.00 |
| 1.6022523.6024517 | 1995 | 2563 | intergenic | | BWZt9 | BWZt13 | BWZt21 | 1.00 | 0.00 |
| 1.6024577.6027831 | 3255 | 1941 | intergenic | | BWZt9 | BWZt13 | BWZt21 | 1.00 | 0.00 |
| 1.6042189.6044283 | 2095 | 2270 | intergenic | | BWZt9 | BWZt13 | BWZt21 | 1.00 | 0.00 |
| 1.6044579.6052106 | 7528 | 5140 | intergenic | | BWZt9 | BWZt13 | BWZt21 | 1.00 | 0.00 |
| 1.6052501.6055814 | 3314 | 1568 | intergenic | | BWZt9 | BWZt13 | BWZt21 | 1.00 | 0.00 |
| 1.6055877.6056728 | 852 | 487 | intergenic | | BWZt9 | BWZt13 | BWZt21 | 1.00 | 0.01 |
| 1.6060199.6062484 | 2286 | 970 | intergenic | | BWZt9 | BWZt13 | BWZt21 | 1.00 | 0.00 |
| 1.6062562.6077812 | 15251 | 13200 | intergenic | | BWZt9 | BWZt13 | BWZt21 | 1.00 | 0.00 |
| 1.6077865.6084551 | 6687 | 4298 | intergenic | | BWZt9 | BWZt13 | BWZt21 | 1.00 | 0.00 |
| 1.610884.611019 | 136 | 2523 | intergenic | | BWZt9 | BWZt13 | BWZt21 | 1.00 | 0.00 |
| 1.681033.681226 | 194 | 1345 | gene | | BWZt9 | BWZt13 | BWZt21 | 1.00 | 0.00 |
| 1.686410.686631 | 222 | 872 | gene | | | | BWZt21 | 1.00 | 0.04 |
| 1.802150.802635 | 486 | 4664 | intergenic | | | BWZt13 | | 1.00 | 0.00 |
| 1.809089.809248 | 160 | 1207 | intergenic | | | BWZt13 | | 1.00 | 0.01 |
| 1.912518.913393 | 876 | 5584 | intergenic | | BWZt9 | BWZt13 | BWZt21 | 1.00 | 0.00 |
| 1.952268.954112 | 1845 | 2446 | gene | | | BWZt13 | BWZt21 | 1.00 | 0.00 |
| 2.1061972.1062929 | 958 | 9003 | intergenic | BWZt4 | BWZt9 | BWZt13 | BWZt21 | 1.00 | 0.00 |
| 2.1128831.1132146 | 3316 | 1909 | gene | | BWZt9 | BWZt13 | BWZt21 | 1.00 | 0.00 |
| 2.1295279.1297239 | 1961 | 338 | gene | | | | BWZt21 | 1.00 | 0.01 |
| 2.1326532.1328523 | 1992 | 721 | gene | | | BWZt13 | | 1.00 | 0.03 |
| 2.1333569.1334006 | 438 | 2840 | intergenic | | | BWZt13 | BWZt21 | 1.00 | 0.00 |
| 2.1441675.1441950 | 276 | 70400 | tRNA_gene | BWZt4 | BWZt9 | BWZt13 | BWZt21 | 1.00 | 0.00 |
| 2.1664358.1664628 | 271 | 513 | intergenic | | | BWZt13 | BWZt21 | 1.00 | 0.00 |
| 2.1741314.1742939 | 1626 | 10100 | gene | BWZt4 | BWZt9 | BWZt13 | BWZt21 | 1.00 | 0.00 |
| 2.17721.18726 | 1006 | 6063 | intergenic | | BWZt9 | BWZt13 | BWZt21 | 1.00 | 0.00 |
| 2.1895220.1896101 | 882 | 3764 | intergenic | BWZt4 | BWZt9 | BWZt13 | BWZt21 | 1.00 | 0.00 |
| 2.1903263.1903455 | 193 | 648000 | intergenic | BWZt4 | BWZt9 | BWZt13 | BWZt21 | 1.00 | 0.00 |
| 2.1909278.1910404 | 1127 | 6408 | intergenic | BWZt4 | BWZt9 | BWZt13 | BWZt21 | 1.00 | 0.00 |
| 2.1925528.1927912 | 2385 | 279 | gene | | | | BWZt21 | 1.00 | 0.03 |
| 2.2093476.2093705 | 230 | 16200 | gene | BWZt4 | | BWZt13 | BWZt21 | 1.00 | 0.00 |
| 2.2362398.2364568 | 2171 | 454 | gene | | | | BWZt21 | 1.00 | 0.00 |
| 2.236665.237674 | 1010 | 8376 | gene | BWZt4 | BWZt9 | BWZt13 | BWZt21 | 1.00 | 0.00 |
| 2.2743117.2744701 | 1585 | 281 | gene | | | | BWZt21 | 1.00 | 0.02 |
| 2.2748785.2750628 | 1844 | 1124 | gene | | | | BWZt21 | 1.00 | 0.01 |
| 2.2777908.2780777 | 2870 | 1254 | gene | | | | BWZt21 | 0.99 | 0.02 |
| 2.2865736.2866983 | 1248 | 1445 | gene | | BWZt9 | | | 1.00 | 0.03 |
| 2.3153789.3154883 | 1095 | 1498 | gene | | | BWZt13 | | 1.00 | 0.00 |
| 2.3187737.3188483 | 747 | 6216 | intergenic | BWZt4 | BWZt9 | BWZt13 | BWZt21 | 1.00 | 0.00 |
| 2.3189040.3189775 | 736 | 7256 | intergenic | BWZt4 | BWZt9 | BWZt13 | BWZt21 | 1.00 | 0.00 |
| 2.3276658.3277710 | 1053 | 8657 | intergenic | BWZt4 | BWZt9 | BWZt13 | BWZt21 | 1.00 | 0.00 |

| | | | | | | | | | |
|---|---|---|---|---|---|---|---|---|---|
| 2.3411312.3412084 | 773 | 2302 | intergenic | | BWZt9 | BWZt13 | BWZt21 | 1.00 | 0.00 |
| 2.341988.345005 | 3018 | 17800 | gene | BWZt4 | BWZt9 | BWZt13 | BWZt21 | 1.00 | 0.00 |
| 2.3632877.3633814 | 938 | 4419 | intergenic | BWZt4 | BWZt9 | BWZt13 | BWZt21 | 1.00 | 0.00 |
| 2.3801864.3816890 | 15027 | 15300 | intergenic | BWZt4 | BWZt9 | BWZt13 | BWZt21 | 1.00 | 0.00 |
| 2.3816969.3821391 | 4423 | 3528 | intergenic | | BWZt9 | BWZt13 | BWZt21 | 1.00 | 0.00 |
| 2.3821445.3822997 | 1553 | 596 | intergenic | | BWZt9 | BWZt13 | BWZt21 | 1.00 | 0.00 |
| 2.3823060.3839541 | 16482 | 16200 | tRNA_gene | | BWZt9 | BWZt13 | BWZt21 | 1.00 | 0.00 |
| 2.3839620.3844042 | 4423 | 3603 | intergenic | | BWZt9 | BWZt13 | BWZt21 | 1.00 | 0.00 |
| 2.3844121.3848381 | 4261 | 1931 | tRNA_gene | | BWZt9 | BWZt13 | BWZt21 | 1.00 | 0.00 |
| 2.400134.400399 | 266 | 6122 | intergenic | BWZt4 | BWZt9 | BWZt13 | BWZt21 | 1.00 | 0.00 |
| 2.47729.49709 | 1981 | 7360 | intergenic | BWZt4 | BWZt9 | BWZt13 | BWZt21 | 1.00 | 0.00 |
| 2.532319.532572 | 254 | 648000 | intergenic | BWZt4 | BWZt9 | BWZt13 | BWZt21 | 1.00 | 0.00 |
| 2.715507.715678 | 172 | 2319 | intergenic | | | BWZt13 | | 1.00 | 0.01 |
| 2.819512.820307 | 796 | 5263 | intergenic | BWZt4 | BWZt9 | BWZt13 | BWZt21 | 1.00 | 0.00 |
| 3.1178.7084 | 5907 | 3149 | intergenic | | BWZt9 | BWZt13 | BWZt21 | 1.00 | 0.00 |
| 3.1848836.1849733 | 898 | 6302 | intergenic | BWZt4 | BWZt9 | BWZt13 | BWZt21 | 1.00 | 0.00 |
| 3.1987288.1987894 | 607 | 2124 | intergenic | | | | BWZt21 | 1.00 | 0.01 |
| 3.2136686.2136750 | 65 | 331 | snRNA_gene | BWZt4 | | | | 1.00 | 0.01 |
| 3.218432.218715 | 284 | 1229 | pseudogenic_tRNA | | | | BWZt21 | 1.00 | 0.00 |
| 3.2252943.2253304 | 362 | 3819 | intergenic | | | BWZt13 | BWZt21 | 1.00 | 0.00 |
| 3.2292170.2292823 | 654 | 248 | intergenic | BWZt4 | | | | 1.00 | 0.05 |
| 3.2312849.2313471 | 623 | 1603 | tRNA_gene | | | | BWZt21 | 1.00 | 0.05 |
| 3.2375162.2376724 | 1563 | 3326 | gene | | | BWZt13 | | 1.00 | 0.00 |
| 3.2469896.2470926 | 1031 | 550 | intergenic | | | | BWZt21 | 1.00 | 0.04 |
| 3.2711296.2712607 | 1312 | 508 | gene | | | | BWZt21 | 1.00 | 0.00 |
| 3.2736368.2736981 | 614 | 5916 | intergenic | BWZt4 | | BWZt13 | BWZt21 | 1.00 | 0.00 |
| 3.2779587.2780080 | 494 | 594 | intergenic | | | BWZt13 | | 1.00 | 0.02 |
| 3.2791693.2793471 | 1779 | 523 | gene | BWZt4 | | | BWZt21 | 1.00 | 0.03 |
| 3.3048306.3049442 | 1137 | 1168 | gene | | | BWZt13 | BWZt21 | 1.00 | 0.00 |
| 3.3241153.3242334 | 1182 | 651000 | gene | BWZt4 | BWZt9 | BWZt13 | BWZt21 | 1.00 | 0.00 |
| 3.3403412.3404238 | 827 | 8607 | intergenic | BWZt4 | BWZt9 | BWZt13 | BWZt21 | 1.00 | 0.00 |
| 3.3468771.3476338 | 7568 | 2969 | intergenic | | BWZt9 | BWZt13 | BWZt21 | 1.00 | 0.00 |
| 3.3486626.3493394 | 6769 | 1790 | intergenic | | BWZt9 | BWZt13 | BWZt21 | 1.00 | 0.00 |
| 3.3496452.3502010 | 5559 | 1689 | gene | | BWZt9 | BWZt13 | BWZt21 | 1.00 | 0.00 |
| 3.387886.388363 | 478 | 867 | intergenic | | | BWZt13 | BWZt21 | 1.00 | 0.01 |
| 3.391037.392727 | 1691 | 5811 | intergenic | BWZt4 | BWZt9 | BWZt13 | BWZt21 | 1.00 | 0.00 |
| 3.401660.407274 | 5615 | 5891 | intergenic | | BWZt9 | BWZt13 | BWZt21 | 1.00 | 0.00 |
| 3.448737.450457 | 1721 | 379 | gene | | | | BWZt21 | 1.00 | 0.01 |
| 3.451342.453408 | 2067 | 395 | gene | | | | BWZt21 | 1.00 | 0.00 |
| 3.587789.590578 | 2790 | 8793 | intergenic | BWZt4 | BWZt9 | BWZt13 | BWZt21 | 1.00 | 0.00 |
| 3.699143.699357 | 215 | 4222 | pseudogenic_tRNA | | | BWZt13 | BWZt21 | 1.00 | 0.00 |
| 3.7151.18710 | 11560 | 12200 | intergenic | BWZt4 | BWZt9 | BWZt13 | BWZt21 | 1.00 | 0.00 |
| 3.759692.759839 | 148 | 1314 | intergenic | | BWZt9 | BWZt13 | BWZt21 | 1.00 | 0.00 |
| 3.822679.823459 | 781 | 8275 | intergenic | BWZt4 | BWZt9 | BWZt13 | BWZt21 | 1.00 | 0.00 |
| 3.957606.958577 | 972 | 6221 | intergenic | BWZt4 | BWZt9 | BWZt13 | BWZt21 | 1.00 | 0.00 |
| 4.10488.12213 | 1726 | 565 | intergenic | | | BWZt13 | BWZt21 | 1.00 | 0.01 |
| 4.1251113.1252024 | 912 | 7241 | intergenic | BWZt4 | BWZt9 | BWZt13 | BWZt21 | 1.00 | 0.00 |
| 4.1428859.1429718 | 860 | 6218 | intergenic | BWZt4 | BWZt9 | BWZt13 | BWZt21 | 1.00 | 0.00 |
| 4.170472.171758 | 1287 | 7978 | intergenic | BWZt4 | BWZt9 | BWZt13 | BWZt21 | 1.00 | 0.00 |
| 4.2028875.2030805 | 1931 | 3750 | gene | BWZt4 | BWZt9 | | BWZt21 | 1.00 | 0.00 |
| 4.2215448.2215980 | 533 | 1231 | intergenic | BWZt4 | BWZt9 | | BWZt21 | 1.00 | 0.00 |
| 4.2228316.2228727 | 412 | 5160 | intergenic | | | BWZt13 | BWZt21 | 1.00 | 0.00 |
| 4.2261891.2262208 | 318 | 780 | intergenic | | | BWZt13 | | 1.00 | 0.02 |
| 4.234947.235725 | 779 | 6692 | intergenic | BWZt4 | BWZt9 | BWZt13 | BWZt21 | 1.00 | 0.00 |
| 4.241532.242398 | 867 | 7016 | intergenic | | BWZt9 | BWZt13 | BWZt21 | 1.00 | 0.00 |
| 4.2562105.2562351 | 247 | 14200 | pseudogenic_tRNA | BWZt4 | BWZt9 | BWZt13 | BWZt21 | 1.00 | 0.00 |
| 4.2644748.2645586 | 839 | 7707 | intergenic | BWZt4 | BWZt9 | BWZt13 | BWZt21 | 1.00 | 0.00 |
| 4.272845.274005 | 1161 | 6626 | intergenic | BWZt4 | BWZt9 | BWZt13 | BWZt21 | 1.00 | 0.00 |
| 4.2866934.2868658 | 1725 | 5776 | intergenic | BWZt4 | BWZt9 | BWZt13 | BWZt21 | 1.00 | 0.00 |
| 4.2869853.2880010 | 10158 | 6478 | intergenic | | BWZt9 | BWZt13 | | 1.00 | 0.00 |
| 4.356163.358445 | 2283 | 577 | gene | | | | BWZt21 | 1.00 | 0.02 |
| 4.595344.597737 | 2394 | 796 | gene | | | | BWZt21 | 1.00 | 0.02 |
| 4.8404.10428 | 2025 | 1273 | intergenic | | BWZt9 | BWZt13 | BWZt21 | 1.00 | 0.00 |
| 4.919278.919425 | 148 | 4403 | pseudogenic_tRNA | BWZt4 | BWZt9 | BWZt13 | BWZt21 | 1.00 | 0.00 |
| 5.1045197.1045805 | 609 | 3609 | snRNA_gene | BWZt4 | BWZt9 | BWZt13 | | 1.00 | 0.00 |
| 5.1092026.1092263 | 238 | 593 | intergenic | | BWZt9 | BWZt13 | | 1.00 | 0.00 |
| 5.1092560.1092727 | 168 | 7446 | snoRNA_gene | BWZt4 | | | | 1.00 | 0.00 |
| 5.1098642.1098834 | 193 | 7171 | intergenic | BWZt4 | BWZt9 | BWZt13 | BWZt21 | 1.00 | 0.00 |
| 5.1130578.1135063 | 4486 | 1317 | gene | | | BWZt13 | BWZt21 | 1.00 | 0.00 |
| 5.181519.182321 | 803 | 7098 | intergenic | BWZt4 | BWZt9 | BWZt13 | BWZt21 | 1.00 | 0.00 |
| 5.1875780.1875912 | 133 | 2761 | intergenic | BWZt4 | BWZt9 | BWZt13 | BWZt21 | 1.00 | 0.00 |
| 5.2094726.2095999 | 1274 | 566 | gene | | | BWZt13 | BWZt21 | 1.00 | 0.00 |
| 5.2318983.2319321 | 339 | 11400 | intergenic | BWZt4 | BWZt9 | BWZt13 | BWZt21 | 1.00 | 0.00 |
| 5.2486971.2487211 | 241 | 3039 | tRNA_gene | | | | BWZt21 | 1.00 | 0.04 |
| 5.250300.250376 | 77 | 680 | gene | | | BWZt13 | | 1.00 | 0.00 |
| 5.2645459.2646021 | 563 | 836 | intergenic | | | | BWZt21 | 1.00 | 0.02 |
| 5.2686132.2687400 | 1269 | 9195 | intergenic | BWZt4 | BWZt9 | BWZt13 | BWZt21 | 1.00 | 0.00 |
| 5.2792872.2793631 | 760 | 138 | gene | BWZt4 | | | | 1.00 | 0.03 |

| | | | | | | | | | |
|---|---|---|---|---|---|---|---|---|---|
| 5.2806545.2846948 | 40404 | 33100 | intergenic | BWZt4 | BWZt9 | BWZt13 | BWZt21 | 1.00 | 0.00 |
| 5.2850297.2861802 | 11506 | 12400 | intergenic | | BWZt9 | BWZt13 | BWZt21 | 1.00 | 0.00 |
| 5.308748.309026 | 279 | 358 | intergenic | | | | BWZt21 | 1.00 | 0.03 |
| 5.514262.514929 | 668 | 648000 | gene | BWZt4 | BWZt9 | BWZt13 | BWZt21 | 1.00 | 0.00 |
| 5.523622.524805 | 1184 | 5827 | intergenic | BWZt4 | BWZt9 | BWZt13 | BWZt21 | 1.00 | 0.00 |
| 5.557661.560726 | 3066 | 17900 | gene | BWZt4 | BWZt9 | BWZt13 | BWZt21 | 1.00 | 0.00 |
| 5.561012.563057 | 2046 | 534 | gene | | | | BWZt21 | 1.00 | 0.01 |
| 5.59060.61544 | 2485 | 1647 | intergenic | | BWZt9 | BWZt13 | BWZt21 | 1.00 | 0.00 |
| 5.748438.748459 | 22 | 4191 | gene | | BWZt9 | BWZt13 | BWZt21 | 1.00 | 0.00 |
| 5.849612.850341 | 730 | 367 | gene | | | | BWZt21 | 1.00 | 0.01 |
| 5.912353.914719 | 2367 | 399 | gene | BWZt4 | | | | 1.00 | 0.02 |
| 5.918436.919203 | 768 | 7368 | intergenic | BWZt4 | BWZt9 | BWZt13 | BWZt21 | 1.00 | 0.00 |
| 5.934505.936848 | 2344 | 6860 | intergenic | BWZt4 | BWZt9 | BWZt13 | BWZt21 | 1.00 | 0.00 |
| 6.1267883.1270184 | 2302 | 1486 | intergenic | | BWZt9 | BWZt13 | BWZt21 | 1.00 | 0.00 |
| 6.13518.16550 | 3033 | 1539 | intergenic | | BWZt9 | BWZt13 | BWZt21 | 1.00 | 0.00 |
| 6.16617.39543 | 22927 | 21100 | intergenic | BWZt4 | BWZt9 | BWZt13 | BWZt21 | 1.00 | 0.00 |
| 6.173879.174893 | 1015 | 7114 | intergenic | | BWZt9 | BWZt13 | BWZt21 | 1.00 | 0.00 |
| 6.1808235.1808688 | 454 | 11300 | tRNA_gene | BWZt4 | BWZt9 | BWZt13 | BWZt21 | 1.00 | 0.00 |
| 6.2022916.2027069 | 4154 | 9928 | intergenic | BWZt4 | BWZt9 | BWZt13 | BWZt21 | 1.00 | 0.00 |
| 6.2108904.2109094 | 191 | 3494 | snRNA_gene | | | BWZt13 | BWZt21 | 1.00 | 0.00 |
| 6.212702.213658 | 957 | 4395 | intergenic | BWZt4 | BWZt9 | BWZt13 | BWZt21 | 1.00 | 0.00 |
| 6.218141.218388 | 248 | 1012 | intergenic | | | BWZt13 | BWZt21 | 1.00 | 0.00 |
| 6.2197358.2198045 | 688 | 1025 | intergenic | | BWZt9 | BWZt13 | BWZt21 | 1.00 | 0.00 |
| 6.231370.231410 | 41 | 303 | miRNA | BWZt4 | BWZt9 | | | 1.00 | 0.01 |
| 6.2414949.2415730 | 782 | 6236 | intergenic | BWZt4 | BWZt9 | BWZt13 | BWZt21 | 1.00 | 0.00 |
| 6.39611.43211 | 3601 | 4096 | intergenic | | BWZt9 | BWZt13 | BWZt21 | 1.00 | 0.00 |
| 6.597638.598424 | 787 | 6271 | intergenic | BWZt4 | BWZt9 | BWZt13 | BWZt21 | 1.00 | 0.00 |
| 6.673455.674376 | 922 | 6257 | intergenic | BWZt4 | BWZt9 | BWZt13 | BWZt21 | 1.00 | 0.00 |
| 6.782298.789507 | 7210 | 8041 | intergenic | BWZt4 | BWZt9 | BWZt13 | BWZt21 | 1.00 | 0.00 |
| 6.804863.807140 | 2278 | 7530 | intergenic | BWZt4 | BWZt9 | BWZt13 | BWZt21 | 1.00 | 0.00 |
| 6.927640.928434 | 795 | 4317 | intergenic | BWZt4 | BWZt9 | BWZt13 | BWZt21 | 1.00 | 0.00 |
| 6.999529.999949 | 421 | 6710 | intergenic | BWZt4 | BWZt9 | BWZt13 | BWZt21 | 1.00 | 0.00 |
| 7.1165974.1167274 | 1301 | 4533 | gene | BWZt4 | | BWZt13 | BWZt21 | 1.00 | 0.00 |
| 7.1263898.1264763 | 866 | 7224 | intergenic | BWZt4 | BWZt9 | BWZt13 | BWZt21 | 1.00 | 0.00 |
| 7.127872.128627 | 756 | 7253 | intergenic | BWZt4 | BWZt9 | BWZt13 | BWZt21 | 1.00 | 0.00 |
| 7.1351801.1352313 | 513 | 5993 | intergenic | BWZt4 | BWZt9 | BWZt13 | BWZt21 | 1.00 | 0.00 |
| 7.1677669.1677729 | 61 | 13600 | rRNA_gene | BWZt4 | BWZt9 | | BWZt21 | 1.00 | 0.00 |
| 7.1678104.1678175 | 72 | 5575 | rRNA_gene | | | BWZt13 | BWZt21 | 1.00 | 0.00 |
| 7.1685939.1685999 | 61 | 13600 | rRNA_gene | BWZt4 | BWZt9 | | BWZt21 | 1.00 | 0.00 |
| 7.1686374.1686445 | 72 | 5575 | rRNA_gene | | | BWZt13 | BWZt21 | 1.00 | 0.00 |
| 7.179855.180653 | 799 | 5162 | intergenic | | BWZt9 | BWZt13 | BWZt21 | 1.00 | 0.00 |
| 7.1804941.1806212 | 1272 | 7862 | intergenic | BWZt4 | BWZt9 | BWZt13 | BWZt21 | 1.00 | 0.00 |
| 7.1814849.1815870 | 1022 | 7794 | intergenic | BWZt4 | BWZt9 | BWZt13 | BWZt21 | 1.00 | 0.00 |
| 7.1870917.1873373 | 2457 | 2167 | gene | | BWZt9 | BWZt13 | BWZt21 | 1.00 | 0.00 |
| 7.2249688.2249714 | 27 | 908 | intergenic | | BWZt9 | BWZt13 | BWZt21 | 1.00 | 0.00 |
| 7.2536483.2537653 | 1171 | 8978 | intergenic | BWZt4 | BWZt9 | BWZt13 | BWZt21 | 1.00 | 0.00 |
| 7.2613424.2626681 | 13258 | 13000 | intergenic | | BWZt9 | BWZt13 | BWZt21 | 1.00 | 0.00 |
| 7.2645542.2651117 | 5576 | 990 | intergenic | | BWZt9 | BWZt13 | BWZt21 | 1.00 | 0.00 |
| 7.299254.300443 | 1190 | 1324 | gene | | | | BWZt21 | 1.00 | 0.05 |
| 7.452408.453395 | 988 | 8466 | intergenic | BWZt4 | BWZt9 | BWZt13 | BWZt21 | 1.00 | 0.00 |
| 7.582594.583749 | 1156 | 6210 | intergenic | BWZt4 | BWZt9 | BWZt13 | BWZt21 | 1.00 | 0.00 |
| 7.584106.585040 | 935 | 7723 | intergenic | BWZt4 | BWZt9 | BWZt13 | BWZt21 | 1.00 | 0.00 |
| 7.594685.595531 | 847 | 3772 | intergenic | BWZt4 | BWZt9 | BWZt13 | BWZt21 | 1.00 | 0.00 |
| 7.64956.65152 | 197 | 651000 | rRNA_gene | BWZt4 | BWZt9 | BWZt13 | BWZt21 | 1.00 | 0.00 |
| 7.755129.758077 | 2949 | 7841 | intergenic | BWZt4 | BWZt9 | BWZt13 | BWZt21 | 1.00 | 0.00 |
| 7.826820.827289 | 470 | 248 | intergenic | | BWZt9 | | | 1.00 | 0.04 |
| 7.883863.884771 | 909 | 1091 | intergenic | | | BWZt13 | | 1.00 | 0.00 |
| 7.927599.928416 | 818 | 6923 | intergenic | BWZt4 | BWZt9 | BWZt13 | BWZt21 | 1.00 | 0.00 |
| 7.95108.95160 | 53 | 7758 | intergenic | | BWZt9 | BWZt13 | BWZt21 | 1.00 | 0.00 |
| 7.951736.951950 | 215 | 854 | intergenic | | | BWZt13 | BWZt21 | 1.00 | 0.00 |
| 8.1105306.1107648 | 2343 | 541 | gene | | | | BWZt21 | 1.00 | 0.00 |
| 8.1152834.1153178 | 345 | 3101 | tRNA_gene | | | BWZt13 | | 1.00 | 0.00 |
| 8.135097.135842 | 746 | 7312 | intergenic | BWZt4 | BWZt9 | BWZt13 | BWZt21 | 1.00 | 0.00 |
| 8.1375496.1375926 | 431 | 4085 | intergenic | BWZt4 | BWZt9 | | BWZt21 | 1.00 | 0.00 |
| 8.1401272.1402008 | 737 | 1823 | intergenic | | | BWZt13 | BWZt21 | 1.00 | 0.00 |
| 8.1402168.1402594 | 427 | 17300 | intergenic | BWZt4 | BWZt9 | BWZt13 | BWZt21 | 1.00 | 0.00 |
| 8.1402809.1403198 | 390 | 16800 | intergenic | BWZt4 | BWZt9 | BWZt13 | BWZt21 | 1.00 | 0.00 |
| 8.1449944.1450478 | 535 | 703 | pseudogenic_tRNA | BWZt4 | | | | 1.00 | 0.04 |
| 8.1496933.1498574 | 1642 | 1560 | intergenic | | BWZt9 | BWZt13 | BWZt21 | 1.00 | 0.00 |
| 8.1619634.1620112 | 479 | 5534 | pseudogenic_tRNA | | | BWZt13 | | 1.00 | 0.00 |
| 8.1778342.1779282 | 941 | 6727 | intergenic | BWZt4 | BWZt9 | BWZt13 | BWZt21 | 1.00 | 0.00 |
| 8.1779347.1780499 | 1153 | 7781 | intergenic | BWZt4 | BWZt9 | BWZt13 | BWZt21 | 1.00 | 0.00 |
| 8.1816353.1818241 | 1889 | 4718 | intergenic | BWZt4 | BWZt9 | BWZt13 | BWZt21 | 1.00 | 0.00 |
| 8.1826676.1828336 | 1661 | 6470 | intergenic | BWZt4 | BWZt9 | BWZt13 | BWZt21 | 1.00 | 0.00 |
| 8.1839987.1840797 | 811 | 6638 | intergenic | BWZt4 | BWZt9 | BWZt13 | BWZt21 | 1.00 | 0.00 |
| 8.192706.195020 | 2315 | 528 | gene | | | BWZt13 | | 1.00 | 0.04 |
| 8.1960420.1962344 | 1925 | 814 | gene | | | | BWZt21 | 1.00 | 0.02 |
| 8.1964648.1966838 | 2191 | 7782 | intergenic | BWZt4 | BWZt9 | BWZt13 | BWZt21 | 1.00 | 0.00 |

| | | | | | | | | | |
|---|---|---|---|---|---|---|---|---|---|
| 8.2000570.2000759 | 190 | 648000 | intergenic | BWZt4 | BWZt9 | BWZt13 | BWZt21 | 1.00 | 0.00 |
| 8.2011199.2012128 | 930 | 4458 | intergenic | BWZt4 | BWZt9 | BWZt13 | BWZt21 | 1.00 | 0.00 |
| 8.2142675.2143478 | 804 | 5490 | intergenic | BWZt4 | BWZt9 | BWZt13 | BWZt21 | 1.00 | 0.00 |
| 8.2168811.2169041 | 231 | 15300 | pseudogenic_tRNA | | BWZt9 | | BWZt21 | 1.00 | 0.00 |
| 8.2202983.2203732 | 750 | 6510 | intergenic | BWZt4 | BWZt9 | BWZt13 | BWZt21 | 1.00 | 0.00 |
| 8.2292152.2298576 | 6425 | 7366 | intergenic | BWZt4 | BWZt9 | BWZt13 | BWZt21 | 1.00 | 0.00 |
| 8.2378470.2380599 | 2130 | 1708 | intergenic | | BWZt9 | BWZt13 | BWZt21 | 1.00 | 0.00 |
| 8.2403753.2409329 | 5577 | 5495 | intergenic | | BWZt9 | BWZt13 | BWZt21 | 1.00 | 0.00 |
| 8.2418353.2420254 | 1902 | 324 | intergenic | | | | BWZt21 | 1.00 | 0.03 |
| 8.2428790.2431187 | 2398 | 443 | gene | BWZt4 | | | | 1.00 | 0.03 |
| 8.2431279.2439751 | 8473 | 5831 | intergenic | | BWZt9 | BWZt13 | BWZt21 | 1.00 | 0.00 |
| 8.420489.421629 | 1141 | 7561 | intergenic | BWZt4 | BWZt9 | BWZt13 | BWZt21 | 1.00 | 0.00 |
| 8.558236.559373 | 1138 | 7380 | intergenic | BWZt4 | BWZt9 | BWZt13 | BWZt21 | 1.00 | 0.00 |
| 8.564251.565700 | 1450 | 7114 | intergenic | BWZt4 | BWZt9 | BWZt13 | BWZt21 | 1.00 | 0.00 |
| 8.78306.79126 | 821 | 143 | gene | | | | BWZt21 | 1.00 | 0.03 |
| 8.821716.821919 | 204 | 919 | intergenic | | | | BWZt21 | 1.00 | 0.01 |
| 8.828807.828884 | 78 | 1147 | intergenic | | | | BWZt21 | 1.00 | 0.00 |
| 8.829026.829103 | 78 | 1210 | intergenic | | | | BWZt21 | 1.00 | 0.00 |
| 9.1327176.1327455 | 280 | 1079 | gene | | | BWZt13 | | 1.00 | 0.00 |
| 9.1413289.1414068 | 780 | 5207 | intergenic | BWZt4 | BWZt9 | BWZt13 | BWZt21 | 1.00 | 0.00 |
| 9.1417709.1418662 | 954 | 7248 | intergenic | BWZt4 | BWZt9 | BWZt13 | BWZt21 | 1.00 | 0.00 |
| 9.1426458.1427268 | 811 | 8397 | intergenic | BWZt4 | BWZt9 | BWZt13 | BWZt21 | 1.00 | 0.00 |
| 9.1433239.1434246 | 1008 | 8398 | intergenic | BWZt4 | BWZt9 | BWZt13 | BWZt21 | 1.00 | 0.00 |
| 9.166474.167593 | 1120 | 427 | gene | | | BWZt13 | | 1.00 | 0.02 |
| 9.170649.171598 | 950 | 1023 | gene | | | BWZt13 | | 1.00 | 0.02 |
| 9.1755672.1756026 | 355 | 486 | intergenic | | BWZt9 | | | 1.00 | 0.01 |
| 9.1792703.1793023 | 321 | 535 | intergenic | | BWZt9 | | | 1.00 | 0.03 |
| 9.1904100.1904911 | 812 | 20600 | gene | BWZt4 | BWZt9 | | | 1.00 | 0.00 |
| 9.1963691.1964465 | 775 | 5928 | intergenic | BWZt4 | BWZt9 | BWZt13 | BWZt21 | 1.00 | 0.00 |
| 9.1998601.1999461 | 861 | 8548 | intergenic | BWZt4 | BWZt9 | BWZt13 | BWZt21 | 1.00 | 0.00 |
| 9.2.19666 | 19665 | 32500 | intergenic | BWZt4 | BWZt9 | BWZt13 | BWZt21 | 0.97 | 0.00 |
| 9.2127241.2137629 | 10389 | 8893 | intergenic | | BWZt9 | BWZt13 | BWZt21 | 1.00 | 0.00 |
| 9.282960.283771 | 812 | 7060 | intergenic | BWZt4 | BWZt9 | BWZt13 | BWZt21 | 1.00 | 0.00 |
| 9.293777.294104 | 328 | 2067 | intergenic | | | BWZt13 | BWZt21 | 1.00 | 0.00 |
| 9.403302.404358 | 1057 | 6079 | intergenic | BWZt4 | BWZt9 | BWZt13 | BWZt21 | 1.00 | 0.00 |
| 9.406820.407637 | 818 | 5652 | intergenic | BWZt4 | BWZt9 | BWZt13 | BWZt21 | 1.00 | 0.00 |
| 9.64052.64887 | 836 | 8422 | intergenic | BWZt4 | BWZt9 | BWZt13 | BWZt21 | 1.00 | 0.00 |
| 9.66682.67503 | 822 | 6768 | intergenic | | BWZt9 | BWZt13 | BWZt21 | 1.00 | 0.00 |
| 9.972252.973061 | 810 | 6699 | intergenic | BWZt4 | BWZt9 | BWZt13 | BWZt21 | 1.00 | 0.00 |
| 10.1059761.1061128 | 1368 | 308 | intergenic | | | BWZt13 | BWZt21 | 1.00 | 0.02 |
| 10.1407605.1408991 | 1387 | 7568 | intergenic | BWZt4 | BWZt9 | BWZt13 | BWZt21 | 1.00 | 0.00 |
| 10.160001.161905 | 1905 | 284 | intergenic | | | | BWZt21 | 1.00 | 0.03 |
| 10.1638022.1644097 | 6076 | 5301 | intergenic | | BWZt9 | BWZt13 | BWZt21 | 1.00 | 0.00 |
| 10.229566.233698 | 4133 | 1942 | intergenic | | | BWZt13 | | 1.00 | 0.00 |
| 10.445195.445392 | 198 | 651000 | intergenic | BWZt4 | BWZt9 | BWZt13 | BWZt21 | 1.00 | 0.00 |
| 10.531312.533737 | 2426 | 1011 | intergenic | BWZt4 | | | | 1.00 | 0.00 |
| 10.57964.59013 | 1050 | 164 | intergenic | | | | BWZt21 | 1.00 | 0.03 |
| 10.797801.797951 | 151 | 7899 | intergenic | BWZt4 | | | | 1.00 | 0.00 |
| 11.1119359.1119964 | 606 | 1513 | intergenic | BWZt4 | BWZt9 | | BWZt21 | 1.00 | 0.00 |
| 11.1235102.1235419 | 318 | 1889 | intergenic | BWZt4 | BWZt9 | BWZt13 | BWZt21 | 1.00 | 0.00 |
| 11.1593332.1594253 | 922 | 5471 | intergenic | BWZt4 | BWZt9 | BWZt13 | BWZt21 | 1.00 | 0.00 |
| 11.1597670.1598473 | 804 | 5719 | intergenic | BWZt4 | BWZt9 | BWZt13 | BWZt21 | 1.00 | 0.00 |
| 11.1615364.1617483 | 2120 | 672 | intergenic | | BWZt9 | BWZt13 | BWZt21 | 1.00 | 0.00 |
| 11.32325.33623 | 1299 | 7625 | intergenic | BWZt4 | BWZt9 | BWZt13 | BWZt21 | 1.00 | 0.00 |
| 11.36845.38682 | 1838 | 6518 | intergenic | BWZt4 | BWZt9 | BWZt13 | BWZt21 | 1.00 | 0.00 |
| 11.760814.761682 | 869 | 6828 | intergenic | BWZt4 | BWZt9 | BWZt13 | BWZt21 | 1.00 | 0.00 |
| 11.768223.768327 | 105 | 1930 | intergenic | | BWZt9 | BWZt13 | BWZt21 | 1.00 | 0.00 |
| 11.984387.985163 | 777 | 5718 | intergenic | BWZt4 | BWZt9 | BWZt13 | BWZt21 | 1.00 | 0.00 |
| 12.1012309.1012492 | 184 | 4578 | intergenic | | BWZt9 | BWZt13 | BWZt21 | 1.00 | 0.00 |
| 12.106740.107675 | 936 | 6795 | intergenic | BWZt4 | BWZt9 | BWZt13 | BWZt21 | 1.00 | 0.00 |
| 12.1121157.1124000 | 2844 | 1213 | intergenic | | | BWZt13 | BWZt21 | 1.00 | 0.00 |
| 12.13820.18839 | 5020 | 2711 | intergenic | | BWZt9 | BWZt13 | BWZt21 | 1.00 | 0.00 |
| 12.1402101.1405654 | 3554 | 303 | intergenic | | | | BWZt21 | 1.00 | 0.02 |
| 12.1427223.1442420 | 15198 | 12800 | intergenic | | BWZt9 | BWZt13 | BWZt21 | 1.00 | 0.00 |
| 12.1442499.1445629 | 3131 | 2138 | intergenic | | BWZt9 | BWZt13 | BWZt21 | 1.00 | 0.00 |
| 12.462748.462907 | 160 | 7669 | intergenic | | BWZt9 | BWZt13 | BWZt21 | 1.00 | 0.00 |
| 12.475.5013 | 4539 | 2488 | intergenic | | BWZt9 | BWZt13 | BWZt21 | 1.00 | 0.00 |
| 12.494199.496373 | 2175 | 629 | intergenic | | | | BWZt21 | 1.00 | 0.04 |
| 12.607359.608250 | 892 | 8532 | intergenic | BWZt4 | BWZt9 | BWZt13 | BWZt21 | 1.00 | 0.00 |
| 12.6522.12487 | 5966 | 2621 | intergenic | | BWZt9 | BWZt13 | BWZt21 | 1.00 | 0.00 |
| 12.681007.681905 | 899 | 7690 | intergenic | BWZt4 | BWZt9 | BWZt13 | BWZt21 | 1.00 | 0.00 |
| 12.732520.732860 | 341 | 7555 | intergenic | BWZt4 | BWZt9 | BWZt13 | BWZt21 | 1.00 | 0.00 |
| 12.778427.778715 | 289 | 26900 | intergenic | BWZt4 | BWZt9 | BWZt13 | BWZt21 | 1.00 | 0.00 |
| 12.832633.835493 | 2861 | 498 | intergenic | | | | BWZt21 | 1.00 | 0.02 |
| 12.870648.871567 | 920 | 4231 | intergenic | | | BWZt13 | BWZt21 | 1.00 | 0.00 |
| 13.1030685.1031551 | 867 | 6198 | intergenic | BWZt4 | BWZt9 | BWZt13 | BWZt21 | 1.00 | 0.00 |
| 13.1163358.1164368 | 1011 | 7628 | intergenic | BWZt4 | BWZt9 | BWZt13 | BWZt21 | 1.00 | 0.00 |
| 13.1168422.1177165 | 8744 | 4812 | intergenic | | BWZt9 | BWZt13 | BWZt21 | 1.00 | 0.00 |

| | | | | | | | | | |
|---|---|---|---|---|---|---|---|---|---|
| 13.1177242.1180173 | 2932 | 556 | intergenic | | BWZt9 | BWZt13 | BWZt21 | 1.00 | 0.00 |
| 13.1180236.1184672 | 4437 | 3083 | intergenic | BWZt4 | BWZt9 | BWZt13 | BWZt21 | 1.00 | 0.00 |
| 13.12950.15733 | 2784 | 343 | intergenic | | | | BWZt21 | 1.00 | 0.03 |
| 13.17800.20460 | 2661 | 581 | intergenic | | | BWZt13 | BWZt21 | 1.00 | 0.00 |
| 13.183698.187962 | 4265 | 18100 | intergenic | BWZt4 | BWZt9 | BWZt13 | BWZt21 | 1.00 | 0.00 |
| 13.320524.320760 | 237 | 1620 | intergenic | | | BWZt13 | BWZt21 | 1.00 | 0.03 |
| 13.3888.6772 | 2885 | 615 | intergenic | | | BWZt13 | BWZt21 | 1.00 | 0.01 |
| 13.442993.443802 | 810 | 2419 | intergenic | | BWZt9 | BWZt13 | | 1.00 | 0.01 |
| 13.479338.479478 | 141 | 651000 | intergenic | BWZt4 | BWZt9 | BWZt13 | BWZt21 | 1.00 | 0.00 |
| 13.645590.645871 | 282 | 651000 | intergenic | BWZt4 | BWZt9 | BWZt13 | BWZt21 | 1.00 | 0.00 |
| 13.6864.9270 | 2407 | 580 | intergenic | BWZt4 | | | | 1.00 | 0.03 |
| 13.775785.776744 | 960 | 7477 | intergenic | BWZt4 | BWZt9 | BWZt13 | BWZt21 | 1.00 | 0.00 |
| 13.846733.850190 | 3458 | 738 | intergenic | | | BWZt13 | BWZt21 | 1.00 | 0.01 |
| 13.910888.911312 | 425 | 6533 | intergenic | BWZt4 | BWZt9 | BWZt13 | BWZt21 | 1.00 | 0.00 |
| 13.93164.93950 | 787 | 7338 | intergenic | BWZt4 | BWZt9 | BWZt13 | BWZt21 | 1.00 | 0.00 |
| 13.951679.951897 | 219 | 960 | intergenic | | BWZt9 | BWZt13 | BWZt21 | 1.00 | 0.00 |
| 14.1.5659 | 5659 | 2757 | intergenic | | BWZt9 | BWZt13 | | 1.00 | 0.00 |
| 14.280035.280949 | 915 | 8400 | intergenic | BWZt4 | BWZt9 | BWZt13 | BWZt21 | 1.00 | 0.00 |
| 14.294633.295658 | 1026 | 6317 | intergenic | BWZt4 | BWZt9 | BWZt13 | BWZt21 | 1.00 | 0.00 |
| 14.32035.35785 | 3751 | 2495 | intergenic | | BWZt9 | BWZt13 | BWZt21 | 1.00 | 0.00 |
| 14.532521.533209 | 689 | 6133 | intergenic | BWZt4 | BWZt9 | BWZt13 | BWZt21 | 1.00 | 0.00 |
| 14.546974.547880 | 907 | 5036 | intergenic | BWZt4 | BWZt9 | BWZt13 | BWZt21 | 1.00 | 0.00 |
| 14.5736.29448 | 23713 | 32900 | intergenic | BWZt4 | BWZt9 | BWZt13 | BWZt21 | 1.00 | 0.00 |
| 14.632345.633222 | 878 | 7820 | intergenic | BWZt4 | BWZt9 | BWZt13 | BWZt21 | 1.00 | 0.00 |
| 14.644522.646581 | 2060 | 456 | intergenic | | | | BWZt21 | 1.00 | 0.02 |
| 14.712306.713157 | 852 | 7576 | intergenic | BWZt4 | BWZt9 | BWZt13 | BWZt21 | 1.00 | 0.00 |
| 14.744382.745201 | 820 | 7678 | intergenic | BWZt4 | BWZt9 | BWZt13 | BWZt21 | 1.00 | 0.00 |
| 14.767975.769865 | 1891 | 348 | intergenic | | BWZt9 | BWZt13 | BWZt21 | 1.00 | 0.05 |
| 14.769927.771285 | 1359 | 6122 | intergenic | | BWZt9 | BWZt13 | BWZt21 | 1.00 | 0.00 |
| 15.14826.24490 | 9665 | 9097 | intergenic | BWZt4 | BWZt9 | BWZt13 | BWZt21 | 1.00 | 0.00 |
| 15.166892.167796 | 905 | 7609 | intergenic | BWZt4 | BWZt9 | BWZt13 | BWZt21 | 1.00 | 0.00 |
| 15.2.14750 | 14749 | 19300 | intergenic | BWZt4 | BWZt9 | BWZt13 | BWZt21 | 1.00 | 0.00 |
| 15.25780.26891 | 1112 | 3419 | intergenic | | BWZt9 | BWZt13 | BWZt21 | 1.00 | 0.00 |
| 15.325288.325837 | 550 | 1175 | intergenic | | | BWZt13 | BWZt21 | 1.00 | 0.00 |
| 15.44255.45101 | 847 | 7099 | intergenic | BWZt4 | BWZt9 | BWZt13 | BWZt21 | 1.00 | 0.00 |
| 15.51821.52932 | 1112 | 8984 | intergenic | BWZt4 | BWZt9 | BWZt13 | BWZt21 | 1.00 | 0.00 |
| 15.583955.585999 | 2045 | 1528 | intergenic | | BWZt9 | BWZt13 | BWZt21 | 1.00 | 0.00 |
| 15.589581.597368 | 7788 | 5681 | intergenic | | BWZt9 | BWZt13 | BWZt21 | 1.00 | 0.00 |
| 15.597420.601006 | 3587 | 2454 | intergenic | | BWZt9 | BWZt13 | BWZt21 | 1.00 | 0.00 |
| 15.601708.639500 | 37793 | 44900 | intergenic | BWZt4 | BWZt9 | BWZt13 | BWZt21 | 0.97 | 0.00 |
| 16.2.25177 | 25176 | 29200 | intergenic | BWZt4 | BWZt9 | BWZt13 | BWZt21 | 1.00 | 0.00 |
| 16.26979.31598 | 4620 | 1045 | intergenic | | BWZt9 | BWZt13 | BWZt21 | 1.00 | 0.00 |
| 16.36395.40088 | 3694 | 1016 | intergenic | | BWZt9 | BWZt13 | BWZt21 | 1.00 | 0.00 |
| 16.412864.413679 | 816 | 8179 | intergenic | BWZt4 | BWZt9 | BWZt13 | BWZt21 | 1.00 | 0.00 |
| 16.420030.421422 | 1393 | 4996 | intergenic | BWZt4 | BWZt9 | BWZt13 | BWZt21 | 1.00 | 0.00 |
| 16.429283.430197 | 915 | 5759 | intergenic | | BWZt9 | BWZt13 | BWZt21 | 1.00 | 0.00 |
| 16.578572.583821 | 5250 | 4864 | intergenic | | BWZt9 | BWZt13 | BWZt21 | 1.00 | 0.00 |
| 16.597962.607044 | 9083 | 11000 | intergenic | BWZt4 | BWZt9 | BWZt13 | BWZt21 | 1.00 | 0.00 |
| 16.84151.85006 | 856 | 6833 | intergenic | | BWZt9 | BWZt13 | BWZt21 | 1.00 | 0.00 |
| 16.89218.90204 | 987 | 7976 | intergenic | BWZt4 | BWZt9 | BWZt13 | BWZt21 | 1.00 | 0.00 |
| 17.20145.23364 | 3220 | 12500 | intergenic | BWZt4 | BWZt9 | BWZt13 | BWZt21 | 1.00 | 0.00 |
| 17.23479.32114 | 8636 | 7584 | intergenic | | BWZt9 | BWZt13 | BWZt21 | 1.00 | 0.00 |
| 17.32194.70869 | 38676 | 34800 | intergenic | BWZt4 | BWZt9 | BWZt13 | BWZt21 | 1.00 | 0.00 |
| 17.561480.566789 | 5310 | 4601 | intergenic | | BWZt9 | BWZt13 | BWZt21 | 1.00 | 0.00 |
| 17.566841.583652 | 16812 | 17100 | intergenic | BWZt4 | BWZt9 | BWZt13 | BWZt21 | 1.00 | 0.00 |
| 17.79759.81052 | 1294 | 6345 | intergenic | BWZt4 | BWZt9 | BWZt13 | BWZt21 | 1.00 | 0.00 |
| 18.112235.113035 | 801 | 6393 | intergenic | BWZt4 | BWZt9 | BWZt13 | BWZt21 | 1.00 | 0.00 |
| 18.237222.238251 | 1030 | 8647 | intergenic | BWZt4 | BWZt9 | BWZt13 | BWZt21 | 1.00 | 0.00 |
| 18.287412.292428 | 5017 | 783 | intergenic | | | BWZt13 | BWZt21 | 1.00 | 0.01 |
| 18.431645.432912 | 1268 | 7542 | intergenic | BWZt4 | BWZt9 | BWZt13 | BWZt21 | 1.00 | 0.00 |
| 18.536203.544521 | 8319 | 5487 | intergenic | | BWZt9 | BWZt13 | BWZt21 | 1.00 | 0.00 |
| 18.544573.545584 | 1012 | 4816 | intergenic | | BWZt9 | BWZt13 | BWZt21 | 1.00 | 0.00 |
| 18.8453.9307 | 855 | 7409 | intergenic | BWZt4 | BWZt9 | BWZt13 | BWZt21 | 1.00 | 0.00 |
| 18.85060.85835 | 776 | 7423 | intergenic | BWZt4 | BWZt9 | BWZt13 | BWZt21 | 1.00 | 0.00 |
| 18.94463.94734 | 272 | 4169 | intergenic | BWZt4 | BWZt9 | BWZt13 | BWZt21 | 1.00 | 0.00 |
| 19.104217.104960 | 744 | 7287 | intergenic | BWZt4 | BWZt9 | BWZt13 | BWZt21 | 1.00 | 0.00 |
| 19.2210.3581 | 1372 | 645 | intergenic | | BWZt9 | BWZt13 | BWZt21 | 1.00 | 0.01 |
| 19.3645.8939 | 5295 | 9526 | intergenic | BWZt4 | BWZt9 | BWZt13 | BWZt21 | 1.00 | 0.00 |
| 19.527513.527919 | 407 | 6989 | intergenic | BWZt4 | BWZt9 | BWZt13 | BWZt21 | 1.00 | 0.00 |
| 20.1166.4761 | 3596 | 2395 | intergenic | | BWZt9 | BWZt13 | BWZt21 | 1.00 | 0.00 |
| 20.230607.231363 | 757 | 6902 | intergenic | BWZt4 | BWZt9 | BWZt13 | BWZt21 | 1.00 | 0.00 |
| 20.297635.298058 | 424 | 620 | intergenic | | | | BWZt21 | 1.00 | 0.02 |
| 20.453037.466416 | 13380 | 13900 | intergenic | | BWZt9 | BWZt13 | BWZt21 | 1.00 | 0.00 |
| 20.467436.471636 | 4201 | 2520 | intergenic | | BWZt9 | BWZt13 | BWZt21 | 1.00 | 0.00 |
| 20.5781.9528 | 3748 | 1156 | intergenic | | BWZt9 | BWZt13 | BWZt21 | 1.00 | 0.00 |
| 21.2.26958 | 26957 | 32800 | intergenic | BWZt4 | BWZt9 | BWZt13 | BWZt21 | 0.99 | 0.00 |
| 21.227542.233177 | 5636 | 5514 | intergenic | | BWZt9 | BWZt13 | BWZt21 | 1.00 | 0.00 |
| 21.27372.40407 | 13036 | 13000 | intergenic | | BWZt9 | BWZt13 | BWZt21 | 1.00 | 0.00 |

| | | | | | | | | | |
|---|---|---|---|---|---|---|---|---|---|
| 21.349300.352594 | 3295 | 1062 | intergenic | | | BWZt13 | BWZt21 | 1.00 | 0.00 |
| 21.400171.401658 | 1488 | 959 | intergenic | BWZt4 | BWZt9 | BWZt13 | BWZt21 | 1.00 | 0.00 |
| 21.401722.409211 | 7490 | 3487 | intergenic | | BWZt9 | BWZt13 | BWZt21 | 1.00 | 0.00 |
| 21.40468.43135 | 2668 | 2069 | intergenic | | BWZt9 | BWZt13 | BWZt21 | 1.00 | 0.00 |
| 21.43194.45416 | 2223 | 3362 | intergenic | | BWZt9 | BWZt13 | BWZt21 | 1.00 | 0.00 |

† The difference of the maximum posterior probability between the fungus containing samples (infected wheat and in vitro culture) and the fungus free samples (mock-inoculated healthy wheat)

‡ Minimum sRNA loci differential expression false discovery rate (FDR) in at least one comparison between the infected wheat samples and the corresponding mock samples

**Table S4.** *Zymoseptoria tritici* IPO323 sRNA loci and mature sRNAs predicted to target wheat transcripts.

**A. sRNA loci generating mature sRNAs, which have predicted mRNAs targets in wheat**

1.168753-169514
1.2-24760
1.24832-33907
1.2630102-2631752
1.3107284-3109571
1.3404853-3405015
1.4420671-4420786
1.4519579-4520495
1.4859784-4862055
1.5208279-5209283
1.52723-55652
1.55724-59340
1.5574650-5575522
1.59399-69532
1.6022523-6024517
1.6024577-6027831
1.6042189-6044283
1.6044579-6052106
1.6052501-6055814
1.6060199-6062484
1.6062562-6077812
1.6077865-6084551
1.610884-611019
1.681033-681226
1.912518-913393
1.952268-954112
2.1061972-1062929
2.1295279-1297239
2.1333569-1334006
2.1741314-1742939
2.1895220-1896101
2.1903263-1903455
2.1909278-1910404
2.236665-237674
2.3187737-3188483
2.3189040-3189775
2.3276658-3277710
2.3411312-3412084
2.341988-345005
2.3632877-3633814
2.3801864-3816890
2.3816969-3821391
2.3839620-3844042
2.400134-400399
2.47729-49709
2.532319-532572
2.819512-820307
3.1178-7084
3.1848836-1849733
3.2252943-2253304
3.2375162-2376724

3.2736368-2736981
3.2779587-2780080
3.3241153-3242334
3.3403412-3404238
3.387886-388363
3.391037-392727
3.401660-407274
3.587789-590578
3.7151-18710
3.822679-823459
4.1251113-1252024
4.170472-171758
4.2228316-2228727
4.234947-235725
4.2644748-2645586
4.272845-274005
4.2866934-2868658
4.2869853-2880010
5.1045197-1045805
5.1092560-1092727
5.181519-182321
5.1875780-1875912
5.2318983-2319321
5.2686132-2687400
5.2806545-2846948
5.2850297-2861802
5.514262-514929
5.523622-524805
5.557661-560726
5.59060-61544
5.748438-748459
5.918436-919203
5.934505-936848
6.1267883-1270184
6.16617-39543
6.173879-174893
6.2022916-2027069
6.2108904-2109094
6.2414949-2415730
6.39611-43211
6.597638-598424
6.673455-674376
6.782298-789507
6.804863-807140
6.927640-928434
6.999529-999949
7.1263898-1264763
7.127872-128627
7.1351801-1352313
7.179855-180653
7.1804941-1806212
7.1814849-1815870
7.2536483-2537653
7.2613424-2626681
7.452408-453395

7.582594-583749
7.584106-585040
7.594685-595531
7.755129-758077
7.951736-951950
8.135097-135842
8.1402168-1402594
8.1402809-1403198
8.1496933-1498574
8.1778342-1779282
8.1779347-1780499
8.1816353-1818241
8.1826676-1828336
8.1839987-1840797
8.1964648-1966838
8.2000570-2000759
8.2011199-2012128
8.2142675-2143478
8.2202983-2203732
8.2292152-2298576
8.2378470-2380599
8.2403753-2409329
8.2431279-2439751
8.420489-421629
8.558236-559373
8.564251-565700
9.1413289-1414068
9.1417709-1418662
9.1426458-1427268
9.1433239-1434246
9.170649-171598
9.1904100-1904911
9.1963691-1964465
9.1998601-1999461
9.2-19666
9.2127241-2137629
9.282960-283771
9.403302-404358
9.406820-407637
9.64052-64887
9.66682-67503
9.972252-973061
10.1407605-1408991
10.1638022-1644097
10.445195-445392
10.797801-797951
11.1119359-1119964
11.1593332-1594253
11.1597670-1598473
11.1615364-1617483
11.32325-33623
11.36845-38682
11.760814-761682
11.768223-768327
11.984387-985163

12.1012309-1012492
12.106740-107675
12.1427223-1442420
12.1442499-1445629
12.607359-608250
12.6522-12487
12.681007-681905
13.1030685-1031551
13.1163358-1164368
13.1168422-1177165
13.1180236-1184672
13.183698-187962
13.479338-479478
13.645590-645871
13.775785-776744
13.910888-911312
13.93164-93950
13.951679-951897
14.1-5659
14.280035-280949
14.294633-295658
14.32035-35785
14.532521-533209
14.5736-29448
14.632345-633222
14.712306-713157
14.744382-745201
14.769927-771285
15.14826-24490
15.166892-167796
15.2-14750
15.25780-26891
15.325288-325837
15.44255-45101
15.51821-52932
15.583955-585999
15.589581-597368
15.597420-601006
15.601708-639500
16.2-25177
16.26979-31598
16.412864-413679
16.420030-421422
16.429283-430197
16.578572-583821
16.597962-607044
16.84151-85006
16.89218-90204
17.20145-23364
17.23479-32114
17.32194-70869
17.561480-566789
17.566841-583652
17.79759-81052
18.112235-113035

18.237222-238251
18.431645-432912
18.536203-544521
18.544573-545584
18.8453-9307
18.85060-85835
19.104217-104960
19.3645-8939
19.527513-527919
20.230607-231363
20.297635-298058
20.453037-466416
21.2-26958
21.227542-233177
21.27372-40407
21.401722-409211
21.40468-43135
21.43194-45416

**B. Mature sRNAs predicted to target wheat mRNAs**

| Mature sRNA ID | sRNA sequence |
|---|---|
| ZtsRNA1 | TTGGGGAATCCGTAGTGTTGT |
| ZtsRNA1.1 | GGGGAATCCGTGGTGTTGTATG |
| ZtsRNA2 | TGCACTGGTTGCTCGAACGCT |
| ZtsRNA3 | TAACCATCTTTCGGGTCTGACT |
| ZtsRNA3.1 | AACCATCTTTCGGGTCTGACT |
| ZtsRNA3.2 | GTTAACCATCTTTCGGGTCTGACT |
| ZtsRNA4 | TGGAGATCGCGAAGGAGGTTC |
| ZtsRNA4.1 | TCCATGGAGATCGCGAAGGAG |
| ZtsRNA5.1 | TGGCTCAGCTGCTGGCTCTGG |
| ZtsRNA5.2 | TGGCTCGGCTGCTGGCTCAGC |
| ZtsRNA6.1 | TAGGATTGCCTCTATAGCCTG |
| ZtsRNA6.2 | TGGGATTGCCTCTATAGCCTG |
| ZtsRNA7.1 | TATGTCTCGGGCTCGTCGATA |
| ZtsRNA7.2 | TGTGTCTCGGGCTCGTCGATA |
| ZtsRNA8.1 | TTCGCGCTTCCTCTTAACCAT |
| ZtsRNA8.2 | TTCGCGCTTCCTCTTAACTAT |
| ZtsRNA9.1 | TAGTCGTCTCGTCCCAGAACT |
| ZtsRNA9.2 | TAGTCGTCTCGTCCTAGAACT |
| ZtsRNA10.1 | TAGTATCGTCCTTCCGTATCG |
| ZtsRNA10.2 | TAGTATCGTCCTTCCGTGTCG |
| ZtsRNA11.1 | TATCTGTTTAACATCTTCATC |
| ZtsRNA11.2 | TATCTGTTTAACATCTTTATC |
| ZtsRNA12 | AACACCTGTACGTGGTGCCGGCTA |
| ZtsRNA13.1 | CAGGGAATTTTTGGCATACCA |
| ZtsRNA13.2 | AGGGAATTTTTGGCATACCAC |
| ZtsRNA14.1 | CACCAACGATGTTCTCTTTTT |
| ZtsRNA14.2 | TCACCAACGATGTTCTCTTTT |
| ZtsRNA15.1 | TGACAGCATCGTCGACAATGG |
| ZtsRNA15.2 | TTGACAGCATCGTCGACAATG |
| ZtsRNA16.1 | AAATGATTTAATGAGCCTGACT |
| ZtsRNA16.2 | AATGATTTAATGAGCCTGACT |

| | |
|---|---|
| ZtsRNA17.1 | TCCGTCGGCCTCAATCACACC |
| ZtsRNA17.2 | TTCCGTCGGCCTCAATCACAC |
| ZtsRNA18.1 | TATTGATGCCGGCGAATCGAT |
| ZtsRNA18.2 | TGATGCCGGCGAATCGATTGG |
| ZtsRNA19.1 | TACTTCGCTGCACGATCTCCC |
| ZtsRNA19.2 | TCGCTGCACGATCTCCCGGAG |
| ZtsRNA20.1 | TCGGTCCACAAAGCTCCAGCC |
| ZtsRNA20.2 | TGCATCGGTCCACAAAGCTCC |
| ZtsRNA21.1 | GGGCTTATGGTGCAGTGGTAGC |
| ZtsRNA21.2 | TTATGGTGCAGTGGTAGCATT |
| ZtsRNA22.1 | CTTTGGATCGGAGTTGTGGGATGG |
| ZtsRNA22.2 | TTTGGATCGGAGTTGTGGGATGGC |
| ZtsRNA23 | AATACACGTGGCGGCGGCGGT |
| ZtsRNA24.1 | CCATCTTTTTTTTTGTCTGATC |
| ZtsRNA24.2 | CTTACCATCTTTTTTTTTGTCTGA |
| ZtsRNA25 | AATCCGAATCCGACTCTGACGACG |
| ZtsRNA26 | CAATTCGTGCACCTTCTTTTT |
| ZtsRNA27 | CCCGCAAGATCAGTCCTCTCG |
| ZtsRNA28 | CGATGAGTAGGACGTAAAGTAG |
| ZtsRNA29 | CTTTCCACTCCGACGTCATGT |
| ZtsRNA30.1 | TATGGAGCGTGCGATTCTTTT |
| ZtsRNA30.2 | TTATGGAGCGTGCGATTCTTT |
| ZtsRNA30.3 | TTATGGAGCGTGCGATTCTTTT |
| ZtsRNA31.1 | TTACGACTTTTCTTCTTACCT |
| ZtsRNA31.2 | TTTACGACTTTTCTTCTTACC |
| ZtsRNA32.1 | TCGAATCCCATTCTGTTCACC |
| ZtsRNA32.2 | TCGAATCCCATTCTGTTCACCA |
| ZtsRNA32.3 | TTCGAATCCCATTCTGTTCACCA |
| ZtsRNA33.1 | CTTCGTCGAACGACCATGTTTT |
| ZtsRNA33.2 | CTTCGTCGAACGACCATGTTTTT |
| ZtsRNA33.3 | TCTTCGTCGAACGACCATGTT |
| ZtsRNA34.1 | CACATCAGATTCTGATTCTGGG |
| ZtsRNA34.2 | CATCACATCAGATTCTGATTC |
| ZtsRNA34.3 | TCATCACATCAGATTCTGATT |
| ZtsRNA35.1 | TCCGGCTCCTCTGATTCGTCG |
| ZtsRNA35.2 | TGTCCGGCTCCTCTGATTCGT |
| ZtsRNA36.1 | CGGAATTTTTGATGCCCCCCA |
| ZtsRNA36.2 | CGTGTTAGGCGGAATTTTTGATGC |
| ZtsRNA36.3 | GCGGAATTTTTGATGCCCCCC |
| ZtsRNA37.1 | TACCAATTCGAGCTGCTCTTT |
| ZtsRNA37.2 | TGCTCTTTGATCTGCTGATAG |
| ZtsRNA37.3 | TTCGAGCTGCTCTTTGATCTG |
| ZtsRNA38.1 | ACTGCCTCTGGCTGACGCGAA |
| ZtsRNA38.2 | CGCCAGTATTGCACTGCCTCTGGC |
| ZtsRNA38.3 | TCGCCAGTATTGCACTGCCTCTGG |
| ZtsRNA39.1 | CCTTTGGACTGCTCTACGGAGC |
| ZtsRNA39.2 | CCTTTGGACTGCTCTACGGAGCC |
| ZtsRNA39.3 | CTTTGGACTGCTCTACGGAGC |
| ZtsRNA39.4 | TTTGGACTGCTCTACGGAGCC |
| ZtsRNA40.1 | ATGGCACTTTCGAAGACTCGG |
| ZtsRNA40.2 | ATGGCACTTTCGAAGACTCGGC |

| | |
|---|---|
| ZtsRNA40.3 | ATGGCACTTTCGAAGACTCGGCG |
| ZtsRNA40.4 | ATGGCACTTTCGAAGACTCGGCGA |
| ZtsRNA41.1 | TGGGCGTTGGGCGTTGGTCAG |
| ZtsRNA41.2 | TGGGCGTTGGTCAGGGTCACC |
| ZtsRNA41.3 | TTGGGCGTTGGGCGTTGGTCA |
| ZtsRNA41.4 | TTGGGCGTTGGTCAGGGTCAC |
| ZtsRNA42.1 | AGAAGGGAAACCAAGACAGTCT |
| ZtsRNA42.2 | GAAGGGAAACCAAGACAGTCT |
| ZtsRNA43.1 | TCAGGCAAAGGCTGACGACGA |
| ZtsRNA43.2 | TTCAGGCAAAGGCTGACGACG |
| ZtsRNA43.3 | TTCAGGCAAAGGCTGACGACGA |
| ZtsRNA44.1 | TATCGTCGTCAGCCTTTGCCT |
| ZtsRNA44.2 | TATCGTCGTCAGCCTTTGCCTG |
| ZtsRNA45.1 | CCGTCCGATCAGCCATGATAA |
| ZtsRNA45.2 | CTTCCCGTCCGATCAGCCATG |
| ZtsRNA45.3 | CTTCCCGTCCGATCAGCCATGA |
| ZtsRNA45.4 | CTTCCCGTCCGATCAGCCATT |
| ZtsRNA45.5 | TTCCCGTCCGATCAGCCATGA |
| ZtsRNA46.1 | TAGGACTGCTATTTTCCTAGG |
| ZtsRNA46.2 | TATTTTCCTAGGACTGCTATC |
| ZtsRNA46.3 | TCTTCCTAGGACTGCTATCTT |
| ZtsRNA46.4 | TGCTATTTTCCTAGGACTGCT |
| ZtsRNA46.5 | TGCTATTTTCCTAGGACTGCTA |
| ZtsRNA47.1 | TATCTTCACGGGCGATGGTCG |
| ZtsRNA47.2 | TGGCGACGATATCTTCACGGG |
| ZtsRNA48.1 | AACGACGACGACGACTCTGAA |
| ZtsRNA48.2 | CAACGACGACGACGACTCTGAA |
| ZtsRNA49.1 | TACCTCGCCGTCTCAAATATC |
| ZtsRNA49.2 | TTTTGCTCTACCTCGCCGTCT |
| ZtsRNA50.1 | ACACTAGTACTCAAGTTGGTG |
| ZtsRNA50.2 | CTAGTACTCAAGTTGGTGACA |
| ZtsRNA51.1 | AAGTTGGTGACAATTGGGGAAT |
| ZtsRNA51.2 | AGTTGGTGACAATTGGGGAAT |
| ZtsRNA51.3 | TCAAGTTGGTGACAATTGGGG |
| ZtsRNA51.4 | TGACAATTGGGGAATCCGTAG |
| ZtsRNA51.5 | TGGTGACAATTGGGGAATCCG |
| ZtsRNA52.1 | TTTGAGACGGCGAGGTAGAGC |
| ZtsRNA52.2 | TTTGAGACGGCGAGGTAGAAC |
| ZtsRNA52.3 | TGAGACGGCGAGGTAGAGCAA |
| ZtsRNA53 | CTTTGACCTCTTCGGAGGCGTGAA |
| ZtsRNA54 | GCACCTTCACGCGGTGGCCGTCGT |
| ZtsRNA55 | GCCCAATCTTCCTTGGATGTCTGA |
| ZtsRNA56 | GGACCCTTGGCGCAGCGGTAGCGT |
| ZtsRNA57 | GGGTAGTTGGTCTAGGGGTAT |
| ZtsRNA58 | GTGAGTATTACTCTAGTTCCGTTA |
| ZtsRNA59 | TAACCACTCGGCCACCTGTCC |
| ZtsRNA60 | TAACGATTTCTTTTAAGCTCT |
| ZtsRNA61 | TAACGCGGGCCGCACCTTCCG |
| ZtsRNA62 | TAAGCCACATCTCCATTGTAG |
| ZtsRNA63 | TAAGGGACGTAGACGAGGTAG |
| ZtsRNA64 | TAATAGGATAGTATAGAAGGC |

| | |
|---|---|
| ZtsRNA65 | TACACTACGTCGTCGAGCTCC |
| ZtsRNA66 | TACACTGCGAACTGCTCTACC |
| ZtsRNA67 | TACACTGGTCCTTTCTTGAAG |
| ZtsRNA68 | TACATACGAGGTGTTCTGCTG |
| ZtsRNA69 | TACCTCTTGGATTTGCTGTCT |
| ZtsRNA70 | TACGGATCTTCCTTCTACGAG |
| ZtsRNA71 | TACTCATCGGACGGAGCCTTT |
| ZtsRNA72 | TACTTACTCGTCCTGCTCGTC |
| ZtsRNA73 | TACTTCCGGACAAAACGCCCT |
| ZtsRNA74 | TACTTTCAAACTCGATCAATT |
| ZtsRNA75 | TAGAAGCGATAGAGGAGGACT |
| ZtsRNA76 | TAGAAGGAATGTCCGTAGATG |
| ZtsRNA77 | TAGAATCTTTGAAATTAGCTA |
| ZtsRNA78 | TAGACGCGGAGCTAGAGCTCG |
| ZtsRNA79 | TAGACTCTGCTGACTAGGCAC |
| ZtsRNA80 | TAGAGCAAAACCGGTGGCTAG |
| ZtsRNA81 | TAGAGGACGAGGAGGACGAGG |
| ZtsRNA82 | TAGAGGACGATTTCGACGACG |
| ZtsRNA83 | TAGAGGTAGAGCTAGTAGACA |
| ZtsRNA84 | TAGCAGCTCTCCGAGATCTCC |
| ZtsRNA85 | TAGCATTGGGCGAGAAATCCG |
| ZtsRNA86 | TAGCCACCGGTTTTGCTCTAC |
| ZtsRNA87 | TAGCCGGCCTGCCCTAGCATC |
| ZtsRNA88 | TAGCGACCGGACACGGACCCT |
| ZtsRNA89 | TAGCGCTCGAGGATCTAGATT |
| ZtsRNA90 | TAGCGGCAAGGTAGGCTAGAA |
| ZtsRNA91 | TAGCGTGCGTAGCTCTCTATT |
| ZtsRNA92 | TAGCGTTCTCGGGAGGATCTG |
| ZtsRNA93 | TAGCTCGAGCGGTAACAGACT |
| ZtsRNA94 | TAGCTCGTCCTGTTTTCCTAG |
| ZtsRNA95 | TAGGATGGGGTAATCGACCCG |
| ZtsRNA96 | TAGGCAGGAGACATCGAGACA |
| ZtsRNA97 | TAGGGACATGCGATTTCGATT |
| ZtsRNA98 | TAGGGCATTGCGAGTGAACAG |
| ZtsRNA99 | TAGGTAAGAAGAAAAGTCGTA |
| ZtsRNA100 | TAGGTATCGAACTCGATAATT |
| ZtsRNA101 | TAGGTCTCTAGAAGCACTTCG |
| ZtsRNA102 | TAGGTCTGGCCCTCCTCTAGC |
| ZtsRNA103 | TAGTATCCGTCGTCGAGCTCG |
| ZtsRNA104 | TAGTCGACGAAGTGCTTCCCG |
| ZtsRNA105 | TAGTCGCATCTGTCGTAATCG |
| ZtsRNA106 | TAGTTAGGGAAGTTCTGCTCT |
| ZtsRNA107 | TATCATGACTGCTGCGTTGCC |
| ZtsRNA108 | TATCCACTGTATACAACACTC |
| ZtsRNA109 | TATCGGATCTCGTGGAGTATG |
| ZtsRNA110 | TATCGGTAGTCTCTCTTTTTG |
| ZtsRNA111 | TATCGTCGATTCTACAAGCCG |
| ZtsRNA112 | TATCGTCGCCAACACCCTCGA |
| ZtsRNA113 | TATCTCTGCCGGAGCTGAATG |
| ZtsRNA114 | TATCTGACTTCACTTGACGAA |
| ZtsRNA115 | TATGCTCGACGAACCAGGTCT |

| | |
|---|---|
| ZtsRNA116 | TATGCTCGGTCGACACCATTC |
| ZtsRNA117 | TATGGCGTGTGACTCAAGCTC |
| ZtsRNA118 | TCAAAGGCGTCGGAGCTGCTG |
| ZtsRNA119 | TCAACCGATCGCTTCGTCTCC |
| ZtsRNA120 | TCAACGAGCGTGGATACTTCA |
| ZtsRNA121 | TCACTTCTGAACCTTGTGACG |
| ZtsRNA122 | TCAGAGTCGTCGTCAGAGTCG |
| ZtsRNA123 | TCAGATCAGGTGCAAAGGTAG |
| ZtsRNA124 | TCCGACGTCCCTCTAGTGCTT |
| ZtsRNA125 | TCCGGAGCGTGCTTCTTTACG |
| ZtsRNA126 | TCCGGATTCGGCTTCGCTATC |
| ZtsRNA127 | TCCTCCGTCTCGTCCGCCTCG |
| ZtsRNA128 | TCCTGTAAGCACTGGAGGGACG |
| ZtsRNA129 | TCGAAGGGATGCTCGTAGAAG |
| ZtsRNA130 | TCGACCGGTCTATTTCCCTCT |
| ZtsRNA131 | TCGACTATGTCACTGGCAGCG |
| ZtsRNA132 | TCGATTCCGGGGTCGACCTCA |
| ZtsRNA133 | TCGCATTCTCGCACCGTCGCA |
| ZtsRNA134 | TCGCCCCGAGCGGCTCTGACC |
| ZtsRNA135 | TCGCCCGTGAAGATATCGTCG |
| ZtsRNA136 | TCGCTTCGGCTCATACTCCTA |
| ZtsRNA137 | TCGGACCCTTTCCCTTTCGAG |
| ZtsRNA138 | TCGGATTCGGATTCAGAGTCG |
| ZtsRNA139 | TCGGCTAGAATTCCTCCTCTA |
| ZtsRNA140 | TCGGTAGGACGTTAAAAGGCT |
| ZtsRNA141 | TCGGTCGCGGTAGAGCTAGCG |
| ZtsRNA142 | TCGTTACTTCGCCGACCTCTC |
| ZtsRNA143 | TCTCATGTGTCCATGTTAGCT |
| ZtsRNA144 | TCTGAACCTCCTTCGCGATCC |
| ZtsRNA145 | TGAAGGGCCTTGTCGAATACA |
| ZtsRNA146 | TGACCAACGACACGAGCTGCT |
| ZtsRNA147 | TGACCAGGTCTAACTCCAACT |
| ZtsRNA148 | TGACGAAGTCTCTAGAGAAGG |
| ZtsRNA149 | TGAGACTTGTCGAGCGGGCGA |
| ZtsRNA150 | TGAGTCTGCTCTTCTTGTTAC |
| ZtsRNA151 | TGATTGTTCCAGAAGATGAGT |
| ZtsRNA152 | TGCACGTGTACCATTAGCCTG |
| ZtsRNA153 | TGCCCGAGCCGTCCTCTGTAG |
| ZtsRNA154 | TGCCGCTGGGCGGATTTCTCG |
| ZtsRNA155 | TGCGACTGCAAGAGCAATTCG |
| ZtsRNA156 | TGCGCTGCGACGTAGGAGACG |
| ZtsRNA157 | TGCGTGCGGCCTTGAGCTTCTGA |
| ZtsRNA158 | TGCTGGAGCTTGCAAACGAGT |
| ZtsRNA159 | TGGACTGCTTCTTTCTTCTTC |
| ZtsRNA160 | TGGCGATGGCGACTGATCTGA |
| ZtsRNA161 | TGGCGCACCGCTGAACTTCCC |
| ZtsRNA162 | TGGGAGACGTGGTAGCCATCG |
| ZtsRNA163 | TGGTCGAGGGTGTTGGCGACG |
| ZtsRNA164 | TGGTGTTTGGCTTCATCTGCG |
| ZtsRNA165 | TGGTTATTCGTCGTGCTGATG |
| ZtsRNA166 | TGTCCGGTCTTGTTCCTTAGG |

| | |
|---|---|
| ZtsRNA167 | TGTCCTCGTCGTCGTCAAATG |
| ZtsRNA168 | TGTCGTTGGTCACCGTCCAGT |
| ZtsRNA169 | TGTCTTGGTTTCCCTTCCGAG |
| ZtsRNA170 | TGTTCGCAACGTAGAGGGCTG |
| ZtsRNA171 | TGTTGATCGAGGGCTCATTCC |
| ZtsRNA172 | TGTTGGCGGCGATATCTCTAC |
| ZtsRNA173 | TGTTGTGGTGGATCAAAGGCG |
| ZtsRNA174 | TGTTTGATCCGGACTGTCTTG |
| ZtsRNA175 | TTAATATCTCCGAGAACGACA |
| ZtsRNA176 | TTACCCCATCCTAGCCACCGG |
| ZtsRNA177 | TTCCACCTCTTCATCATTCCG |
| ZtsRNA178 | TTCCAGATCCAGGTCGTGAACATC |
| ZtsRNA179 | TTCCGCGGCAGTACTATGTCT |
| ZtsRNA180 | TTCCTCTGCCGTCTCTCCTCG |
| ZtsRNA181 | TTCCTTTTCCGACTGTTCTCG |
| ZtsRNA182 | TTCGCCACCCTCTGAACGGTG |
| ZtsRNA183 | TTCGTGTGCTCGTATTCGTGA |
| ZtsRNA184 | TTGCGGTCGACGAATCAGAGG |
| ZtsRNA185 | TTGCTACGTCAGTCTGAGCTA |
| ZtsRNA186 | TTGGATTGAGGTCTGGTGAAT |
| ZtsRNA187 | TTGGCGACCACGATCTCCTCG |
| ZtsRNA188 | TTGTTGCTCTGATCAATGCAC |
| ZtsRNA189 | TTTCCAGCCCACCTTGCCGCT |
| ZtsRNA190 | TTTCGAGAAGGATTTCGGGCG |
| ZtsRNA191 | TTTCTGCTCTGCTAGCTTTCT |
| ZtsRNA192 | TTTGCACCTGATCTGAACTGA |
| ZtsRNA193 | TTTGCCGAGGACTCTCCTAAG |

**C. sRNA loci generating the four mature sRNAs (ZtsRNA1 - ZtsRNA4) selected for an in depth study**

| sRNA locus | Mature sRNA ID |
|---|---|
| 1.3404853-3405015 | ZtsRNA1 |
| 2.532319-532572 | ZtsRNA1 |
| 2.1903263-1903455 | ZtsRNA1 |
| 3.3241153-3242334 | ZtsRNA1 |
| 5.514262-514929 | ZtsRNA1 |
| 8.2000570-2000759 | ZtsRNA1 |
| 10.445195-445392 | ZtsRNA1 |
| 13.479338-479478 | ZtsRNA1 |
| 13.645590-645871 | ZtsRNA1 |
| 1.3107284-3109571 | ZtsRNA2 |
| 2.341988-345005 | ZtsRNA2 |
| 5.557661-560726 | ZtsRNA2 |
| 13.183698-187962 | ZtsRNA2 |
| 3.2375162-2376724 | ZtsRNA3 |
| 8.1402809-1403198 | ZtsRNA4 |

**Table S5.** Wheat transcripts predicted to be targeted by *Zymoseptoria tritici* IPO323 mature sRNAs.

| targeted wheat transcript | wheat transcript annotation |
|---|---|
| Traes_1AL_0F92F56F8.1 | mitochondrial-processing peptidase subunit, mitochondrial precursor, putative, expressed;0.0;723 |
| Traes_1AL_14F4067BA.1 | cytochrome b6, putative, expressed;4e-31;133 |
| Traes_1AL_1CB3A83EE.1 | IQ calmodulin-binding motif family protein, putative, expressed;0.0;843 |
| Traes_1AL_2B919E578.3 | crcB-like protein, expressed;0.0;680 |
| Traes_1AL_375367151.1 | fatty acid hydroxylase, putative, expressed;4e-144;508 |
| Traes_1AL_486E54C74.1 | expressed protein;0.0;761 |
| Traes_1AL_4A458DF14.1 | PPR repeat containing protein, expressed;0.0;844 |
| Traes_1AL_6CD69EE7E.1 | cobalt ion transporter, putative, expressed;9e-150;527 |
| Traes_1AL_99F5BB0D8.1 | methyltransferase, putative, expressed;0.0;1089 |
| Traes_1AL_A3B85CAE6.1 | tesmin/TSO1-like CXC domain containing protein, expressed;1e-09;60.8 |
| Traes_1AL_ABE93C8E7.1 | peptidyl-tRNA hydrolase, mitochondrial precursor protein, putative, expressed;3e-97;352 |
| Traes_1AL_B33A889C4.1 | endonuclease/exonuclease/phosphatase family domain containing protein, expressed;0.0;783 |
| Traes_1AL_C4650CF30.3 | selT-like protein precursor, putative, expressed;6e-65;243 |
| Traes_1AL_C4B40A214.1 | NAD dependent epimerase/dehydratase family protein, putative, expressed;0.0;737 |
| Traes_1AL_D1E192C9C.2 | leucine-rich repeat family protein, putative, expressed;0.0;711 |
| Traes_1AL_D74509506.2 | ternary complex factor MIP1, putative, expressed;0.0;757 |
| Traes_1AL_E7F4CC5A1.1 | zinc finger, C3HC4 type domain containing protein, expressed;6e-121;431 |
| Traes_1AL_EB5141D4C.7 | NBS-LRR disease resistance protein, putative, expressed;0.0;667 |
| Traes_1AS_AC509D7F5.1 | protein kinase APK1A, chloroplast precursor, putative, expressed;5e-171;598 |
| Traes_1AS_B3DDADFFA.1 | AGC_PVPK_like_kin82y.12 - ACG kinases include homologs to PKA, PKG and PKC, expressed;0.0;653 |
| Traes_1AS_D2C38EEF3.1 | nuclear factor related to kappa-B-binding protein, related, putative, expressed;0.0;1539 |
| Traes_1AS_EB5297361.1 | white-brown complex homolog protein 12, putative, expressed;0.0;996 |
| Traes_1BL_0EABC6DF5.1 | expressed protein;0.0;929 |
| Traes_1BL_190920E1E.1 | glucose-1-phosphate adenylyltransferase large subunit, chloroplast precursor, putative, expressed;0.0;860 |
| Traes_1BL_29B4E633D.1 | IQ calmodulin-binding motif family protein, putative, expressed;0.0;829 |
| Traes_1BL_39AE9ED1C.1 | SAM dependent carboxyl methyltransferase, putative, expressed;1e-111;400 |
| Traes_1BL_3EEA722DF.1 | ZOS5-12 - C2H2 zinc finger protein, expressed;6e-36;149 |
| Traes_1BL_42C7AFE7A.2 | OsFBX53 - F-box domain containing protein, expressed;3e-110;396 |
| Traes_1BL_44DACABDA.1 | expressed protein;0.0;634 |
| Traes_1BL_4B54A24F9.1 | n/a |
| Traes_1BL_6ABEEF6DC.1 | peptidyl-tRNA hydrolase, mitochondrial precursor protein, putative, expressed;8e-82;300 |
| Traes_1BL_7175F3833.1 | zinc finger, C3HC4 type domain containing protein, expressed;2e-129;460 |
| Traes_1BL_73EBB04B5.2 | glycosyl hydrolase, family 31, putative, expressed;0.0;795 |
| Traes_1BL_84EEAC5D5.4 | mitochondrial-processing peptidase subunit, mitochondrial precursor, putative, expressed;4e-133;472 |
| Traes_1BL_9046B5730.2 | cytoplasmic membrane protein, putative, expressed;5e-33;137 |
| Traes_1BL_93285025D.1 | lipoxygenase, putative, expressed;2e-85;315 |
| Traes_1BL_A0644DA53.1 | ternary complex factor MIP1, putative, expressed;4e-128;455 |
| Traes_1BL_B722F2F54.1 | Leucine Rich Repeat family protein, expressed;1e-177;620 |
| Traes_1BL_BEB9B5EF3.1 | myosin-2 heavy chain, non muscle, putative, expressed;0.0;749 |
| Traes_1BL_C943EF0D8.2 | selT-like protein precursor, putative, expressed;3e-65;244 |
| Traes_1BL_DF929E48F.1 | crcB-like protein, expressed;0.0;681 |
| Traes_1BL_E9106179C.2 | AGC_PVPK_like_kin82y.12 - ACG kinases include homologs to PKA, PKG and PKC, expressed;8e-63;239 |
| Traes_1BL_EABF29D66.1 | OsSub7 - Putative Subtilisin homologue, expressed;2e-67;252 |
| Traes_1BL_EC9A57F9E.1 | endonuclease/exonuclease/phosphatase family domain containing protein, expressed;0.0;780 |
| Traes_1BL_FDCE71D9E.1 | zinc finger, C3HC4 type domain containing protein, expressed;2e-73;274 |
| Traes_1BS_307AF1E47.1 | peroxidase precursor, putative, expressed;4e-64;242 |
| Traes_1BS_3D04F8758.1 | receptor-like protein kinase 2 precursor, putative, expressed;0.0;1704 |
| Traes_1BS_4A63D1DBC.2 | phosphatidate cytidylyltransferase, putative, expressed;0.0;700 |
| Traes_1BS_6799B4072.1 | cytochrome b6, putative, expressed;2e-25;114 |
| Traes_1BS_77D2B4C0D.2 | haloacid dehalogenase-like hydrolase family protein, putative, expressed;3e-165;578 |
| Traes_1BS_ACC0652C7.1 | transmembrane amino acid transporter protein, putative, expressed;0.0;785 |
| Traes_1BS_B64557385.1 | disease resistance protein RGA1, putative, expressed;0.0;691 |
| Traes_1BS_B734EDEA7.1 | metal cation transporter, putative, expressed;7e-132;468 |
| Traes_1BS_DB050544B.1 | KIP1, putative, expressed;9e-180;629 |
| Traes_1BS_FEE01077D.1 | expressed protein;3e-21;99.0 |
| Traes_1DL_14F4067BA.1 | cytochrome b6, putative, expressed;4e-31;133 |
| Traes_1DL_1F7D352CA.1 | NAD dependent epimerase/dehydratase family protein, putative, expressed;0.0;737 |
| Traes_1DL_23506ECCE.2 | IQ calmodulin-binding motif family protein, putative, expressed;0.0;843 |
| Traes_1DL_4518A1BF6.1 | peptidyl-tRNA hydrolase, mitochondrial precursor protein, putative, expressed;3e-86;315 |
| Traes_1DL_4CFBFAC6C.1 | flavonol synthase/flavanone 3-hydroxylase, putative, expressed;2e-53;206 |
| Traes_1DL_560607990.1 | DUF630/DUF632 domains containing protein, putative, expressed;0.0;744 |

**Table S6.** *Zymoseptoria tritici* (*Zt*) genes selected for gene silencing using BSMV-HIGS.

| **Target** | **Function** | **Expected phenotype** | **HIGS construct size* (in bp)** | **si-Fi software predictions (all siRNA hits *vs* effective siRNA hits)** | **si-Fi predicted off-target siRNA hits**** |
|---|---|---|---|---|---|
| *ZtCYP51* (construct M5) | Ergosterol biosynthesis | Lethal; fungicide target | 343 bp | 234 / 89 | Zt: no<br>Wheat: no |
| *ZtTUBa* (construct M4) | Microtubules | Lethal or reduced virulence | 281 bp | 193 / 68 | Zt: YES, to putative uncharacterised protein (siRNA hits - 61/22)<br>Wheat: no |
| *ZtTUBb* (construct M1) | Microtubules | Lethal or reduced virulence; fungicide target | 303 bp | 193 / 90 | Zt: no<br>Wheat: no |
| *ZtALG2* (construct M7) | Alpha-1,2-mannosyl-transferase | Loss of pathogenicity towards wheat | 315 bp | 209 / 86 | Zt: no<br>Wheat: no |

*size of protein coding sequences inserted into the BSMV vector (not including the LIC adaptor sequences)
**Tested for cross-hits against *Z. tritici* and wheat transcripts databases
